# Single-cell transcriptomics highlights sexual cues among reproductive life stages of uncultivated Acantharia (Radiolaria)

**DOI:** 10.1101/2024.07.02.601653

**Authors:** Iris Rizos, Sarah Romac, Caroline Juery, Charlotte Berthelier, Johan Decelle, Juliana Bernardes, Erwan Corre, Lucie Bittner, Fabrice Not

## Abstract

As an innate property of life, the ability to reproduce is a key process for the perpetuation of organisms. Along the evolution of protist reproductive strategies, the molecular machinery of sexual recombination is estimated to have been inherited from the last eukaryotic common ancestor (LECA). Nevertheless, unraveling the sexual cycles of extant free-living protist lineages remains challenging, given the enigmatic roles of many uncultivated life stages. Among the uncultivated planktonic group of Acantharia (Radiolaria), a hypothetical sexual cycle has been proposed since the late 19th century, including the existence of a gamete-like life stage of undetermined ploidy, referred to as swarmers. In order to investigate the sexual nature of acantharian reproductive stages, we conducted single-cell transcriptomic analysis across various acantharian life stages. Our results show distinct functional profiles for reproductive and vegetative life stages, while revealing the expression of the reference eukaryotic genes involved in gamete fusion, HAP2/GCS1 and GEX1-KAR5, in swarmers and pre-swarmer stages. Annotation of differentially expressed life stage-specific genes, also highlights putative meiosis-related functions among swarmers, while suggesting the existence of a potential swarmer/vegetative intermediate stage expressing putative growth-related genes. This original life stage-specific genetic data is coherent with morphological evidence supporting the existence of an acantharian sexual cycle, with swarmers acting as gametes. Moreover, it paves the way for a deeper understanding of radiolarian cell biology and ecology at a single-cell scale.

**Highlights:**

- Acantharia demonstrate both morphological and genetic evidence of a sexual cycle
- Acantharian reproductive stages are enriched in functions related to cell division
- Nuclear fusion gene family GEX1-KAR5 is up-regulated in putative acantharian gametes
- Most expressed genes specific to acantharian reproductive stages are unassigned
- Reproduction-specific unassigned genes include putative sex-related functions

## Introduction

Perpetuation by reproduction is a common and essential feature of all living beings on Earth (Fusco and Minelli, 2019). Among prokaryotes, binary fission is the dominant mode of reproduction, while eukaryotic microorganisms (protists) can present diverse sexual cycle strategies during their reproduction (Rizos et al., 2024 in press; Krueger-Hadfield, 2024). Up to now, the description of protist asexual (i.e., replicative) and sexual (i.e., recombinative) processes remain fairly limited and incomplete (Margulis and Sagan, 1986; Lahr et al., 2011). A recent phylogeny (Richter et al., 2022), has estimated that at least half of morphologically described free-living protists have an uncharacterised sexual cycle (Rizos et al., 2024 in press). Protist sexual cycles can be complex and involve multiple life stages (Margulis and Sagan, 1986; Grell, 1973; Lee et al., 2000). For instance, despite cyst life stages being frequently associated with resting stages, there are many cases where cysts have been reported to play a role in protist sexual cycles (e.g., diatoms and dinoflagellates (Archibald et al., 2017; Montagnes et al., 2011). Even among well studied, cultivable pathogenic protists, such as trypanosomes and plasmodiums, novel sexual life stages have been recently identified by functional analysis of genetic data (Howick et al., 2019; Howick et al., 2021).

Sexual recombination is an ancestral eukaryotic trait (Goodenough and Heitman, 2014; Speijer, 2016), whose genes have spread across the eukaryotic tree of life from a common genetic basis (Speijer et al., 2015). The genetic toolkit underlying sexual recombination is composed of genes conserved across many eukaryotic lineages and that are essential for two basic sexual cycle processes, i.e. meiosis and gamete fusion or syngamy (Hofstatter and Lahr, 2019). For instance, the meiosis specific gene SPO11, which encodes for an enzyme creating double-strand breaks (DSBs) in homologous chromosomes, and the gamete-specific gene HAP2/GCS1, responsible for gamete membrane fusion (plasmogamy), are both conserved across distant unicellular and multicellular lineages (Schurko and Logsdon, 2008); (Villeneuve and Hillers, 2001; Fedry et al., 2018). Among protists bearing unclear reproductive mechanisms, sexual genetic toolkit candidate genes have been successfully identified in genomic data of *Giardia* and *Trichomonas* (Metamonada, (Ramesh et al., 2005; (Malik et al., 2008) and multiple Amoebozoa (Hofstatter et al., 2018), but also in single-cell transcriptomes of *Cochlopodium* (Amoebozoa, (Tekle et al., 2020). Given the extensive diversity of protists with unobserved or hypothetical sexual cycles (Rizos et al., 2024 in press), genetic data constitute a powerful resource for inferring sexual recombination in protists with functionally ambiguous or uncultivated life stages.

Among uncultivated protists stand Acantharia, amoeboid-shaped planktonic cells bearing a mineral skeleton of sulfate strontium (SrSO4) (Suzuki and Not, 2015; ^A^nderson, 1983^)^. They inhabit the ocean, from surface waters to bathypelagic zones across all latitudes and are mostly abundant in photic layers, where they have established a successful symbiosis with photosynthetic microalgae, allowing them to thrive in oligotrophic conditions (Decelle and Not, 2015; Decelle et al., 2015). Acantharia are representatives of the group Radiolaria, the sister lineage of Foraminifera, and are part of the diverse TSAR clade (i.e., Telonemia-Stramenopiles-Alveolata-Rhizaria) (Burki et al., 2020). Radiolaria lack genomic references and only few transcriptomic data are available for a limited number of species (Suzuki and Not, 2015; Richter et al., 2022). Knowledge of their life cycle is restrained to field observations that have been going on since the H.M.S. Challenger’s expedition in the latest 19th century (Anderson, 1983; Haeckel, 1887). For Acantharia, the detailed descriptions and schemes of Schewiakoff (1926), Haeckel (1887) and other scientists (Hollande and Enjumet, 1957; Hollande et al., 1965; Febvre, 1977), have converged to a hypothetical life cycle that includes a gamete-like life stage of unknown ploidy, named “swarmers” (Suzuki and Not, 2015). In the hypothetical acantharian sexual cycle, swarmers are produced either via the vegetative cell or by a cyst (i.e., also referred to as “litholophus” stage for some Acantharia) (Febvre and Febvre, 1994; Decelle et al., 2013). The acantharian swarmers are morphologically similar to foraminiferan gametes. The sexual cycle of benthic Foraminifera has been described by the alternation between haploid and diploid life phases observed in established cultures (Darling et al., 2023; Lehmann et al., 2006). During the diploid life phase, the diploid foraminiferan known as agamont, undergoes meiosis, giving rise to haploid juveniles referred to as gamonts. The gamonts grow during the haploid phase and produce gametes. The gametes produced by the foraminiferan gamont have been confirmed to fuse and develop into diploid agamont juveniles (Darling et al., 2023; Lehmann et al., 2006). Nevertheless, unlike Foraminifera, the processes of swarmer fusion and zygote development have never been observed for Acantharia (Röttger, 1974; Anderson, 1983; Suzuki and Not, 2015).

Observing acantharian reproductive life stages is challenging since the triggers initiating radiolarian reproduction in the environment remain underscribed. While correlations of reproductive periods with lunar cycles have been identified for planktonic Foraminifera (Hohenegger et al., 2019), and many protist sexual cycles can be experimentally triggered by modifying their nutrient source (Rizos et al., 2024 in press), the short maintenance periods hinder the determination of stress factors that induce radiolarian swarmer production. Here, we have generated single-cell transcriptomes of cyst and swarmer life stages obtained during the maintenance of various Acantharia collected in the field. We comparatively analyze the gene expression of these functionally uncharacterised life stages and investigate their potential implication in a sexual cycle. With this approach, we provide genetic evidence for the acantharian hypothetical sexual cycle and highlight up-regulated protein groups with putative key functions for each life stage. Due to the abundance of Acantharia (Decelle and Not, 2015), their sexual stages significantly influence biogeochemical cycles through the creation of strontium gradients during cyst formation (Decelle et al., 2013). Thus, elucidating the sexual cycle of Acantharia represents a significant step forward in the comprehension of marine protist ecology.

## Results

### 1. Acantharian vegetative and reproductive life stages show different functional profiles

In the acantharian life cycle, nine different life stages (LS) can be distinguished according to morphological variations of the cells (Fig. 1A). The vegetative life stage (LS 1) can enter the process of swarmer formation (LS 2) following two distinct cellular pathways, depending on its symbiotic nature. To date, Acantharia hosting unicellular photosynthetic symbionts (i.e., photosymbionts, Fig. 1A - LS 1b) have been observed to produce swarmers by maintaining their amoeboid shape and undergoing cytoplasm compartmentalization (Fig. 1A - LS 2b). At this stage, the cytoplasm develops an homogeneous pink/brown granular appearance, accompanied by a subtle intracellular motion observable at high magnification (20-40X). Progressively, the cytoplasm compartments become smaller until they detach from the vegetative cell, giving the impression that the cell is dissolving (Fig. 1A - LS 3b) (Video S1). Two types of bi-flagellated swimming cells (i.e., swarmers) have been observed to ultimately form, produced by distinct vegetative cells (i.e., the two swarmer types have not been observed to emerge from the same cell). The two types of swarmers can be distinguished based on their size, morphology and movement. Small swarmers (Fig. 1A - LS 4) had an oval/conical shape of 2-3 um and appeared to be swimming rapidly in a brownian motion (Photo S1, Video S1). On the contrary, big swarmers (Fig. 1A - LS 5) appeared oval/round with a size twice larger (estimated ∼5-10 um) and moved in a slow rotating motion (Photo S1, Video S2). Neither fusion nor further development has been observed for any of the swarmer types. Big swarmers have been witnessed to emerge solely from a symbiotic vegetative Acantharia. Non-symbiotic vegetative Acantharia (Fig. 1A - LS 1a) either form swarmers directly from the amoeboid vegetative cell (Fig. 1A - pathway b) or via the formation of a cyst (Fig. 1A - pathway a), a pathway characterized by drastic morphological changes. During cyst formation the cellular membrane of the vegetative cell expands until all the spicules are internalized and then progressively degraded. A thick envelope is formed (of various shapes, i.e., here pear shaped but can be round or diamond shaped), while the integrity of the cell is maintained by few central spicules (Fig. 1A - LS 2a; Photo S2). The cellular wall becomes opaque and dense, ultimately forming a cyst (Fig. 1A - LS 2a’). Swarmer emergence from cysts occurs either from an opercule (Fig. 2A - A4ViCY, lighter edge on the right) or from holes at the periphery (Fig. 1A - LS 3a; Photo S1). Only small swarmers have been observed to emerge from cysts. Overall, the transition from the swarmer to the juvenile life stage (Fig. 1A - LS 6) remains hypothetical, as well as the molecular modifications underlying the morphological changes of the acantharian cell during its life cycle. The most prevalent hypothesis suggests that meiosis occurs during cytoplasm compartmentalisation and inside the cyst, while juveniles result from syngamy. Thus, we analyzed the transcriptomic profiles of 10 acantharian cells across different life stages: the vegetative life stage (Fig. 1A - LS 1a,b; Fig. 1B - samples A6GrVE, A7GrVE, A8GrVE, A9GrVE), the granular amoeba stage (Fig. 1A - LS 2b; Fig. 1B - sample A5ViME), the cyst stage (Fig. 1A - LS 2a,a’; Fig. 1B; Photo S2 - samples A2ViCY and A4ViCY, ongoing cyst formation and fully formed cyst respectively) and swarmer life stages (Fig. 1A - LS 4,5; Fig 1B; Photo S1; Video S1,S2 - samples A1ViSW, A2ViSW, A3ViSW) (cf. Materials & Methods, Table S1). The sampled cells correspond to three different acantharian clades (Fig. 1B). All swarmer samples belong to symbiotic acantharian Clade F, as the vegetative cells A6GrVE and A8GrVE. The cysts (i.e., A2ViCY and A4ViCY) and vegetative cell A9GrVE correspond to the non-symbiotic Clade C, while the granular amoeba stage (i.e., A5ViME) and the vegetative cell A7GrVE belong to the non-symbiotic Clade D. We studied life stages corresponding to various acantharian phylogenetic clades, in order to highlight common interspecies functional cues relative to their role in the life cycle. As the process of swarmer formation is not yet confirmed to be part of a sexual cycle, we will henceforth refer to the pre-swarmer (i.e., granular amoeba and cyst) and swarmer stages as reproductive.

**Fig. 1.**
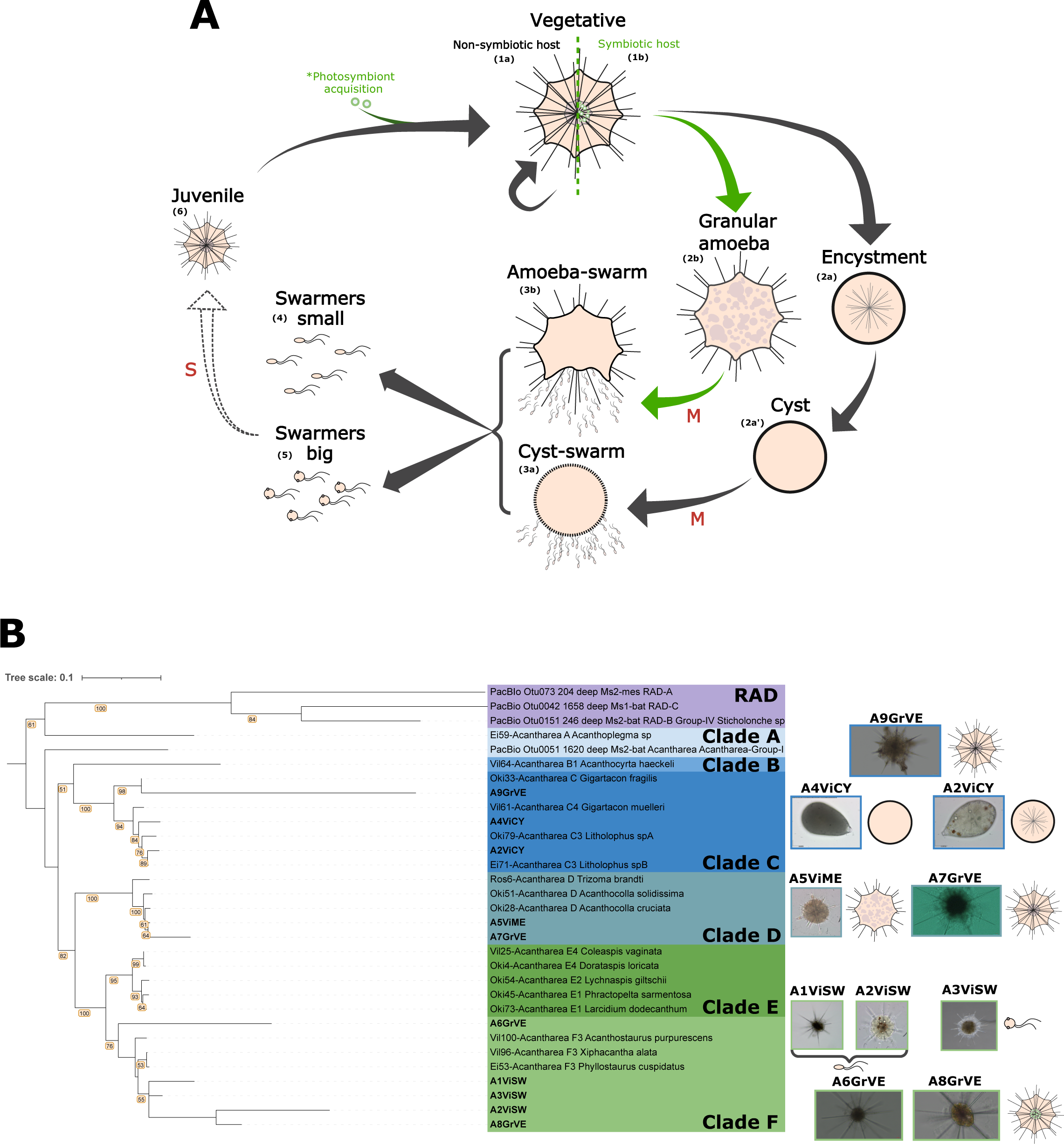
Placement of the sampled life stages in the acantharian hypothetical life cycle and phylogeny. (A) Hypothetical acantharian life cycle. Arrows illustrate life cycle transitions and red annotations indicate hypothetical processes, i.e. M=meiosis and S=syngamy. The life cycle is generic for all Acantharia and the developmental pathway followed by species harboring photosymbionts is represented in green. Supportive microscopy images of each life stage can be found on Fig. S?. (B) Acantharian 18S-28S rRNA maximum likelihood phylogeny. Branches support are indicated in orange boxes for bootstrap values >= 50. The scale bars are 100 um for A1_Vi_SW, 20 um for A2_Vi_SW, A5_Vi_ME, A4_Vi_CY and A2_Vi_CY and 14.594 um for A6GrVE, A7GrVE, A8GrVE and A9GrVE. The binocular magnification 2.5x for the image of A3_Vi_SW.

**Fig. 2.**
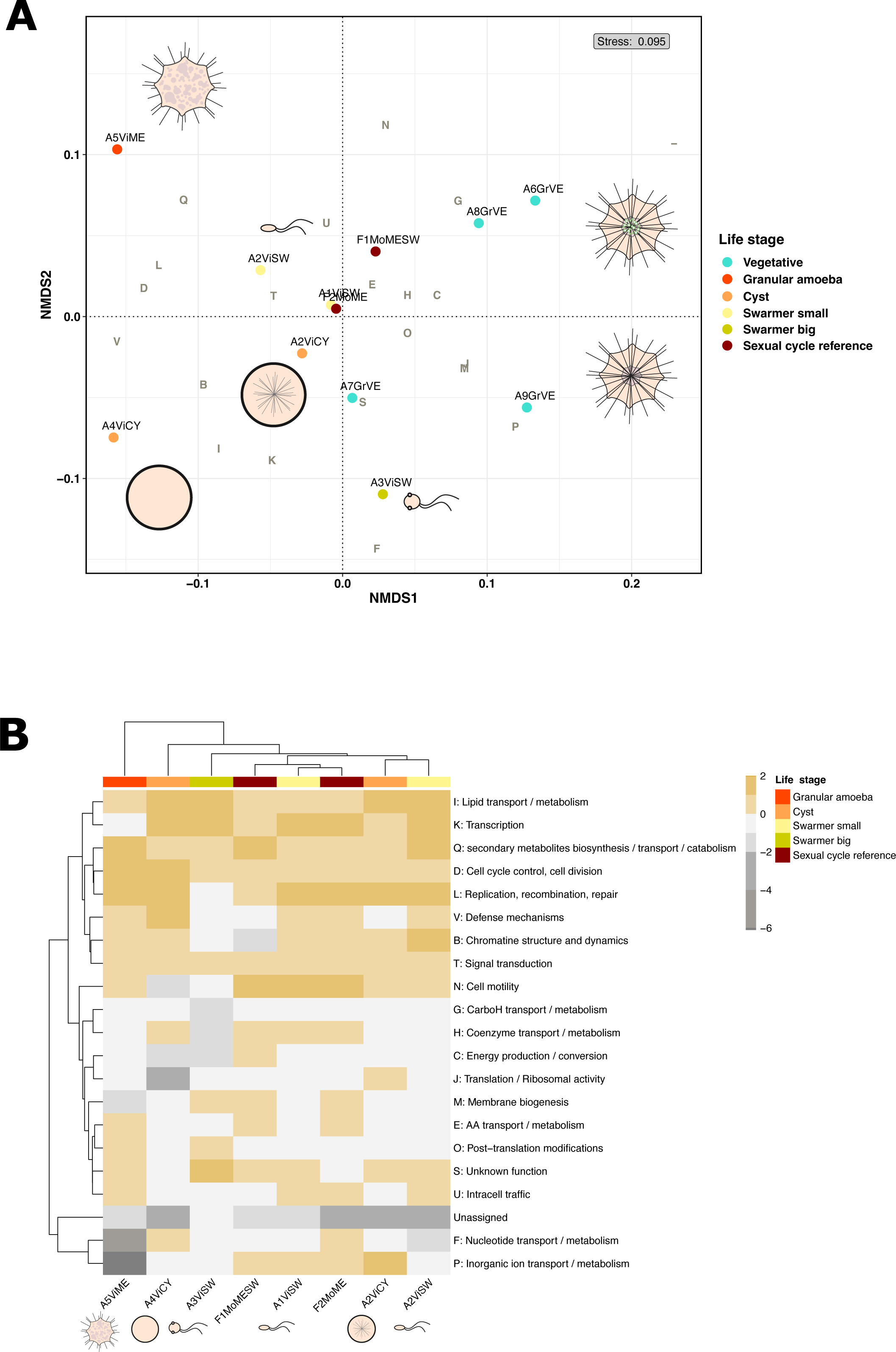
Functional description of acantharian life stages across the life cycle. (A) Dimensionality reduction analysis (non-metric multidimensional scaling, NMDS) of acantharian single-cells transcriptomes corresponding to different life stages (cf. colored codes) based on their functional expression in Clusters of Orthologous Groups (i.e., COGs). The letters correspond to different functional gene categories, the detail of which is available on the following part of the Figure. (B) Differential functional expression between reproductive stages of Radiolaria and Foraminifera in comparison with vegetative life stages. The scale bar represents the log2 fold-change of normalised COG expression.

In order to investigate the role of acantharian reproductive life stages, we functionally annotated predicted proteins of each life stage in Clusters of Orthologous Groups (COGs). As we hypothesize that acantharian reproductive stages are sexual, we integrated in our analysis two sexual life stages of Foraminifera: one meiotic life stage F2MoME (i.e., equivalent to the granular amoeba stage of Acantharia in terms of intracellular morphology) and one life stage ongoing gamete release, F1MoMESW (that is thus an intermediate life stage between the meiotic and gamete stage) (Photo S3). In the multivariate analysis NMDS, (i.e., Non Metric Multidimensional Scaling, Fig. 2A, NMDS1, stress score < 0.1), COG expression appeared to differentiate vegetative and reproductive life stages based on the first axis (Fig. 2A, NMDS1). Major COGs driving this differentiation oppose functions associated to reproductive life stages: lipids (I), transcription (K), signal transduction (T), intracellular traffic (U), defense (V), secondary metabolites (Q), replication (L), chromatin (B) and cell cycle (D), with functions associated to vegetative life stages: unassigned proteins (-,S), carbohydrate transport (G), energy production (C), membrane biogenesis (M), translation (J) and inorganic transport (P). Small and big swarmers were clustered separately according to axis 2 (Fig. 2A, NMDS2), being driven either by intracellular traffic (U) and signal transduction (T) or nucleotide transport/metabolism (F) and inorganic ion transport/metabolism (P) respectively. Overall, big swarmers were located equally proximate to vegetative and reproductive life stages (Fig. 2A, NMDS1). Within the reproductive life stages, cysts seemed to form their own cluster on the bottom left part of the graph, based on NMDS1. This trend was driven by the expression of COGs related to lipids (I), transcription (K), chromatin (B) and defense (V), while the granular amoeba life stage was located opposite to the cyst cluster (Fig. 2A, NMDS2). The swarmer A1ViSW and foraminiferan sexual cycle reference (F2MoME) were not clearly differentiated from the other life stages. Among vegetative life stages, symbiotic (A6GrVE, A8GrVE) and non-symbiotic (A7GrVE, A9GrVE) Acantharia are differentiated based on the second dimension of the analysis (Fig. 2A, NMDS2), where symbiotic cells are driven by carbohydrate transport (G) and unassigned proteins (-), while non-symbiotic cells are driven by inorganic ion transport/metabolism (P) and another group of unassigned proteins (S).

In order to highlight key functions of reproductive life stages, we performed a differential expression analysis between vegetative and reproductive stages using vegetative transcriptomes as references. The COGs characterizing the functional profiles of the reproductive life stages in the NMDS (Fig. 2A), also showed to be up-regulated compared to vegetative life stages (Fig. 2B). The functional profiles of the sexual cycle reference samples (Foraminifera meiosis and swarmer life stages, F1MoMESW, F2MoME) were not clustered according to taxonomy but rather to their life stage, as they were placed along with acantharian reproductive stages and closer to the acantharian swarmer A1ViSW. The most divergent functional profiles appeared to be those of big swarmers (A3ViSW), the fully developed cyst (A4ViCY) and the granular amoeba (A5ViME) life stages (Fig. 2B). In all reproductive and sexual reference stages, the lipid metabolism (I), secondary metabolites (Q), cell cycle control (D) and signal transduction (T) functional categories were up-regulated, in contrast to the carbohydrate metabolism (G) and unassigned COGs that were down-regulated. No distinct expression patterns specific to cysts or swarmers were observed. All acantharian reproductive life stages showed a 2-fold higher expression of functional categories involved in replication, recombination and repair (L) compared to vegetative stages, apart from big swarmers (Fig. 2B). The functional profile of big swarmers differed from small swarmers also in the down-regulation of COGs related to motility (N), chromatin dynamics (B), intracellular traffic (U), defense (V) and up-regulation of membrane biogenesis (M) and post-translational modifications (O). The functional category related to amino acid transport/metabolism was up-regulated both in the granular amoeba life stage of Acantharia and meiosis of Foraminifera sexual references (A5ViME, F1MoMESW, F2MoME), while it was down-regulated in swarmers and cysts. The two acantharian cysts were not clustered together, as the functional profiles of the fully formed cyst A4ViCY showed down-regulated translation (J), cell motility (N), and inorganic ion transport (P) COGs, in contrast to up-regulated coenzyme and nucleotide transport/metabolism (H, F) COGs (Fig. 2B). In contrast, the functional profile of the cyst A2ViCY ongoing cyst formation appeared to be closer to the small swarmer A2ViSW, although COGs involved in inorganic ion transport (P) and translation / ribosomal activity (J) were up-regulated in the cyst A2ViCY compared to the swarmer A2ViSW.

### 2. Acantharian reproductive life stages are enriched in meiosis related and syngamy specific reference proteins

Out of the 35 meiosis-related query proteins selected based on bibliography (cf. Materials and Methods, Table S2), 9 10 were found to be up-regulated in acantharian reproductive life stages (Fig. 3A). Among them, 5 meiosis-related predicted proteins were recovered in the acantharian cyst, A2ViCY. The meiosis functions found up-regulated in the cyst (A2ViCY) are related to the beginning and the end of the meiotic process and were involved in sister chromatid cohesion (i.e. SMC domains, PDS5), cross-over and mismatch repair (i.e. RAD52, EXO domain 1). The granular amoeba putative meiotic life stage (A5ViME) was found to be highly enriched in only 1 cross-over/mismatch repair protein (i.e. PMS1). Other cross-over/mismatch repair functions were found among swarmers (i.e. RAD1, MSH domain 2) but not in vegetative life stages. The double-strand DNA break protein NSB1 was enriched among swarmers and vegetative predicted proteins, but not pre-swarmer reproductive stages. The rhizarian sequence of the meiosis structural protein MNS1 was found up-regulated in the cyst A2ViCY and the big swarmer A3ViSW predicted proteins. No meiosis-related function was found to be common among all acantharian reproductive stages (i.e., the swarmers, granular amoeba and cyst stages). Syngamy-related functions associated to plasmogamy, karyogamy and flagella were found enriched in swarmer predicted proteins (Fig. 3A). The gamete-specific eukaryotic fusogen HAP2-GCS1 was up-regulated in granular amoeba, cyst and big swarmers (A3ViSW) life stages, with an expression higher in pre-swarmer stages. Apart from HAP2-GCS1, the cyst predicted proteins appeared to be enriched in the ciliar/flagellar domain CFA20. The karyogamy-specific protein GEX1-KAR5 was found enriched exclusively in swarmer predicted proteins, with a 3-fold expression in swarmer A1ViSW (Fig. 3A). Overall, apart from cyst A4ViCY, for which no sexual cycle reference was found and which has the lowest gene expression among all life stages (Fig. S1), more meiosis and syngamy related reference functions were up-regulated in reproductive life stages compared to vegetative ones (Fig. 3A). Nevertheless, in reproductive life stages, the predicted proteins with the highest expression were proteins either without annotation or corresponding to COGs (cf. Fig. 3B). In terms of relative number, the proportion of non-annotated predicted proteins was the highest among all transcriptomes, varying from ∼60% among vegetative and swarmer life stages, up to ∼90% among the granular amoeba and cyst life stages (Fig. 3B; Fig. S2). Thus, sexual cycle reference functions represented a minor proportion (<1%) of up-regulated predicted proteins in acantharian reproductive life stages, while the most expressed predicted proteins lack precise functional annotation (Fig. 3B; Fig. S2).

**Fig. 3.**
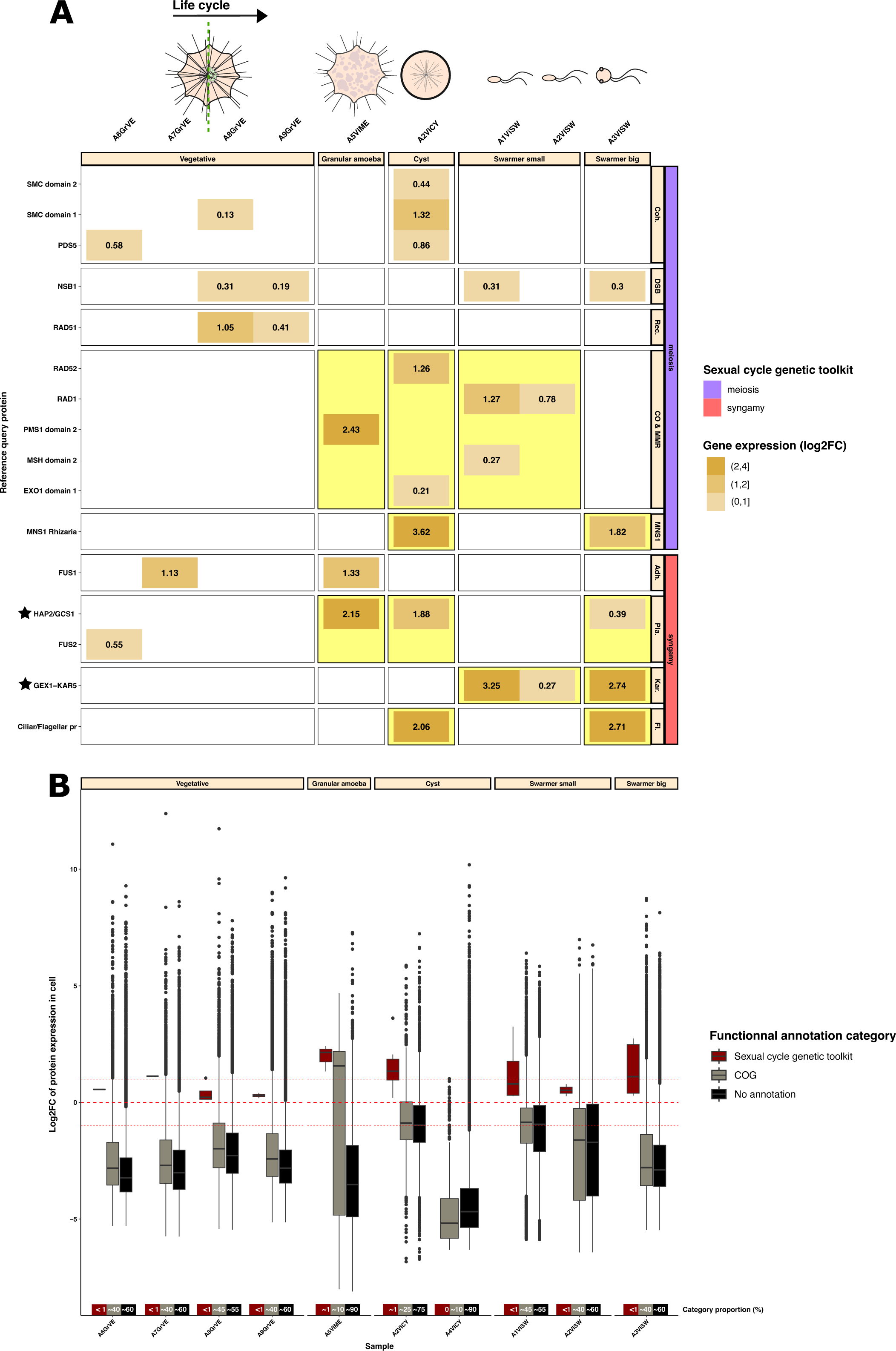
Expression of sex-related reference eukaryotic genes among acantharian reproductive life stages. (A) HMM profiles of sexual cycle genetic toolkit proteins up-regulated among the predicted proteome of acantharian life stages (fold-change > 0, i.e. gene expression is higher than average expression in cell). Proteins are categorized according to their function in the process of meiosis (i.e. Coh.: cohesion of sister chromatids; DSB: double-strand DNA break; Rec.: recombination; CO & MMR: Cross-Over resolution & mismatch repair mechanism) and syngamy (i.e. Adh.: gamete adhesion; Pla.: plasmogamy; Kar.: karyogamy; Fl.: flagellar protein). The HMM protein profile recovered only in reproductive life stages is highlighted in yellow and the stars indicate proteins that are specific to protist sexual cycles. The syngamy specific proteins HAP2/GCS1 and GEX1-KAR5 were recovered with the lineage-specific HMM profile search analysis (cf. Materials and Methods) and the query coverage values for these profiles are above 0.1 and 0.6, respectively. The complete input list of query HMM profiles can be found in Tables S2 and S4. (B) Expression of recovered sex-related HMM protein profiles in comparison to the rest of predicted proteins in acantharian life stages. Predicted proteins of acantharian transcriptomes are functionally categorized according to their level of functional annotation from COGs to precise protein functions (cf. color code). Red dashed lines indicate average transcriptome expression (i.e. 0) and 1-fold difference to average expression (i.e. −1, 1). The overall relative proportion of transcripts (i.e., TPM %) that each functional category represents is indicated above sample names (cf. Fig. S2).

### 3. Life stage-dependent ortholog gene clusterization reveals novel life stage-specific functions

Since the majority of predicted proteins in acantharian single-cell transcriptomes lacked functional annotation, we conducted a comparative sequence similarity analysis to identify unassigned protein groups that are common across life stages and specific to each life stage. Vegetative acantharian cells appeared to have the largest proportion of life stage-specific proteins (63.1% of their vegetative OGs). Big swarmers and cysts both showed a 5-times lower proportion of life stage specific proteins (12.9% of OGs). Big swarmers shared almost as many proteins with vegetative stages (11.2%) and, thus, turned out to be the reproductive stage having the closest functional profile to vegetative cells (Fig. 4A,B). In contrast, small swarmers shared the smallest predicted protein proportion with vegetative cells (2.8%) and showed 2-times more life stage specific predicted proteins compared to other reproductive stages (24.7% small swarmer specific OGs). Cysts appeared as an intermediate life stage between vegetative cells and swarmers, with 7.8% OGs shared with vegetative cells (Fig. 4A,B), while having the same predicted protein proportion in common with both types of swarmers (2% shared with big swarmers and 1.5% shared with small swarmers). Predicted protein groups shared between big and small swarmers represented 4.8% OGs, thus setting small swarmers as the second closest functional life stage to big swarmers, after vegetative stages (Fig. 4B). The predicted protein pool common between all reproductive life stages was rated among the smallest proportions (0.6%) (Fig. 4A) and exclusively included transcripts lacking COG functionnal annotation (Fig. S3).

**Fig. 4.**
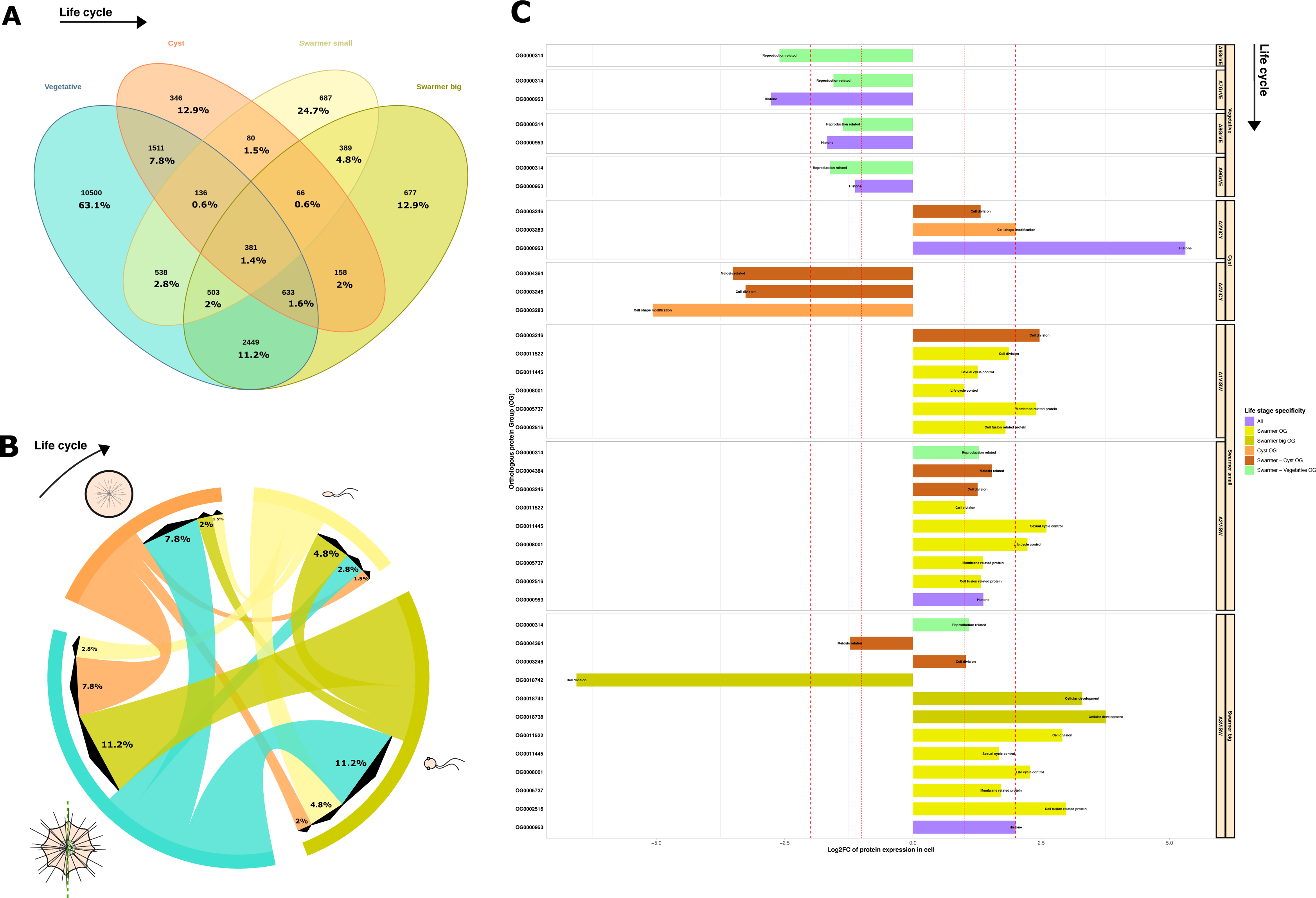
Presence of life stage-specific predicted protein families across the acantharian life cycle. (A) Numbers of Orthologous Groups (OGs) of predicted proteins shared between acantharian life stages. The proportion of OGs among the life stages is indicated by the percentages. The meiosis life stage was excluded due to its low transcript number (cf. Fig. S1). (B) Representation of pair-wise proportion of shared OGs across the acantharian life cycle. The arrow sizes indicate the dominant sharing patterns for each life stage. The percentages are added from Fig. 3A. (C) Expression of life cycle-related OGs enriched in acantharian reproductive stages (i.e., either 1 or 2 fold higher/lower than average gene expression in cell).

The functional and structural annotation of most expressed OGs revealed putative life cycle related functions (Table S3) in cyst and swarmer predicted proteins. An histone protein family common between all life stages, was found to be up-regulated among the two swarmers types and cysts, while being down-regulated in vegetative cells (i.e., OG0000953, Fig. 4B; Table S3). The same trend was observed for a potential signaling protein family (i.e., OG0000185, Fig. S4; Table S3). The vast majority of vegetative-specific predicted protein families were down-regulated in reproductive stages (Fig. S4) and only one family of unknown function was found up-regulated in all vegetative cells (i.e., OG0007516, Fig. S4; Table S3). In contrast, small and big swarmers showed 9 up-regulated predicted protein families in common with either unknown, signaling or life cycle-related consensus annotations and 8 down-regulated predicted protein families potentially involved in signaling and unknown functions (Fig. S4; Table S3). Among up-regulated swarmer-specific predcicted proteins, 5 had a life cycle-related annotation: a protein family annotated by a domain found in bacterial pili (i.e., OG0002516); a membrane-related protein family (i.e., OG0005737); a protein family bearing the combined annotations of a sensor protein linked to life-cycle transitions in bacteria and a flagella-related protein (i.e., OG0008001); a protein family bearing the combined annotations of a protein controlling yeast sexual cycle and an anaphase-promoting complex subunit (i.e., OG0011445), as well as a protein family related to the centrosome (i.e., OG0011522). Moreover, the annotations of OGs common between swarmers and vegetative cells highlighted a predicted protein family containing a yeast sporulation-specific component, up-regulated in small/big swarmers and down-regulated in vegetative cells (i.e., OG000314, Table S3). Cyst-specific predicted protein families of interest were detected solely based on cell A2ViCY as the global transcript expression of A4ViCY was very low (Fig. S1; Fig. 3B). Among the three up-regulated cyst-specific OGs, one showed structural homology with a contractile protein that could indicate a link with cell shape modification (i.e., OG0003283), while down-regulated predicted protein families remained uncharacterised (Table S3). In the group of proteins exclusive to small/big swarmers and cyst life stages were included an up-regulated predicted protein family annotated with a spindle body protein and an eukaryotic translation initiation factor (i.e., OG0003246) and a predicted protein family that was identified as a synaptonemal complex protein as well as an homologous recombination repair protein (i.e., OG0004364) that was up-regulated in small swarmers (A2ViSW) and down-regulated in big swarmers (Fig. 4B; Table S3). The expression of big swarmers specific OGs was further studied, as they appeared to be an intermediate functional stage between small swarmers and vegetative cells (Fig. 4B). Among the 9 big swarmer specific up-regulated OGs, one was annotated with a tissue development-related protein of vertebrates (i.e., OG0018738) and one with a cell morphogenesis protein (i.e., OG0018740) (Table S3). The other 7 up-regulated predicted protein families of big swarmers were uncharacterised and one was related to cell secretion, while among the 8 down-regulated predicted protein families identified one was associated to cell division (i.e., OG0018742) and the others to post-transcriptional modifications, signaling and uncharacterised functions (Fig. S4; Table S3). When looking at swarmer-specific predicted protein families differentially expressed between big swarmers and small swarmers, OGs up-regulated in big swarmers (and down-regulated in small swarmers) were related to post-transcriptional modifications, extracellular signaling and uncharacterised functions (Fig. S4; Table S3). OGs down-regulated in big swarmers (and up-regulated in small swarmers) were involved in protein-protein interactions, structural modifications or uncharacterised functions (Fig. S4; Table S3). All predicted protein families exclusively shared between big swarmers and vegetative cells were found to be down-regulated (Fig. S4) and were associated with protein modifications, signaling, protein-protein interactions and catabolism (Table S3).

## Materials & Methods

### Sampling and RNA extraction

In total, 10 acantharian cells were studied: 4 vegetative life stages, 2 cysts, 1 granular amoeba stage and 3 batches of swarmers each obtained from 3 different vegetative cells. The initial vegetative cells of the 10 acantharian samples had different morphologies (i.e., indicated in the sample names from A1 to A9) except for the vegetative cell that formed the cyst A2ViCY and the vegetative cell that produced the swarmers A2ViSW (Fig. 1B; Photo S2). Putative acantharian sexual stages were sampled on the 25/01/2022 (A1ViSW, A2ViSW, A2ViCY, A4ViCY, A5ViME) and 26/01/2022 (A3ViSW) at Villefranche-sur-MerOceanographic Observatory by subsurface horizontal towing at low speed with a 64 um plankton net, ∼1.5 km from the coast. Two sexual stages of Foraminifera (i.e., two cells) were isolated from net tows on 22/09/2022 (F1MoMESW, F2MoME; Photo S3) during the campaign MOOSE-GE 2022 (Mediterranean Ocean Observing System for the Environment, campaign DOI: <u>10.17600/18001854</u>). Directly after sampling, cells were isolated by pipetting and individually transferred in petri dishes filled with filtered sea water. Observations were done every 4 hours with binoculars and light microscopy. Life stage transitions were observed 1, 2 or 5 days after isolation (Table S1). Once observed, the different life stages were transferred in 1.5 ml microtubes filled with Tissue and Lysis Buffer (LGC Biosearch Technologies) and flash-freezed in liquid nitrogen. RNA/DNA extraction was done following a modified protocol from the Masterpure Complete RNA and DNA Purification kit (LGC Biosearch Technologies). For the vegetative acantharian cells (A6GrvE, A7GrVE, A8GrVE, A9GrVE), sampling was done in september 2022 at Villefranche-sur-Mer Oceanographic Observatory by subsurface horizontal towing at low speed with a 150 um plankton net, ∼1.5 km from the coast.

### cDNA library creation, sequencing and assembly of single-cell transcriptomes

cDNA synthesis and amplification were performed using SMARTSeq V4 Ultra Low Input RNA kit (Takara Bio). Then dual-indexing library preparation was created in equimolar conditions following the SMARTSeq Library Prep kit (Takara Bio) and Illumina sequencing 2x 150 reads was done on a Novaseq SP by Fasteris-GeneSupport SA (Switzerland). The acantharian single-cell transcriptomes of swarmer and cyst and granular amoeba stages (i.e., A1ViSW, A2ViSW, A3ViSW, A2ViCY, A4ViCY, A5ViME) and foramineferan sexual life stages (i.e., F1MoMESW, F2MoME) were assembled on the ABiMS bioinformatic platform of Roscoff Marine Station. Read filtering and quality measures were done with fastp (v0.23.) (Chen et al., 2018) and prinseq (v0.20.4) (Schmieder and Edwards, 2011). The transcriptomes were assembled using SPAdes (v3.15.2) (Prjibelski et al., 2020) and hard filtered version of the assemblies were retained (i.e., “hard filtered” results are the high-quality, reliable contigs remaining after strict filtering, likely representing accurate genomic sequences instead of errors or artifacts). (Pseudo)remapping of raw reads on assembled contigs was done with Salmon (v1.6.0, default parameters) (Patro et al., 2017). Quality and completeness of the assemblies were assessed using BUSCO (v5.4.4) (Manni et al., 2021). Coding regions were predicted by TransDecoder (v5.5.0) (Haas et al., 2013). The single-cell transcriptomes of vegetative acantharian cells (A6GrVE, A7GrVE, A8GrVE, A9GrVE) were shared by co-author Caroline Juery, after being processed with the same tools cited above apart from the (pseudo)remapping step that was done with Kallisto (default parameters) (Bray et al., 2016). Raw reads and single-cell transcriptome assemblies are available in the European Nucleotide Archive (ENA) at EMBL-EBI under the accession number PRJEB77000 (https://www.ebi.ac.uk/ena/browser/view/PRJEB77000).

### 18S/28S recovery and phylogenetic reconstruction

For reproductive life stages of Acantharia (i.e., A1ViSW, A2ViSW, A3ViSW, A2ViCY, A4ViCY, A5ViME), the ribosomal marker genes 18S and 28S were recovered *in silico* from the single-cell transcriptomes with barrnap (v0.9) (Seemann T., https://github.com/tseemann/barrnap). As multiple copies of the ribosomal genes were present in some cells (Table S1), the most expressed copy of each marker (18S and 28S) was chosen and the two markers were concatenated before phylogenetic reconstruction (Fig. S5). For vegetative life stages of Acantharia (i.e., A6GrVE, A7GrVE, A8GrVE, A9GrVE) the most abundant V4 sequences of each single-cell transcriptome were communicated by co-author Caroline Juery. Additional 18S/28S markers sequenced from environmental samples were communicated by Miguel Mendez Sandin from his thesis work (Sandin, 2019). Sequence alignment was done with MAFFT (v7.515) (Katoh, 2002) using an algorithm adapted to sequences with similar lengths and belonging to closely related groups (mafft--maxiterate 1000 —globalpair). The multiple sequence alignment was trimmed with trimAI (v1.4.1, -gt 0.3) (Capella-Gutiérrez et al., 2009) and the phylogeny was calculated with raxML-ng (v1.1.0) (Kozlov et al., 2019) under the evolutionary model GTR+G for nucleotides.

### Functional annotation, HMM profiles search and relative expression

Transcripts were functionally annotated with the eggNOG-mapper (v2.1.8) (Cantalapiedra et al., 2021). A list of 35 meiosis related proteins and 12 gamete related proteins was searched in the predicted proteomes (Table S2). The HMM profiles of these proteins were downloaded from PFAM (http://pfam.xfam.org/, (Mistry et al., 2021)) and compared to the predicted proteomes with HMMER suite tool (v3.2.1, hmmsearch program, internal analysis threshold retrieved from PFAM with –cut_ga option) ((Eddy, 2009). In order to enhance the taxonomic specificity of the HMM profiles and, thus, their sensitivity during the analysis, we created 59 lineage-specific HMM profiles for sexual cycle specific genes (Table S4). Seed sequences within the SAR clade (i.e. Rhizaria, Alveolata, Stramenopiles) were downloaded from PFAM (http://pfam.xfam.org/, (Mistry et al., 2021)). The selected SAR sequences were aligned with MAFFT (v7.515, parameter –auto) (Katoh et al., 2002) and their HMM profile was created with HMMER suite tool (v3.2.1, hmmbuild program) ((Eddy, 2009). The internal analysis threshold of the hmmsearch program ((Eddy, 2009) for these lineage-specific HMM profiles was calculated manually in an R script (R 4.1.1, Rstudio Team 2020, cf. Data availability). The results were parsed with homemade bash scripts and R scripts (cf. Data availability) for combining graphically relevant information and generating files regrouping the sequences of significant hits (applied thresholds: domain_eval <=1E-06, accuracy >=0.5, query_coverage >=0.5 for general HMM profile alignments and domain_eval <=1E-06 for lineage-specific HMM profile alignments). The graphics were created with Rstudio (R 4.1.1, Rstudio Team 2020). Gene expression was calculated in TPMs based on the remapping rate for each gene and in fold-change (i.e., compared either to average expression in vegetative cells or to average expression within each cell, log2FC = log2(TPM) - log2(AVG)).

### Comparative transcriptomics and novel predicted protein family description

In order to identify life stage specific protein families and, in parallel, account for the high proportion of poorly annotated transcripts, the predicted proteomes of acantharian single-cells were compared with OrthoFinder (v2.5.5) (Emms and Kelly, 2019). The Orthologous protein Groups (OGs) enriched in each life stage (i.e. log2FC of each OG compared to average expression in cell > 2 and if no hits were recovered the threshold was lowered to 1, Fig. S6) were described by 5 levels of annotation (Table S3): general annotation with NCBI blastp (https://blast.ncbi.nlm.nih.gov/Blast.cgi?PROGRAM=blastp&PAGE_TYPE=BlastSearch&LINK_LOC=blasthome, default parameters), protein family prediction with InterProscan (v98.0, https://www.ebi.ac.uk/interpro/) (Paysan-Lafosse et al., 2023), functional PFAM domain identification with myCLADE (http://www.lcqb.upmc.fr/myclade/large_annotation.php, complete model library, e-value < 1E-06) (Vicedomini et al., 2021) and structural homology identification Phyre (v2.0, http://www.sbg.bio.ic.ac.uk/phyre2/html/page.cgi?id=index) (Kelley et al., 2015). OGs with consensus annotation relevant to the life cycle were selected as potential acantharian life cycle markers.

## Discussion

As part of an uncultivated protist lineage, the role of acantharian reproductive life stages has remained hypothetical since the first speculations of swarmers being gametes, around 100 years ago (Schewiakoff, 1926). Here, we generated the first single-cell transcriptomes of acantharian reproductive stages in order to study the gene expression of each of these poorly understood life stages individually.

### Acantharian swarmer formation pathway is governed by genes common to both meiosis and mitosis

The transition from vegetative acantharian cells to putative meiotic life stages (i.e., granular amoeba and cyst) is accompanied by a transcriptional switch in the expression of broad functional categories (i.e., COGs). The vegetative acantharian cell transits from a metabolically active stage invested in energy production, sugar metabolism and ribosomal activity to a dividing life stage characterized by the up-regulation of genes involved in cell cycle control/cell division, replication/recombination and chromatin dynamics (Fig 1C,D). Differential single-cell expression of the amoeba *Cochlopodium* (Amoebozoa) is coherent with such opposed functional profiles, with higher metabolic activity found in vegetative stages compared to reproductive ones (Tekle et al., 2020). As in the case of *Cochlopodium* fused cells (i.e., sexual stages), acantharian putative meiotic stages were enriched in Structural Maintenance of Chromosomes (SMC), Post-Meiotic Segregation Increased 1 (PMS1) and Exonuclease 1 (EXO1) protein domains, while sister chromatid cohesion protein PDS5 domain was equally expressed in both vegetative and putative meiotic stages. The SMC domains involved in the cohesion of sister chromatids were enriched in the cyst cell on-going formation, along with cross-over and mismatch repair protein domains RAD52 (i.e., binds to DSB breaks) and EXO1 (i.e., DNA exonuclease), while the mismatch repair protein domain PMS1 was highly enriched in the non-cyst meiotic stage. Nevertheless, in contrast to *Cochlopodium* fused cells (Tekle et al., 2020), none of the essential meiosis genes required for sister chromatid cohesion (REC8), synaptonemal complex formation (HOP1, SPO22, PCH2), double-strand breaks (SPO11), recombination (HOP2, MND1, DMC1) and cross-over resolution (MSH4, MSH5, MER3) (Hofstatter et al., 2018) was found enriched in acantharian putative meiotic stages. As part of a distantly related major eukaryotic lineage (i.e., Rhizaria) and highly diverse protist clade (Burki et al., 2020), Radiolaria likely harbor lineage-specific meiotic genes, potentially arising through gene neo-functionalization mechanisms (Maciver et al., 2019), as observed in other protist lineages such as Amoebozoa and Ciliophora, which deviate from the essential meiotic gene-set (Hofstatter et al., 2018; Hofstatter and Lahr, 2019). Such meiosis-related acantharian genes could be hidden among the massive proportion of non-referenced proteins in putative meiotic stages, that represent ∼85% of predicted proteins of a single-cell (Fig 2B). A first annotation of the most expressed protein clusters among acantharian putative meiotic life stages highlighted only one candidate meiosis-related predicted protein group (i.e., OG0004364 involved in synaptonemal complex and homologous recombination), exclusively expressed by both cysts and swarmers, as well as the histone H2A up-regulated in both cyst and swarmers. The expression of histones H2A seems to be linked to the sexual cycle, as they have also been found enriched in rice sperm and egg cells and were shown to be particularly up-regulated in rice zygote after fertilization (Abiko et al., 2013). Even though meiosis-related genes were expressed in acantharian putative meiotic stages, the current genetic data does not allow to determine the meiotic or mitotic nature of the on-going molecular process in the cyst and granular amoeba life stages. Therefore, we will maintain the designation of these life stages as pre-swarmer reproductive stages in order to avoid ambiguity. At this point, we can propose 2 possible scenarios for the molecular process underlying the transition of the acantharian vegetative cell to the pre-swarmer reproductive stages and the role of swarmers: (1) the vegetative acantharian cell is either diploid or haploid and undergoes mitosis to produce swarmers, that are not gametes but develop into juveniles, or (2) the vegetative acantharian cell is diploid and undergoes meiosis by expressing currently undescribed acantharian-specific meiosis genes to produce swarmers, that are gametes and fuse to give rise to a zygote.

### Identification of gamete specific genes and putative sexual cycle genes in acantharian swarmer transcriptomes

Following the sequence of events in the acantharian life cycle (Fig 1A), the granular amoeba and cyst stages rapidly give rise to swarmers. This progressive transition is illustrated by the proximity of functional profiles between swarmers and pre-swarmer stages (Fig 1C) and the presence of swarmer-related proteins in pre-swarmer reproductive life stages (Fig 2A). Like cysts and granular amoeba stages, swarmer transcriptomes are enriched in COGs relative to cell cycle control/cell division, as well as transcription, lipid transport and cell motility, while energy production, sugar metabolism and ribosomal activity appear down-regulated. Large scale scRNA-seq conducted throughout the life cycles of malaria parasites have also demonstrated a higher replication activity in gamete life stages (Howick et al., 2019). Among acantharian swarmers, relatively more swarmer-specific predicted proteins were found compared to the number of predicted proteins shared between swarmers and pre-swarmer reproductive stages. The cell division and meiosis-related functions identified among common swarmer and pre-swarmer predicted proteins (i.e., OG0003246 and OG0004364) suggested that swarmers share genes with cysts, while the high proportion of swarmer-specific predicted proteins implied a specialization in different life cycle functions for swarmers and pre-swarmer life stages. In fact, the recovery of gamete reference genes, revealed an enrichment of genes essential for syngamy among swarmers. Notably, the eukaryotic gamete fusogen essential for gamete membrane fusion HAP2/GCS1 and the GEX1-KAR5 protein family mediating gamete nuclear fusion (Speijer et al., 2015), were both up-regulated in swarmers. The Hapless 2 / Germ Cell Specific 1 (HAP2/GCS1) is a fusion protein (i.e. fusogen) involved in sperm-egg fusion in flowering plants (e.g. *Arabidopsis*, Fedry et al., 2018), invertebrates (e.g. *Drosophila*, Hernández and Podbilewicz, 2017) and gamete fusion in many protists such as the green microalgae *Chlamydomonas* (Archaeplastida, Liu et al., 2008), the ciliate *Tetrahymena* (Alveolata, Pinello et al., 2023), the social amoeba *Dictyostelium* (Amoebozoa, Okamoto et al. 2016) and the parasites *Trypanosoma* (Excavata, Howick et al. 2021) and *Plasmodium* (Alveolata, Liu et al. 2008). Its wide distribution across eukaryotic lineages with divergent life cycles suggests that HAP2/GCS1 is an ancestral gamete fusogen of eukaryotes (Speijer et al., 2015). The gene Gamete EXpressed Protein 1 (GEX1) is related to fertilization in flowering plants (e.g. *Arabidopsis*) and has been related to both nuclear fusion and meiosis during evolution. It is essential for gamete fusion in several species such as green microalgae (i.e., *Chlamydomonas*) and alveolates (i.e., *Plasmodium*). Moreover, it is a member of the same family as Karyogamy 5 (KAR5), an essential gene for nuclear fusion during yeast sexual recombination (Ning et al., 2013; Nishikawa et al., 2020). The expression of these gamete specific genes strongly suggests that swarmers are able to fuse, while the enrichment of meiosis-related MSH2 and RAD1 cross-over and mismatch repair domains indicates a potential for gene shuffling. The functional differentiation between pre-swarmer stages and swarmers probably lies within the implication of swarmers in a sexual cycle. In contrast to pre-swarmer stages, some mitosis related functions appear down-regulated among swarmers (i.e., replication related OG0011509 and mitotic checkpoint protein OG0001070), while some swarmer specific up-regulated protein clusters appear to be related to life cycle transitions and sexual cycle regulation. Therefore, *de novo* annotation of uncharacterised swarmer-specific protein clusters is also consistent with the implication of swarmers in the sexual cycle, most probably as gametes. The up-regulated membrane related swarmer genes could stand as acantharian specific fusion genes. Thus, gamete formation from pre-swarmer stages seems to be progressive, as suggested by the expression of mitosis related genes among swarmers but also essential gamete fusion genes in pre-swarmer stages. Since vegetative acantharian cells have multiple nuclei, we speculate that acantharian swarmer formation occurs progressively and independently in each nuclei, with the continuous fragmentation of replicative vesicles, up to the individualisation of swarmers (also proposed by Caron and Swanberg, 1990). Such a progressive gamete production process could explain the expression of the HAP2/GCS1 fusogen in pre-swarmer stages in addition to the up-regulation of the ciliar/flagellar domain (CFA20) in the cyst life stage, suggesting that the morphological transformation of the cell occurs in parallel to gamete formation in the cyst. We, therefore, propose that acantharian swarmer production follows the process observed for the rhizarian lineage of Cercozoa, in which swarmers emerge from the fragmentation of plasmodial spheres containing multiple nuclei (Boltovskoy et al., 2017).

### Comparison of life stage functional profiles converges towards putative early-stage acantharian zygote

Even though both small and big swarmers express gamete specific and putative gamete-related genes, their functional profiles appear to be divergent, as their morphologies. The *de novo* annotated meiosis related predicted protein cluster OG0004364 was down-regulated in big swarmers, in contrast to small swarmer and pre-swarmer stages. Additionally, the expression of some swarmer specific predicted protein clusters related to post-translational modification (i.e., OG0008021 and OG0007993) and lipid metabolism (i.e., OG0007950) was contrasted between small and big swarmers, a tendency also observed with the differential expression of COGs. More strikingly, the expression of motility, replication/recombination and chromatin dynamics COGs of big swarmers was closer to vegetative cells than small swarmers. Overall, the COG expression profile of big swarmers matched more with vegetative cells, reflecting the larger number of predicted protein clusters shared between big swarmers and vegetative stages. However, gamete specific genes HAP2/GCS1 and GEX1-KAR5 were both up-regulated in big swarmers. These findings prompt us to propose that big swarmers are an intermediate developmental stage between small swarmers and vegetative cells. Given that both the cell and transcriptome sizes of big swarmers were twice larger than small swarmers, we investigated the potential role of big swarmers as acantharian zygotes. Similar to big swarmers, scRNA-seq data of *Plasmodium* zygotic life stages was poorly enriched in cell motility gene ontology (GO) terms, while lipid metabolism was up-regulated compared to gamete stages (Howick et al., 2019). Additionally, *de novo* annotation of predicted protein clusters exclusively identified in big swarmers uncovered one replication-associated protein exhibiting a down-regulated expression of over fivefold, in contrast to two cellular development/growth functions showing an up-regulated expression of over twofold (Fig. 4C; Table S3). More specifically, one of the cellular development/growth functions is a cell morphogenesis domain identified in the predicted protein cluster OG0018740. This domain is not associated with sexual recombination, but plays an essential role in establishing growth polarity during fission in yeasts (Hirata, 2002). Indeed, the morphogenesis domain acts by coordinating the recruitment of actin in the cytoskeleton in order to structure and maintain growth zones within the cell (Hirata, 2002). Cell polarization, facilitated by this process, has been described as necessary for zygote development in flowering plants (Ueda et al., 2011) and brown algae (Bogaert et al., 2023). The other development/growth related predicted protein cluster exclusively expressed in big swarmers, OG0018738, contained the functional domain of noggin proteins. Noggin proteins are known to control multiple cell signaling pathways (Karunaraj et al., 2022) and play a crucial role in the proper development of vertebrate embryonic tissues (Ishibashi et al., 2008). Therefore, the putative functions of these big swarmer specific proteins do not align with those of gametes but could be consistent with a putative early-stage zygote. The combined expression of OG0018738 and OG0018740 proteins may contribute to furnishing the newly fertilized cell with the requisite information to facilitate the vegetative growth of the acantharian zygote. We sustain the hypothesis that these putative acantharian zygotes are not issued from multiple fission, like commonly documented in the sister lineage of Foraminifera (Meilland et al., 2024), since the expression of gamete fusion genes in cells capable of development without fertilization wouldn’t be expected. Instead, the expression of these gamete specific genes could be explained by an incomplete gene suppression, as it has been described in zygotes of the green microalgae *Chlamydomonas* (Joo et al., 2017).

Hence, by integrating the inferred genetic roles of the life stages studied here in the state-of-the-art hypothetical acantharian life cycle, we speculate that the vegetative acantharian cell is diploid and undergoes a monophasic life cycle (i.e., including one diploid life phase). The cysts and granular amoebas go through a progressive meiotic process, mediated by currently undescribed lineage specific meiosis genes and give rise to small swarmer gametes. In certain cases, Acantharia seem to be capable of autogamy, producing putative big swarmer zygotes characterized by a rotating swimming motion and the presence of two small vesicles potentially representing the strontium inclusions originating from each small swarmer gamete. Although we did not directly observe the fusion of small swarmers, the unprecedented life stage-specific genetic data produced here the first insights into the role of acantharian putative sexual stages. Overall, this data contributes to the understanding of the acantharian life cycle by linking swarmers and vegetative cells, while supporting the existence of a sexual cycle among Acantharia.

### Current limits and ecological perspectives of the acantharian sexual cycle

Despite the advances in sequencing technologies and computational methods, knowledge regarding Acantharian life cycles is scarce, reflecting the difficulty of studying sexual recombination among uncultivated protists. In order to support indirect morphological evidence of sexual cycles with genetic data, it is crucial to obtain and correctly identify putative sexual stages. In the case of Acantharia, the trigger for swarmer formation remains unclear and, thus, cannot be fully controlled experimentally. While nutrient and temperature stressors seem to trigger the “sexual maturation” of vegetative cells, only a few cells within the population swarm and the process remains asynchronous (from personal observation). Furthermore, the precise morphological identification of pre-swarmer stages represents a challenge, especially among non-cysting Acantharia. As a consequence, the lack of replicates is nearly unavoidable and can only be mitigated by isolating sexual life stages from multiple taxa. Life cycle transitions have been even less frequently observed for the other three groups of Radiolaria, as swarmer formation has been recently documented for only four species of Spumellaria (*Didymocyrtis ceratospyris, Tetrapyle* sp., *Triastrum aurivillii,* (Yuasa and Takahashi, 2016); *Cypassus irregularis,* (Kimoto et al., 2011)), one species of Nassellaria (*Pterocanium praetextum,* (Yuasa and Takahashi, 2016)) and mainly two species of Collodaria (*Thalassicola nucleata,* (Anderson, 1983); *Sphaerozoum punctatum,* (Yuasa and Takahashi, 2014)). Concentrating efforts towards culturing Radiolaria would represent a tremendous advance in understanding their life cycle. In addition to facilitating the documentation of additional developmental stages within their sexual life cycle, it would also enable the collection of a sufficient number of single-cell replicates for statistical analysis, as has been done for *Plasmodium* and *Trypanosoma* parasites (Howick et al., 2019; Howick et al., 2021). Moreover, establishing a radiolarian culture would significantly accelerate the generation of radiolarian genomes, enabling the validation of recovered transcript sequences and providing a reference framework for functional annotation. The study of radiolarian biology is challenging and thus the majority of radiolarian genes are yet uncharacterized. Meanwhile, lineage specific variations of reference sexual cycle genes have been documented among some protist lineages (Malik et al., 2008; Thangavel et al., 2023). Subsequently, even though the expression of reference sexual cycle genes stands as direct evidence for the existence of a sexual cycle, the presence of sex-related genes requires confirmation through approaches such as *in situ* hybridisation in putative meiotic stages (Tekle et al., 2020) and gametes (Howick et al., 2021). This would also provide a comprehensive assessment of whether sequence conservation is a relevant proxy for function conservation in the lineage (Maciver et al., 2019). Validating acantharian gamete genes and further describing acantharian sexual cycle genetics has important implications for understanding the mysterious life cycles within the supergroup of Rhizaria (Burki and Keeling, 2014). Among free-living representatives of Rhizaria, a complete life cycle description has been provided only for benthic Foraminifera (Lehmann et al., 2006), while the available genetic resources for other lineages (e.g. Cercozoa, Aquavolonida) are limited (Richter et al., 2022). Focusing on lineage-specific sexual cycle genes, would allow to establish the role of various free-living rhizarian life stages not only in their life cycle, but also in their environment. Understanding the life cycle of a protist lineage provides valuable information on its ecology (Richards et al., 2019). Recent estimates have placed Rhizaria and more specifically Radiolaria and Foraminifera, as primary actors of the biological carbon pump during the past 30 years (Lampitt et al., 2023). For Radiolaria this major implication in global biogeochemical cycles is likely to vary during the life cycle, as they are expected to sink before swarmer release (Martin et al., 2010; (Yuasa and Takahashi, 2016). In the case of Acantharia, cell density is significantly increased by cyst formation, contributing to rapid particle sedimentation and the creation of strontium gradients (Decelle et al., 2013; Martin et al., 2010). Considering this, the development of rhizarian sexual cycle genetic probes would enable the estimation of the rhizarian lifespan and contribution to global biogeochemical cycles at a life stage resolution. This would represent a significant advancement in rhizarian single-cell ecology.

### Conclusion

Acantharia demonstrate both morphological and genetic evidence supporting the existence of a sexual cycle in which swarmers are gametes, produced either via the vegetative cell or via the intermediate of a cyst stage. Even though swarmers express gamete specific reference genes, the meiotic nature of the molecular process underlying swarmer formation remains undetermined. The identification and functional annotation of life-stage specific genes, highlighted a new life stage unprecedented in prior descriptions. Based on its morphological resemblance with swarmers and its transcriptional proximity with vegetative cells, we speculate that this newly observed acantharian life stage is a putative zygote resulting from autogamy.

These findings allow us to propose a novel model for the acantharian life cycle integrated with life-stage specific genetic data.

## Data availability

The data for this study have been deposited in the European Nucleotide Archive (ENA) at EMBL-EBI under accession number PRJEB77000 (https://www.ebi.ac.uk/ena/browser/view/PRJEB77000). Scripts and Rmarkdown files necessary to run all the analyses included in this work are publicly available on the GitHub page: https://github.com/IrisRizos/SexCy_lifestage_protistID.

## Acknowledgments

The authors thank EMBRC (https://www.embrc.eu/) and its French partner site, Institut de la Mer de Villefranche (IMEV, https://www.imev-mer.fr/web/), for providing the opportunity to organize sampling trips in the Mediterranean Sea and for on-site access to the EMBRC visitor installations. The authors acknowledge the MOOSE program (Mediterranean Ocean Observing System for the Environment, <u>10.18142/235</u>) coordinated by CNRS-INSU and the Research Infrastructure ILICO (CNRS-IFREMER). Also, the authors thank Adriana Alberti for her advice and are grateful to Miguel Mendez Sandin for his valuable insights into phylogenetic reconstructions and for sharing acantharian 18S/28S marker sequences which served as references in our phylogeny. All analyses of this work were performed on the AbiMS cluster of the marine station of Roscoff (http://abims.sb-roscoff.fr). Lucie Bittner acknowledges the Institut Universitaire de France for her 5-year nomination as Junior Member (2020-2025).

## CRediT authorship contribution statement

**Iris Rizos:** Conceptualization, Data curation, Formal analysis, Investigation, Methodology, Visualization, Writing - original draft. **Sarah Romac:** Data curation, Investigation, Methodology. **Caroline Juery:** Data curation, Resources. **Charlotte Berthelier:** Data curation, Methodology. **Johan Decelle:** Resources, Supervision. **Juliana Bernardes:** Methodology, Supervision. **Erwan Corre:** Data curation, Methodology. **Lucie Bittner:** Conceptualization, Methodology, Supervision. **Fabrice Not:** Conceptualization, Investigation, Methodology, Project administration, Supervision. All authors have participated in Writing - review and editing and to the Validation of this work.

## References

Abiko, M., Maeda, H., Tamura, K., Hara-Nishimura, I., Okamoto, T., 2013. Gene expression profiles in rice gametes and zygotes: identification of gamete-enriched genes and up-or down-regulated genes in zygotes after fertilization. Journal of Experimental Botany 64, 1927–1940. 10.1093/jxb/ert054

Anderson, O.R., 1983. Radiolaria. Springer New York, New York, NY. 10.1007/978-1-4612-5536-9

Archibald, J.M., Simpson, A.G.B., Slamovits, C.H. (Eds.), 2017. Handbook of the Protists, 18. Springer International Publishing, Cham. 10.1007/978-3-319-28149-0

Bogaert, K.A., Zakka, E.E., Coelho, S.M., De Clerck, O., 2023. Polarization of brown algal zygotes. Seminars in Cell & Developmental Biology 134, 90–102. 10.1016/j.semcdb.2022.03.008

Boltovskoy, D., Anderson, O.R., Correa, N.M., 2017. Radiolaria and Phaeodaria, in: Archibald, J.M., Simpson, A.G.B., Slamovits, C.H. (Eds.), Handbook of the Protists. Springer International Publishing, Cham, pp. 731–763. 10.1007/978-3-319-28149-0_19

Bray, N.L., Pimentel, H., Melsted, P., Pachter, L., 2016. Near-optimal probabilistic RNA-seq quantification. Nat Biotechnol 34, 525–527. 10.1038/nbt.3519

Burki, F., Keeling, P.J., 2014. Rhizaria. Current Biology 24, R103–R107. 10.1016/j.cub.2013.12.025

Burki, F., Roger, A.J., Brown, M.W., Simpson, A.G.B., 2020. The New Tree of Eukaryotes. Trends in Ecology & Evolution 35, 43–55. 10.1016/j.tree.2019.08.008

Cantalapiedra, C.P., Hernández-Plaza, A., Letunic, I., Bork, P., Huerta-Cepas, J., 2021. eggNOG-mapper v2: Functional Annotation, Orthology Assignments, and Domain Prediction at the Metagenomic Scale. Molecular Biology and Evolution 38, 5825–5829. 10.1093/molbev/msab293

Capella-Gutiérrez, S., Silla-Martínez, J.M., Gabaldón, T., 2009. trimAl: a tool for automated alignment trimming in large-scale phylogenetic analyses. Bioinformatics 25, 1972–1973. 10.1093/bioinformatics/btp348

Caron, D.A., Swanberg, N.R., 1990. The ecology of planktonic sarcodines. Reviews in Aquatic Sciences 3(2-3): 147–180. https://eurekamag.com/research/033/763/033763230.php

Chen, S., Zhou, Y., Chen, Y., Gu, J., 2018. fastp: an ultra-fast all-in-one FASTQ preprocessor. Bioinformatics 34, i884–i890. 10.1093/bioinformatics/bty560

Darling, K.F., Husum, K., Fenton, I.S., 2023. The biphasic life cycle of the non-spinose planktonic foraminifera is characterised by an aberrant coiling signature. Marine Micropaleontology, 54 185, 102295. 10.1016/j.marmicro.2023.102295

Decelle, J., Colin, S., Foster, R.A., 2015. Photosymbiosis in Marine Planktonic Protists, in: Ohtsuka, S., Suzaki, T., Horiguchi, T., Suzuki, N., Not, F. (Eds.), Marine Protists. Springer Japan, Tokyo, pp. 465–500. 10.1007/978-4-431-55130-0_19

Decelle, J., Martin, P., Paborstava, K., Pond, D.W., Tarling, G., Mahé, F., de Vargas, C., Lampitt, R., Not, F., 2013. Diversity, Ecology and Biogeochemistry of Cyst-Forming Acantharia (Radiolaria) in the Oceans. PLoS ONE, 34 8, e53598. 10.1371/journal.pone.0053598

Decelle, J., Not, F., 2015. A cantharia, in: John Wiley & Sons, Ltd (Ed.), eLS. Wiley, pp. 1–10. 10.1002/9780470015902.a0002102.pub2

Eddy, S.R., 2009. A NEW GENERATION OF HOMOLOGY SEARCH TOOLS BASED ON PROBABILISTIC INFERENCE, in: Genome Informatics 2009. Presented at the Proceedings of the 20th International Conference, PUBLISHED BY IMPERIAL COLLEGE PRESS AND DISTRIBUTED BY WORLD SCIENTIFIC PUBLISHING CO., Pacifico Yokohama, Japan, pp. 205–211. 10.1142/9781848165632_0019

Emms, D.M., Kelly, S., 2019. OrthoFinder: phylogenetic orthology inference for comparative genomics. Genome Biol 20, 238. 10.1186/s13059-019-1832-y

Febvre, J., 1977. La division nucléaire chez les Acanthaires. Journal of Ultrastructure Research 60, 279–295. 10.1016/S0022-5320(77)80014-2

Febvre-Chevalier, C., and Febvre, J., 1994. Buoyancy and Swimming in Marine Planktonic Protists in: Maddock, L., Bone, Q., Rayner, J.M.V. (Eds.), The Mechanics and Physiology of Animal Swimming. L. Cambridge University Press, pp. 13–26

Fedry, J., Forcina, J., Legrand, P., Péhau-Arnaudet, G., Haouz, A., Johnson, M., Rey, F.A., Krey, T., 2018. Evolutionary diversification of the HAP2 membrane insertion motifs to drive gamete fusion across eukaryotes. PLoS Biol, 80 16, e2006357. 10.1371/journal.pbio.2006357

Fusco, G., Minelli, A., 2019. The Biology of Reproduction, 1st ed. Cambridge University Press. 10.1017/9781108758970

Goodenough, U., Heitman, J., 2014. Origins of Eukaryotic Sexual Reproduction. Cold Spring Harbor Perspectives in Biology, 12 6, a016154–a016154. 10.1101/cshperspect.a016154

Grell, K.G., 1973. Protozoology 2. Springer Berlin Heidelberg, Berlin, Heidelberg. 10.1007/978-3-642-61958-8

Haas, B.J., Papanicolaou, A., Yassour, M., Grabherr, M., Blood, P.D., Bowden, J., Couger, M.B., Eccles, D., Li, B., Lieber, M., MacManes, M.D., Ott, M., Orvis, J., Pochet, N., Strozzi, F., Weeks, N., Westerman, R., William, T., Dewey, C.N., Henschel, R., LeDuc, R.D., Friedman, N., Regev, A., 2013. De novo transcript sequence reconstruction from RNA-seq using the Trinity platform for reference generation and analysis. Nat Protoc 8, 1494–1512. 10.1038/nprot.2013.084

Haeckel, E, 1887. Report on the Radiolaria collected by H.M.S. Challenger during the years 1873–1876. Rep. Sci. Res. Voy HMS Challenger Zool 18: 1-1803

Hernández, J.M., Podbilewicz, B., 2017. The hallmarks of cell-cell fusion. Development 144, 4481–4495. 10.1242/dev.155523

Hirata, D., 2002. Fission yeast Mor2/Cps12, a protein similar to Drosophila Furry, is essential for cell morphogenesis and its mutation induces Wee1-dependent G2 delay. The EMBO Journal 21, 4863–4874. 10.1093/emboj/cdf495

Hofstatter, P.G., Brown, M.W., Lahr, D.J.G., 2018. Comparative Genomics Supports Sex and Meiosis in Diverse Amoebozoa. Genome Biology and Evolution, 68 10, 3118–3128. 10.1093/gbe/evy241

Hofstatter, P.G., Lahr, D.J.G., 2019. All Eukaryotes Are Sexual, unless Proven Otherwise: Many So-Called Asexuals Present Meiotic Machinery and Might Be Able to Have Sex. BioEssays, 69 41, 1800246. 10.1002/bies.201800246

Hohenegger, J., Kinoshita, S., Briguglio, A., Eder, W., Wöger, J., 2019. Lunar cycles and rainy seasons drive growth and reproduction in nummulitid foraminifera, important producers of carbonate buildups. Sci Rep, 61 9, 8286. 10.1038/s41598-019-44646-w

Hollande A, Cachon-Enjumet, M., 1957. Enkystement et reproduction isosporogénétique chez les acanthaires. CR Acad Sci 244: 508–510

Hollande, A., Cachon, J., Cachon-Enjumet, M., 1965. Les modalités de l’enkystement présporogénétique chez les acanthaires. Protistologica 1: 91–112

Howick, V.M., Peacock, L., Kay, C., Collett, C., Gibson, W., Lawniczak, M.K.N., 2021. Single-cell transcriptomics reveals expression profiles of Trypanosoma brucei sexual stages, 82. 10.1101/2021.10.13.463681

Howick, V.M., Russell, A.J.C., Andrews, T., Heaton, H., Reid, A.J., Natarajan, K., Butungi, H., Metcalf, T., Verzier, L.H., Rayner, J.C., Berriman, M., Herren, J.K., Billker, O., Hemberg, M., Talman, A.M., Lawniczak, M.K.N., 2019. The Malaria Cell Atlas: Single parasite transcriptomes across the complete *Plasmodium* life cycle. Science 365, eaaw2619. 10.1126/science.aaw2619

Ishibashi, H., Matsumura, N., Hanafusa, H., Matsumoto, K., Robertis, E.M.D., Kuroda, H., 2008. Expression of Siamois and Twin in the blastula Chordin/Noggin signaling center is required for brain formation in Xenopus laevis embryos. Mechanisms of Development 125, 58–66. 10.1016/j.mod.2007.10.005

Joo, S., Nishimura, Y., Cronmiller, E., Hong, R.H., Kariyawasam, T., Wang, M.H., Shao, N.C., El Akkad, S.-E.-D., Suzuki, T., Higashiyama, T., Jin, E., Lee, J.-H., 2017. Gene Regulatory Networks for the Haploid-to-Diploid Transition of *Chlamydomonas reinhardtii*. Plant Physiol. 175, 314–332. 10.1104/pp.17.00731

Karunaraj, P., Tidswell, O., Duncan, E.J., Lovegrove, M.R., Jefferies, G., Johnson, T.K., Beck, C.W., Dearden, P.K., 2022. Noggin proteins are multifunctional extracellular regulators of cell signaling. Genetics 221, iyac049. 10.1093/genetics/iyac049

Katoh, K., 2002. MAFFT: a novel method for rapid multiple sequence alignment based on fast Fourier transform. Nucleic Acids Research 30, 3059–3066. 10.1093/nar/gkf436

Kelley, L.A., Mezulis, S., Yates, C.M., Wass, M.N., Sternberg, M.J.E., 2015. The Phyre2 Web Portal for Protein Modeling, Prediction and Analysis. Nature Protocols, 10, 6, 845–58. 10.1038/nprot.2015.053

Kimoto, K., Yuasa, T., Takahashi, O., 2011. Molecular identification of reproductive cells released from Cypassis irregularis Nigrini (Radiolaria): Pico-size reproductive cells released from radiolaria. Environmental Microbiology Reports 3, 86–90. 10.1111/j.1758-2229.2010.00191.x

Kozlov, A.M., Darriba, D., Flouri, T., Morel, B., Stamatakis, A., 2019. RAxML-NG: a fast, scalable and user-friendly tool for maximum likelihood phylogenetic inference. Bioinformatics 35, 4453–4455. 10.1093/bioinformatics/btz305

Krueger-Hadfield, S.A., 2024. Let’s talk about sex: Why reproductive systems matter for understanding algae. Journal of Phycology 60, 581–597. 10.1111/jpy.13462

Lahr, D.J.G., Parfrey, L.W., Mitchell, E.A.D., Katz, L.A., Lara, E., 2011. The chastity of amoebae: re-evaluating evidence for sex in amoeboid organisms. Proc. R. Soc. B., 19 278, 2081–2090. 10.1098/rspb.2011.0289

Lampitt, R.S., Briggs, N., Cael, B.B., Espinola, B., Hélaouët, P., Henson, S.A., Norrbin, F., Pebody, C.A., Smeed, D., 2023. Deep ocean particle flux in the Northeast Atlantic over the past 30 years: carbon sequestration is controlled by ecosystem structure in the upper ocean. Front. Earth Sci. 11, 1176196. 10.3389/feart.2023.1176196

Lehmann, G., Rottger, R., Hohenegger, J., 2006. LIFE CYCLE VARIATION INCLUDING TRIMORPHISM IN THE FORAMINIFER TROCHAMMINA INFLATA FROM NORTH EUROPEAN SALT MARSHES. The Journal of Foraminiferal Research, 42 36, 279–290. 10.2113/gsjfr.36.4.279

Lee, J. J., Leedale, G. F., Bradbury, P. C. 2000. An Illustrated Guide to the Protozoa: Organisms Traditionally Referred to As Protozoa, Or Newly Discovered Groups. 2nd ed. Society of Protozoologists, Lawrence (Kan.): Allen press. USA. https://protistologists.org/publications/illustrated-guide-to-the-protozoa/

Liu, Y., Tewari, R., Ning, J., Blagborough, A.M., Garbom, S., Pei, J., Grishin, N.V., Steele, R.E., Sinden, R.E., Snell, W.J., Billker, O., 2008. The conserved plant sterility gene *HAP2* functions after attachment of fusogenic membranes in *Chlamydomonas* and *Plasmodium* gametes. Genes Dev. 22, 1051–1068. 10.1101/gad.1656508

Maciver, S.K., Koutsogiannis, Z., De Obeso Fernández Del Valle, A., 2019. ‘Meiotic genes’ are constitutively expressed in an asexual amoeba and are not necessarily involved in sexual reproduction. Biol. Lett. 15, 20180871. 10.1098/rsbl.2018.0871

Malik, S.-B., Pightling, A.W., Stefaniak, L.M., Schurko, A.M., Logsdon, J.M., 2008. An Expanded Inventory of Conserved Meiotic Genes Provides Evidence for Sex in Trichomonas vaginalis. PLoS ONE, 67 3, e2879. 10.1371/journal.pone.0002879

Manni, M., Berkeley, M.R., Seppey, M., Simão, F.A., Zdobnov, E.M., 2021. BUSCO Update: Novel and Streamlined Workflows along with Broader and Deeper Phylogenetic Coverage for Scoring of Eukaryotic, Prokaryotic, and Viral Genomes. Molecular Biology and Evolution 38, 4647–4654. 10.1093/molbev/msab199

Margulis, L., Sagan, D., 1986. Origins of sex: three billion years of genetic recombination, The bio-origins series. Yale Univ. Press, New Haven London.

Martin, P., Allen, J.T., Cooper, M.J., Johns, D.G., Lampitt, R.S., Sanders, R., Teagle, D.A.H., 2010. Sedimentation of acantharian cysts in the Iceland Basin: Strontium as a ballast for deep ocean particle flux, and implications for acantharian reproductive strategies. Limnol. Oceanogr. 55, 604–614. 10.4319/lo.2010.55.2.0604

Meilland, J., Siccha, M., Morard, R., Kucera, M., 2024. Continuous reproduction of planktonic foraminifera in laboratory culture. J Eukaryotic Microbiology 71, e13022. 10.1111/jeu.13022

Mistry, J., Chuguransky, S., Williams, L., Qureshi, M., Salazar, G.A., Sonnhammer, E.L.L., Tosatto, S.C.E., Paladin, L., Raj, S., Richardson, L.J., Finn, R.D., Bateman, A., 2021. Pfam: The protein families database in 2021. Nucleic Acids Research 49, D412–D419. 10.1093/nar/gkaa913

Montagnes, D.J.S., Lowe, C.D., Martin, L., Watts, P.C., Downes-Tettmar, N., Yang, Z., Roberts, E.C., Davidson, K., 2011. Oxyrrhis marina growth, sex and reproduction. Journal of Plankton Research, 36 33, 615–627. 10.1093/plankt/fbq111

Ning, J., Otto, T.D., Pfander, C., Schwach, F., Brochet, M., Bushell, E., Goulding, D., Sanders, M., Lefebvre, P.A., Pei, J., Grishin, N.V., Vanderlaan, G., Billker, O., Snell, W.J., 2013. Comparative genomics in *Chlamydomonas* and *Plasmodium* identifies an ancient nuclear envelope protein family essential for sexual reproduction in protists, fungi, plants, and vertebrates. Genes Dev., 62 27, 1198–1215. 10.1101/gad.212746.112

Nishikawa, S., Yamaguchi, Y., Suzuki, C., Yabe, A., Sato, Yuzuru, Kurihara, D., Sato, Yoshikatsu, Susaki, D., Higashiyama, T., Maruyama, D., 2020. Arabidopsis GEX1 Is a Nuclear Membrane Protein of Gametes Required for Nuclear Fusion During Reproduction. Front. Plant Sci. 11, 548032. 10.3389/fpls.2020.548032

Patro, R., Duggal, G., Love, M.I., Irizarry, R.A., Kingsford, C., 2017. Salmon provides fast and bias-aware quantification of transcript expression. Nat Methods 14, 417–419. 10.1038/nmeth.4197

Paysan-Lafosse, T., Blum, M., Chuguransky, S., Grego, T., Pinto, B.L., Salazar, G.A., Bileschi, M.L., Bork, P., Bridge, A., Colwell, L., Gough, J., Haft, D.H., Letunić, I., Marchler-Bauer, A., Mi, H., Natale, D.A., Orengo, C.A., Pandurangan, A.P., Rivoire, C., Sigrist, C.J.A., Sillitoe, I., Thanki, N., Thomas, P.D., Tosatto, S.C.E., Wu, C.H., Bateman, A., 2023. InterPro in 2022. Nucleic Acids Research 51, D418–D427. 10.1093/nar/gkac993

Pinello, J., Loidl, J., Seltz, E., Cassidy-Hanley, D., Kolbin, D., Abdelatif, A., Rey, F., An, R., Newberger, N., Bisharyan, Y., Papoyan, H., Byun, H., Aguilar, H., Cole, E., Clark, T., 2023. Novel requirements for HAP2-mediated gamete fusion in Tetrahymena. (preprint). In Review. 10.21203/rs.3.rs-2928984/v1

Prjibelski, A., Antipov, D., Meleshko, D., Lapidus, A., Korobeynikov, A., 2020. Using SPAdes De Novo Assembler. Current Protocols in Bioinformatics 70. 10.1002/cpbi.102

Ramesh, M.A., Malik, S.-B., Logsdon, J.M., 2005. A Phylogenomic Inventory of Meiotic Genes. Current Biology, 66 15, 185–191. 10.1016/j.cub.2005.01.003

Richards, T.A., Massana, R., Pagliara, S., Hall, N., 2019. Single cell ecology. Phil. Trans. R. Soc. B 374, 20190076. 10.1098/rstb.2019.0076

Richter, D.J., Berney, C., Strassert, J.F.H., Poh, Y.-P., Herman, E.K., Muñoz-Gómez, S.A., Wideman, J.G., Burki, F., de Vargas, C., 2022. EukProt: A database of genome-scale predicted proteins across the diversity of eukaryotes. Peer Community Journal, 64 2, e56. 10.24072/pcjournal.173

Rizos, I., Frada, M., Bittner L., Not, F., 2024. Life cycle strategies in free-living unicellular eukaryotes: diversity, evolution and current molecular tools to unravel the private life of microorganisms. J. Eukaryot. Microbiol. (in press)

Röttger, R., 1974. Larger foraminifera: Reproduction and early stages of development in Heterostegina depressa. Mar. Biol. 26, 5–12. 10.1007/BF00389081

Sandin, M., 2019. Diversité et Évolution des Nassellaires et Spumellaires (Radiolaires). Dissertation. Sorbonne Université. 202 p.

Schewiakoff, W.T., 1926. The Acantharia. Fauna e Flora del Golfo di Napoli 37: 1–755

Schmieder, R., Edwards, R., 2011. Quality control and preprocessing of metagenomic datasets. Bioinformatics 27, 863–864. 10.1093/bioinformatics/btr026

Schurko, A.M., Logsdon, J.M., 2008. Using a meiosis detection toolkit to investigate ancient asexual “scandals” and the evolution of sex. Bioessays, 11 30, 579–589. 10.1002/bies.20764

Speijer, D., 2016. What can we infer about the origin of sex in early eukaryotes? Phil. Trans. R. Soc. B, 8 371, 20150530. 10.1098/rstb.2015.0530

Speijer, D., Lukeš, J., Eliáš, M., 2015. Sex is a ubiquitous, ancient, and inherent attribute of eukaryotic life. Proc. Natl. Acad. Sci. U.S.A. 112, 8827–8834. 10.1073/pnas.1501725112

Suzuki, N., Not, F., 2015. Biology and Ecology of Radiolaria, in: Ohtsuka, S., Suzaki, T., Horiguchi, T., Suzuki, N., Not, F. (Eds.), Marine Protists. Springer Japan, Tokyo, pp. 179–222. 10.1007/978-4-431-55130-0_8

Tekle, Y.I., Wang, F., Heidari, A., Stewart, A.J., 2020. Differential gene expression analysis and cytological evidence reveal a sexual stage of an amoeba with multiparental cellular and nuclear fusion. PLoS ONE 15, e0235725. 10.1371/journal.pone.0235725

Thangavel, G., Hofstatter, P.G., Mercier, R., Marques, A., 2023. Tracing the evolution of the plant meiotic molecular machinery. Plant Reprod. 10.1007/s00497-022-00456-1

Ueda, M., Zhang, Z., Laux, T., 2011. Transcriptional Activation of Arabidopsis Axis Patterning Genes WOX8/9 Links Zygote Polarity to Embryo Development. Developmental Cell 20, 264–270. 10.1016/j.devcel.2011.01.009

Vicedomini, R., Blachon, C., Oteri, F., Carbone, A., 2021. MyCLADE: a multi-source domain annotation server for sequence functional exploration. Nucleic Acids Research 49, W452–W458. 10.1093/nar/gkab395

Villeneuve, A.M., Hillers, K.J., 2001. Whence Meiosis? Cell, 65 106, 647–650. 10.1016/S0092-8674(01)00500-1

Yuasa, T., Takahashi, O., 2016. Light and electron microscopic observations of the reproductive swarmer cells of nassellarian and spumellarian polycystines (Radiolaria). European Journal of Protistology 54, 19–32. 10.1016/j.ejop.2016.02.007

Yuasa, T., Takahashi, O., 2014. Ultrastructural morphology of the reproductive swarmers of Sphaerozoum punctatum (Huxley) from the East China Sea. European Journal of Protistology 50, 194–204. 10.1016/j.ejop.2013.12.001

