## Supplementary material for "Single-cell transcriptomics highlights sexual cues among reproductive life stages of uncultivated Acantharia (Radiolaria)": Suppl Figures

**Fig. S1. Overall gene expression of studied single-cell transcriptomes.** The value of gene expression is presented in log(TPM), while the number of predicted proteins in each transcriptome is indicated along the x axis.

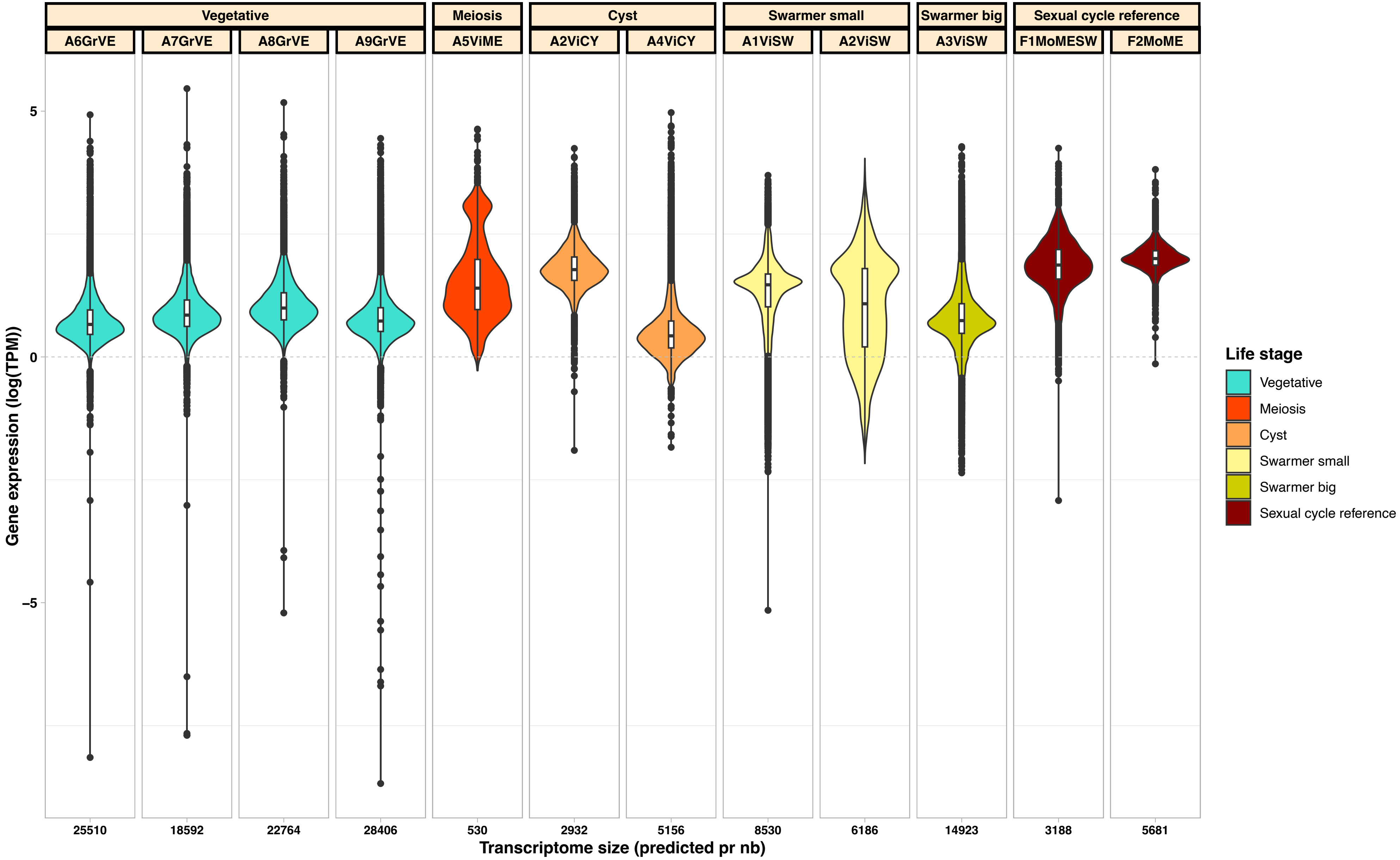

**Fig. S2. Relative proportion of annotated genes in single-cell transcriptomes.** Gene annotation is represented by three functionnal categories: match with HMM profile of sexual cycle genetic toolkit (red); match with functional COG in eggNOG database (grey); no functional annotation (black).

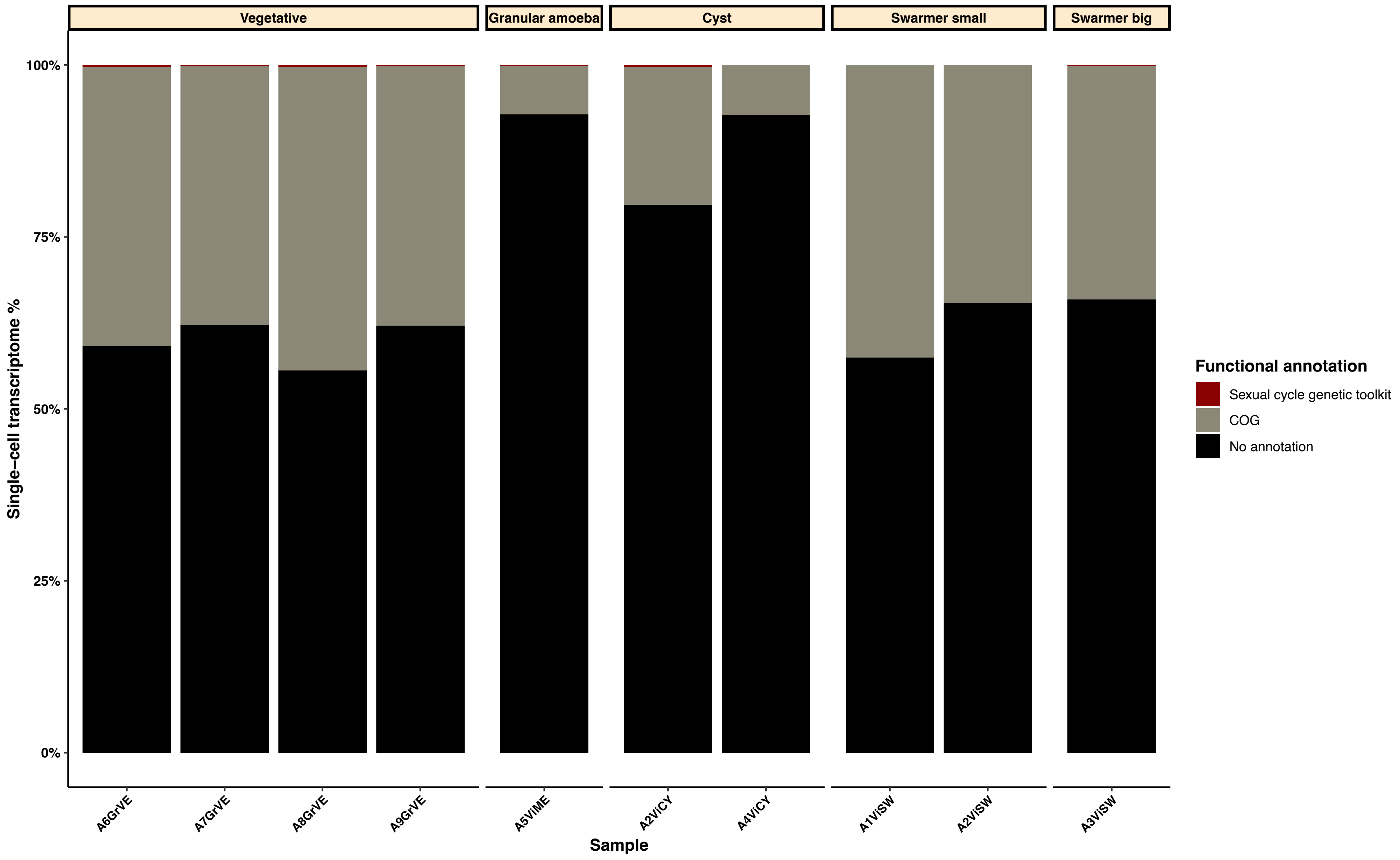

**Fig. S3. Functional annotation of transcripts in orthologous gene clusters.** The relative number of trnascripts corresponding to each functional cluster of orthologous genes (COGs) is shown for the identified life stage-specific OGs. Transcripts lacking COG annotation are indicated as "No COG" in light green.

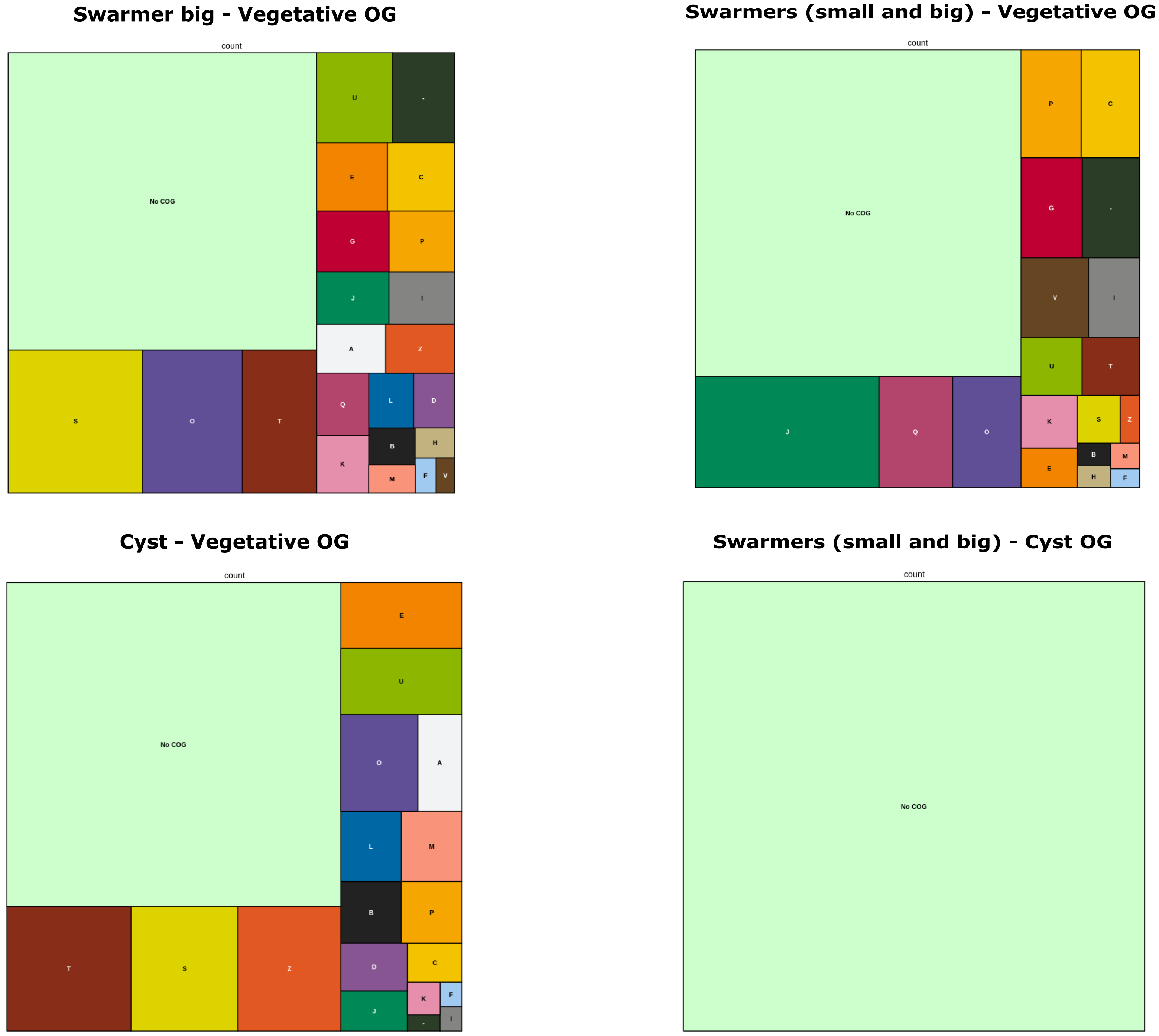

### All life stage OG

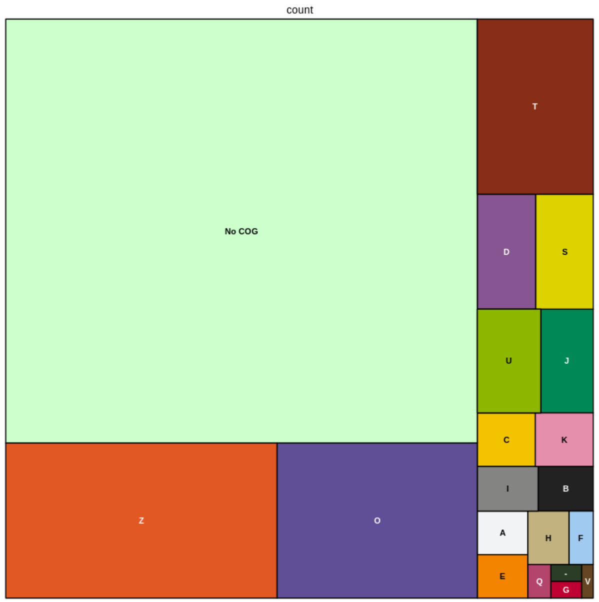


### Vegetative OG

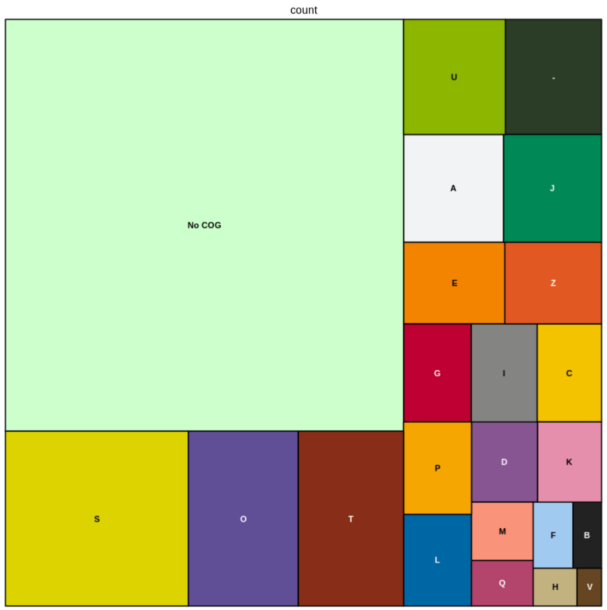


### Cyst OG

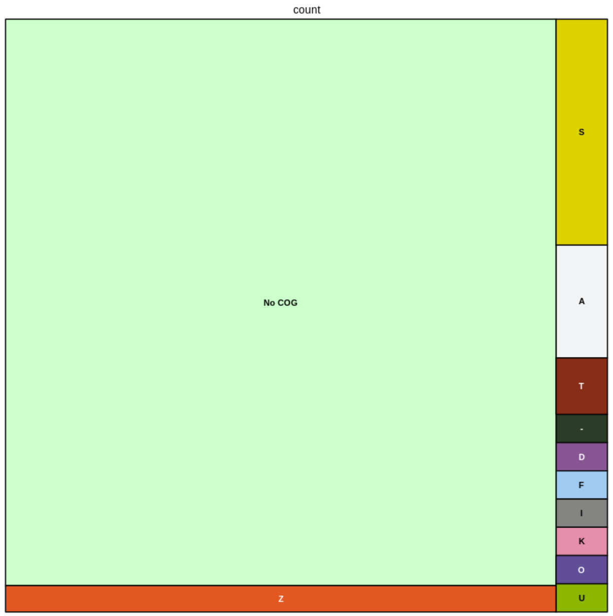


### Swarmers (small and big) OG

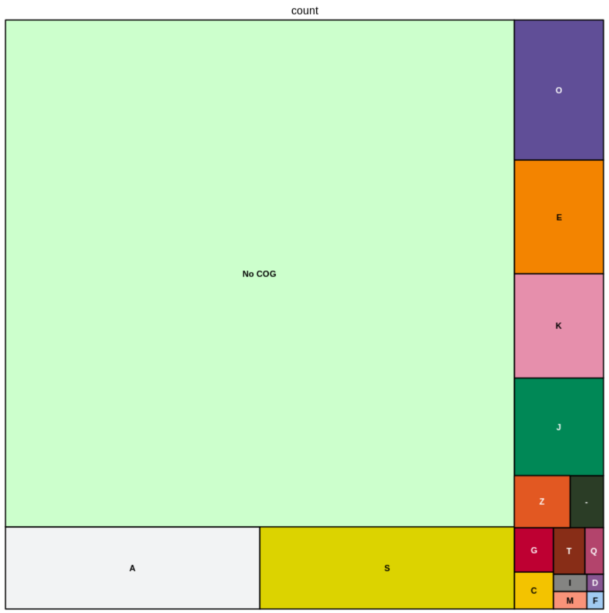


### Swarmers big OG

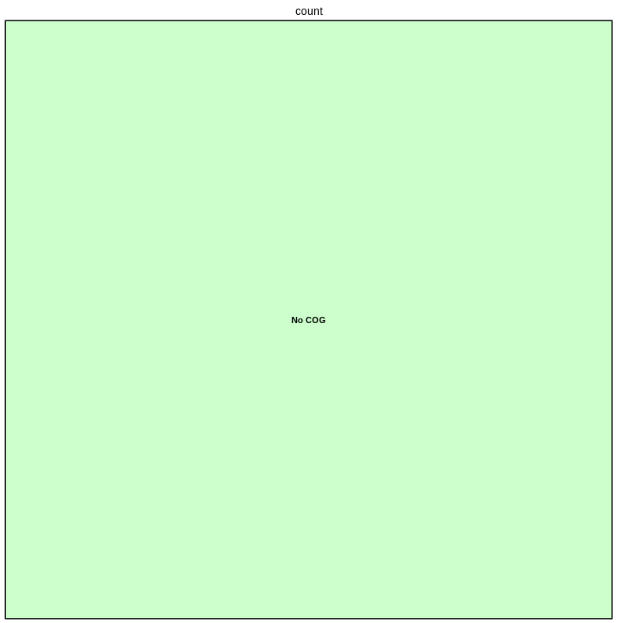

**Fig. S4. Top life stage-specific orthologous gene groups differentially expressed in single-cell transcriptomes.** The orthologous gene groups (OGs) were estimated by Orthofinder (cf. Materials and Methods). Predicted functional annotations of these groups are available in Table S3.

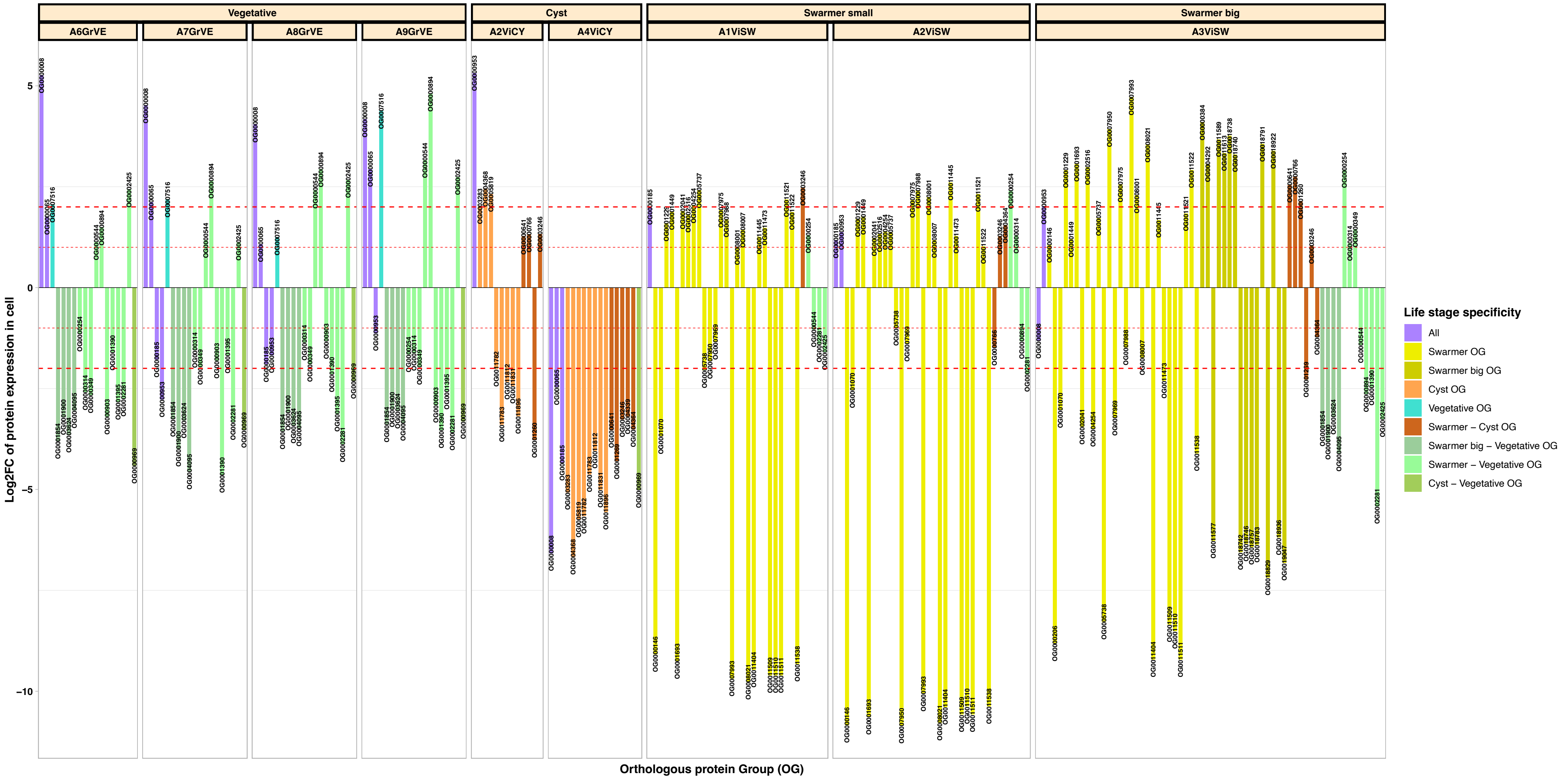

**Fig. S5. Relative expression of 18S and 28S ribosomal marker gene copies in each single-cell transcriptome.** For each transcriptome, the most expressed copy of the 18S ribosomal gene (top) and 28S ribosomal gene (bottom) were concatenated for the phylogenetic recostruction (cf. Materials and Methods).

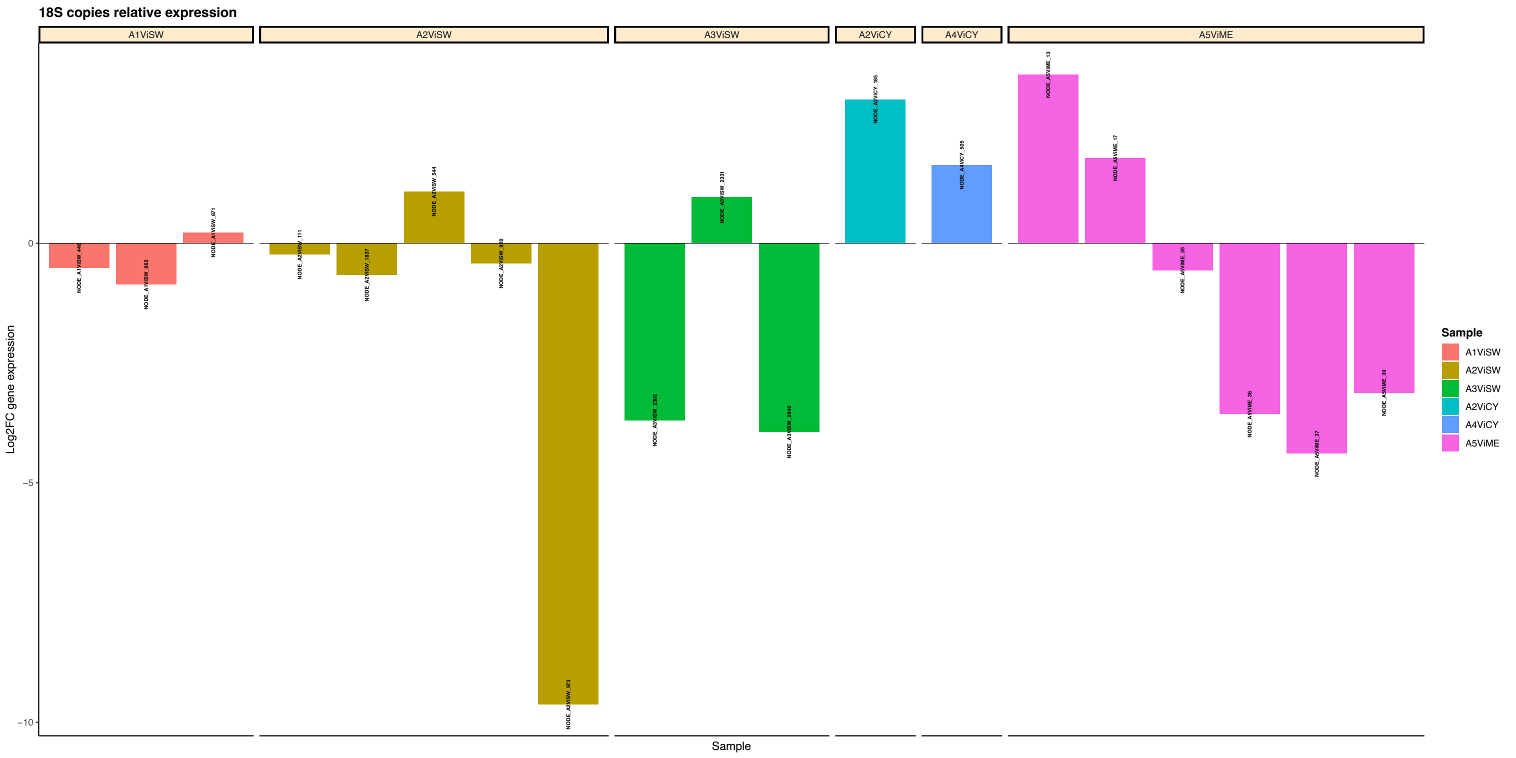


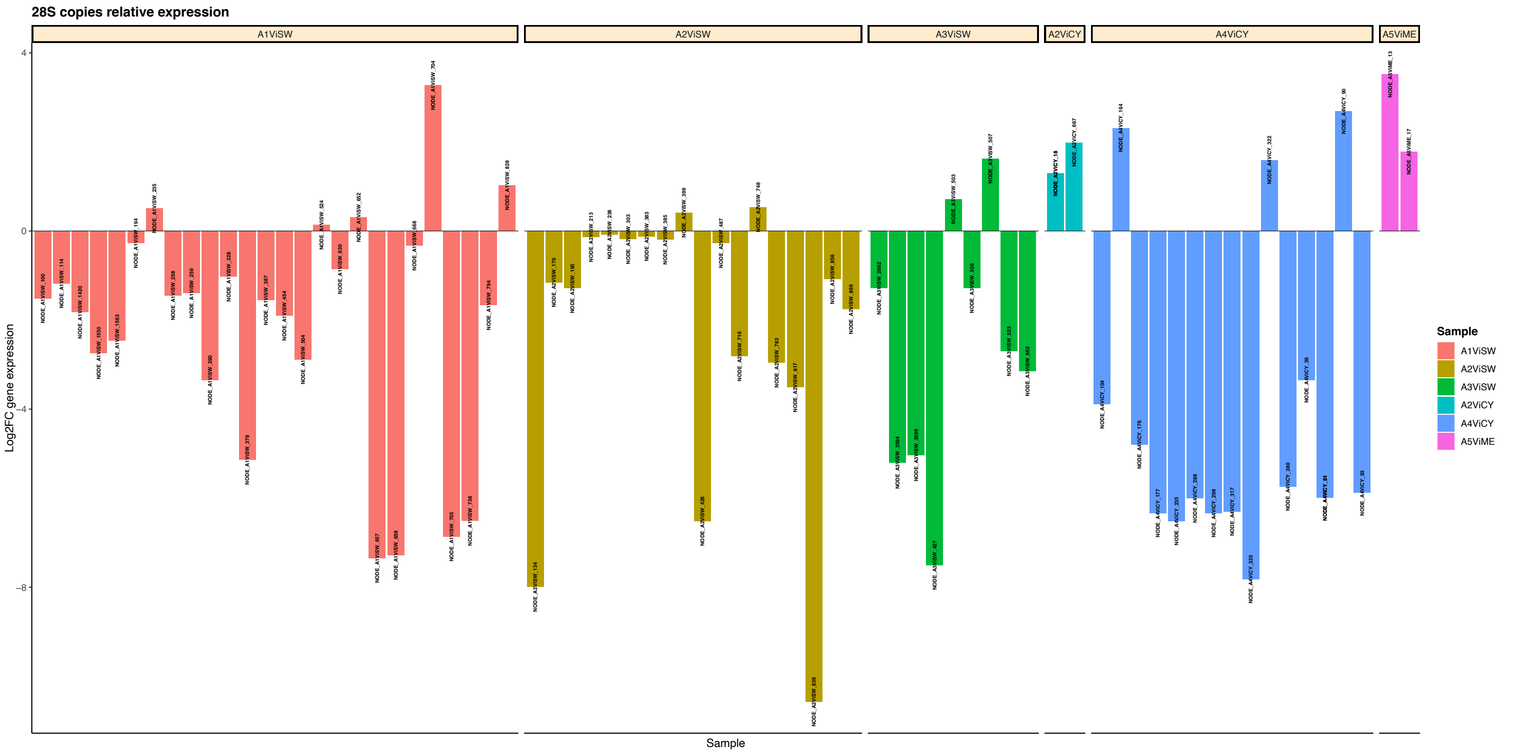

**Fig. S6. Most expressed life stage-specific orthologous genes groups differentially expressed in single-cell transcriptomes.** The expression of each orthologous gene group (OG) is indicated by range values of log2FC (i.e., group2). Each plot represents the OGs specific to one life stage (e.g., cysts) or multiple life stages (e.g.., swarmers and cysts).

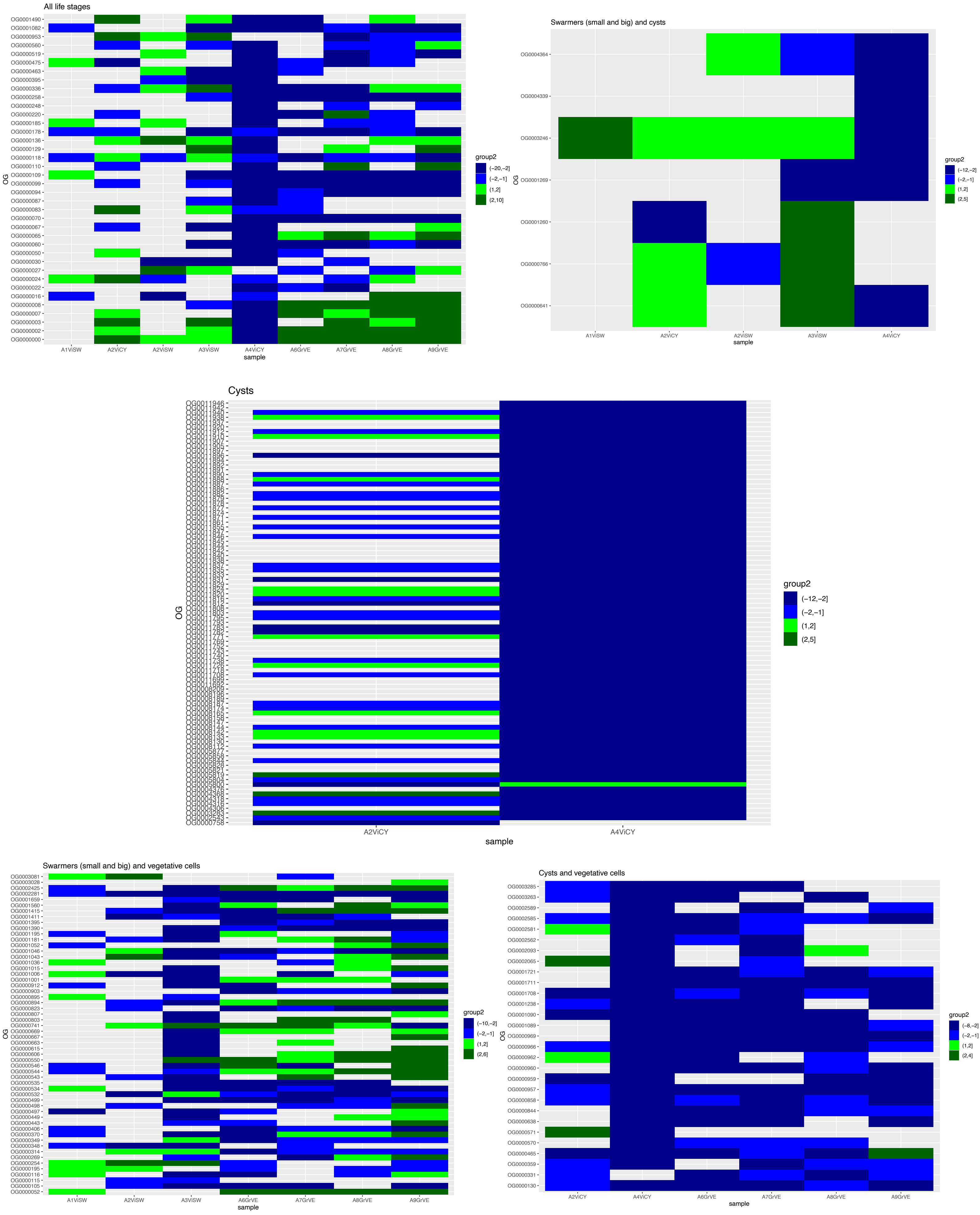

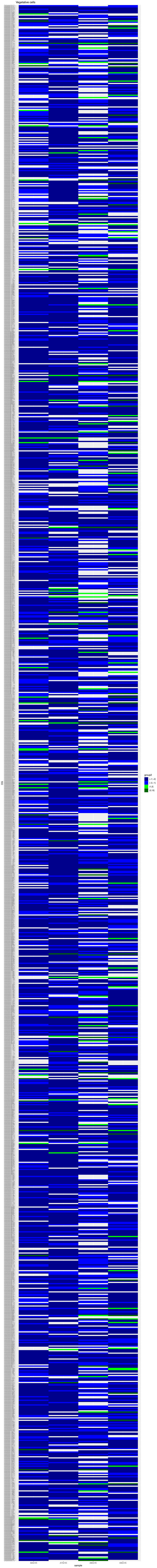
Vegetative cells
OG
A6GrVE
A7GrVE
A8GrVE
A9GrVE
sample
group2
(−7,−2]
(−2,−1]
(1,2]
(2,13]

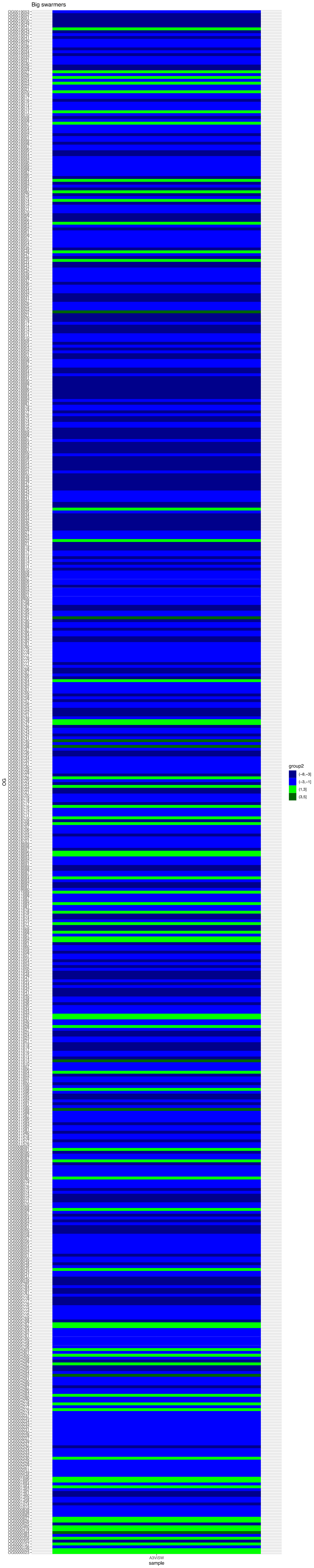
Big swarmers
OG
sample
A3VISW
group2
(-8,-3]
(-3,-1]
(1,3]
(3,5]

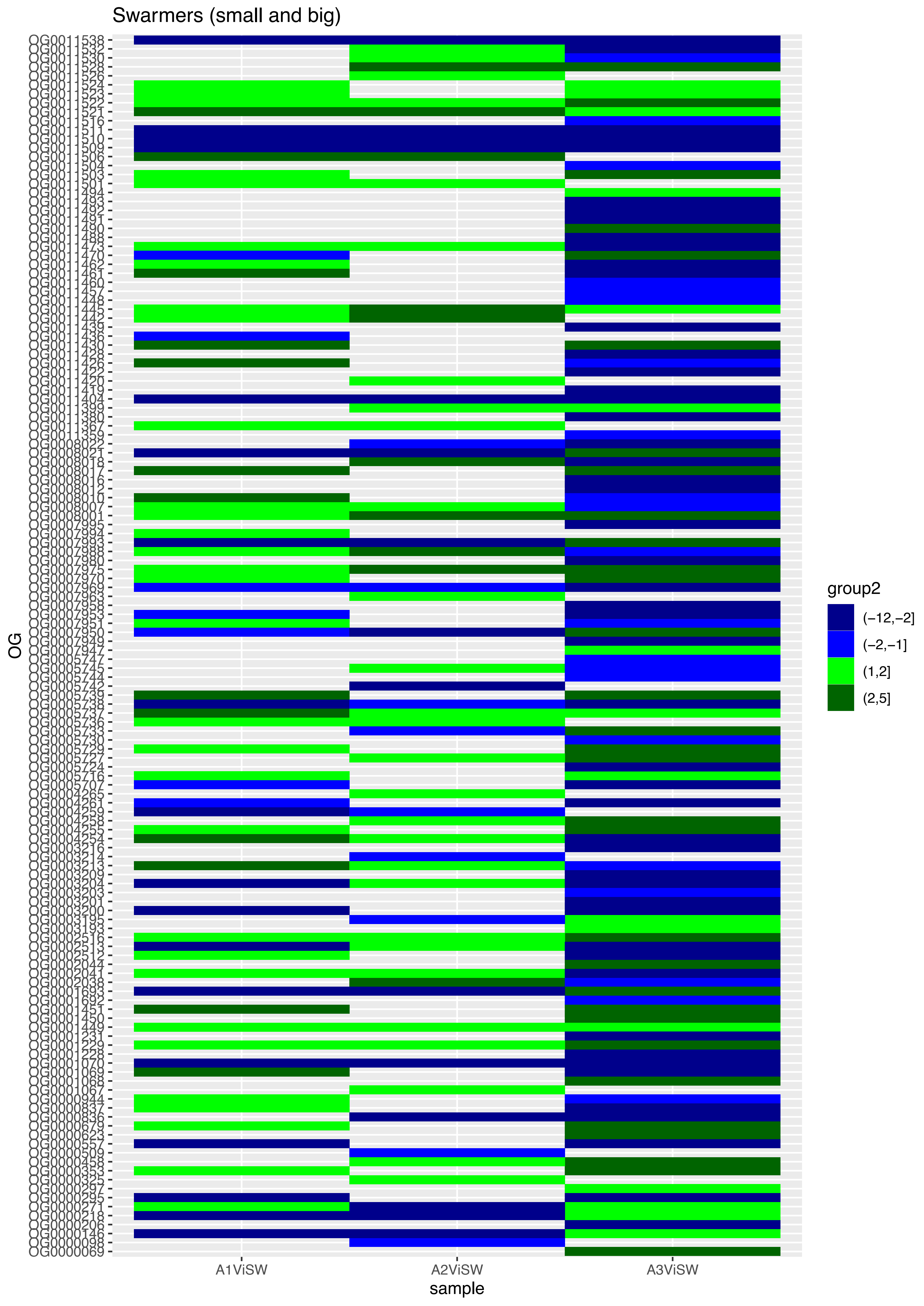
Swarmers (small and big)
OG
A1ViSW
A2ViSW
A3ViSW
sample
group2
(−12,−2]
(−2,−1]
(1,2]
(2,5]

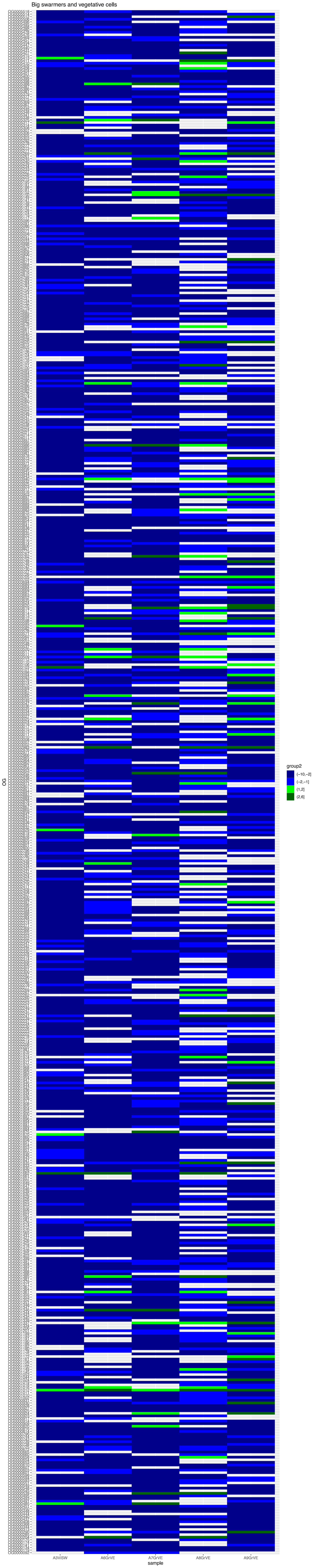

Big swarmers and vegetative cells
OG
A3ViSW
A6GrVE
A7GrVE
A8GrVE
A9GrVE
sample
group2
(−10,−2]
(−2,−1]
(1,2]
(2,6]
