## Supplementary material for "Single-cell transcriptomics highlights sexual cues among reproductive life stages of uncultivated Acantharia (Radiolaria)": Suppl Tables

**Table S2. List of query reference proteins related to meiosis and syngamy that were searched in single-cell transcriptomes.** The HMM profiles of these proteins were downloaded from PFAM (http://pfam.xfam.org/). The UniProt annotation score (https://www.uniprot.org/) is indicated as a proxy to the level of knowledge regarding each protein and illustrates overall syngamy-related proteins have lower UniProt scores compared to meiosis-related proteins. The bibliography in which the proteins were cited in included in the column Reference: complete bibliography in given only for papers that are absent from the main text. For references cites multiple times, detailed bibliography is provided at first occurence.

| MEIOSIS GENETIC TOOLKIT (gene id) | Protein name | Domain ID | PFAM | Function | Cellular process | Molecular process | Specificity | UniProt annotation score (protist sequences) | Reference |
|---|---|---|---|---|---|---|---|---|---|
| **REC8** | Recombination protein 8 | RAD21_REC8_N, RAD21_REC8 | PF04825, PF04824 | Sister chromatid cohesion | Meiosis | Cohesion complex | Meiosis specific | 5 | Hoftstatter et al 2018; Hofstatter & Lahr 2019; Malik et al 2008; Schurko, A.M., Logsdon, J.M., 2008. Using a meiosis detection toolkit to investigate ancient asexual "scandals" and the evolution of sex. Bioessays, 11 30, 579–589. https://doi.org/10.1002/bies.20764 |
| **HOP1** | Meiosis-specific protein HOP1 | HORMA | PF02301 | Synaptonemal complex (SC) protein involved in chromosome pairing during meiosis | Meiosis | Homologous alignment and synaptonemal complex | Meiosis specific | 5 | Hofstatter et al 2018; Hofstatter & Lahr 2019; Malik et al 2008; Ramesh et al 2005; Schurko & Logsdon 2008 |
| **SPO11** | Meiosis-specific protein SPO11 | TP6A_N | PF04406 | Creation of double strand breaks to initiation meiotic recombination | Meiosis | DSB | Meiosis specific | 5 | Hofstatter et al 2018; Hofstatter & Lahr 2019; Malik et al 2008; Ramesh et al 2005; Schurko & Logsdon 2008; Villeneuve, A.M., Hillers, K.J., 2001. Whence Meiosis? Cell, 65 106, 647–650. https://doi.org/10.1016/S0092-8674(01)00500-1 |
| **HOP2** | Homologous-pairing protein 2 | TBIP | PF07106 | Forms heterodimer with MND1 promoting interhomolog meiotic recombination | Meiosis | Recombination | Meiosis specific | 5 | Hofstatter et al 2018; Hofstatter & Lahr 2019; Malik et al 2008; Ramesh et al 2005; Schurko & Logsdon 2008 |
| **MND1** | Meiotic nuclear division protein 1 | MND1 | PF03962 | Forms heterodimer with HOP2 promoting interhomolog meiotic recombination | Meiosis | Recombination | Meiosis specific | 4 | Hofstatter et al 2018; Hofstatter & Lahr 2019; Malik et al 2008; Ramesh et al 2005; Schurko & Logsdon 2008 |
| **DMC1** | Meiotic recombination protein DMC1 | RAD51 | PF08423 | Meiosis-specific homolog of Rad51, has similar function but promotes interhomolog recombination | Meiosis | Recombination | Meiosis specific | 5 | Hofstatter et al 2018; Hofstatter & Lahr 2019; Malik et al 2008; Ramesh et al 2005; Schurko & Logsdon 2008; Villeneuve & Hillers 2001 |
| **MSH4** | MutS protein homolog 4 | MUTS domains: MUTS_III, MUTS_IV, MUTS_V | PF05192, PF05190, PF00488 | Meiosis specific MutS homologs functioning as a heterodimer in meiotic recombination and Holliday junction resolution | Meiosis | CO RESOLUTION | Meiosis specific | 5 | Hofstatter et al 2018; Hofstatter & Lahr 2019; Malik et al 2008; Ramesh et al 2005; Schurko & Logsdon 2008; Villeneuve & Hillers 2001 |
| **MSH5** | MutS protein homolog 5 | MUTS domains: MUTS_III, MUTS_IV, MUTS_V | PF05192, PF05190, PF00488 | Meiotic recombination | Meiosis | CO RESOLUTION | Meiosis specific | 4 | Hofstatter et al 2018; Hofstatter & Lahr 2019; Malik et al 2008; Ramesh et al 2005; Schurko & Logsdon 2008; Villeneuve & Hillers 2001 |
| **SCC3** | Sister-chromatid cohesion protein 3 | STAG | PF08514 | Necessary for sister chromatid cohesion and required for DSB repair | Meiosis | Cohesion complex | Meiosis/mitosis | 5 | Malik et al 2008 |
| **MSH2** | MutS protein homolog 2 | MUTS domains: MUTS_III, MUTS_IV, MUTS_V | PF05192, PF05190, PF00488 | Mismatch repair and promotion of meiotic crossing over | Meiosis | Mismatch repair | Meiosis/mitosis | 4 | Hofstatter et al 2018; Hofstatter & Lahr 2019; Malik et al 2008; Ramesh et al 2005; Schurko & Logsdon 2008; Villeneuve & Hillers 2001 |
| **MSH6** | MutS protein homolog 6 | MUTS domains: MUTS_III, MUTS_IV, MUTS_V | PF05192, PF05190, PF00488 | Mismatch repair and promotion of meiotic crossing over | Meiosis | Mismatch repair | Meiosis/mitosis | 5 | Hofstatter et al 2018; Hofstatter & Lahr 2019; Malik et al 2008; Ramesh et al 2005; Schurko & Logsdon 2008; Villeneuve & Hillers 2001 |
| **PDS5** | Sister chromatid cohesion protein pds5 | CND1 | PF12717 | Maintenance of sister chromatid cohesion in late prophase | Meiosis | Cohesion complex | Meiosis/mitosis | 5 | Malik et al 2008 |
| **SMC1** | Structural maintenance of chromosomes protein 1 | SMC_N, SMC_hinge | PF02463, PF06470 | This domain is found at the N terminus of SMC proteins. The SMC (structural maintenance of chromosomes) superfamily proteins have ATP-binding domains at the N- and C-termini, and two extended coiled-coil domains separated by a hinge in the middle. The eukaryotic SMC proteins form two kind of heterodimers: the SMC1/SMC3 and the SMC2/SMC4 types. These heterodimers constitute an essential part of higher order complexes, which are involved in chromatin and DNA dynamics | Meiosis | Cohesion complex | Meiosis/mitosis | 5 | Malik et al 2008 |
| **SMC3** | Structural maintenance of chromosomes protein 3 | SMC_N, SMC_hinge | PF02463, PF06470 | | Meiosis | Cohesion complex | Meiosis/mitosis | 5 | Malik et al 2008 |
| **MRE11** | Double-strand break repair protein MRE11 | MRE11_DNA_bind | PF04152 | 3′-5′ dsDNA exonuclease and ssDNA endonuclease; forms complex with Rad50 and Xrs2/Nbs1 | Meiosis | DSB | Meiosis/mitosis | 5 | Malik et al 2008; Ramesh et al 2005; Villeneuve & Hillers 2001 |
| **RAD51** | DNA repair protein RAD51 homolog 1 | RAD51 | PF08423 | Forms helical filaments on single-stranded and double-stranded DNA and catalyzes homologous DNA pairing and strand exchange. (Intrahomologous recombination) | Meiosis | Recombination | Meiosis/mitosis | 5 | Hofstatter et al 2018; Malik et al 2008 |
| **RAD1** | DNA repair protein RAD1 | RAD1 | PF02144 | Forms a heterodimer with Rad10 (Ercc1) Required for meiotic crossing over, normal meiotic chromosome disjunction, to repair mismatches in heteroduplex DNA and to resolve reciprocally exchanged recombination intermediates in Drosophila | Meiosis | CO RESOLUTION | Meiosis/mitosis | 5 | Malik et al 2008 |
| **RAD50** | DNA repair protein RAD50 | RAD50_zn_hook | PF04423 | ATPase, DNA binding protein; in a complex with Mre11/Xrs2, holds broken DNA ends together while Mre11 trims | Meiosis | DSB | Meiosis/mitosis | 5 | Malik et al 2008; Ramesh et al 2005 |
| **RAD52** | DNA repair protein RAD52 | RAD52_RAD22 | PF04098 | Binds DSBs and initiates assembly of meiotic recombination complexes | Meiosis | DSB | Meiosis/mitosis | 5 | Malik et al 2008; Ramesh et al 2005 |
| **MLH1** | DNA mismatch repair protein Mlh1 | MLH1_C | PF16413 | Mismatch repair and promotion of meiotic crossing over; interacts with Msh2/Msh6 and Msh4/Msh5; forms heterodimers with Mlh2, Mlh3, and Pms1 | Meiosis | CO RESOLUTION | Meiosis/mitosis | 5 | Malik et al 2008; Ramesh et al 2005; Villeneuve & Hillers 2001 |
| **MUS81** | Crossover junction endonuclease MUS81 | ERCC4 | PF02732 | This domain is a family of nucleases. The family includes EME1 which is an essential component of a Holliday junction resolvase [2-3]. EME1 interacts with MUS81 to form a DNA structure-specific endonuclease.Structure-specific DNA repair endonuclease responsible for the 5-prime incision during DNA repair. Involved in homologous recombination that assists in removing interstrand cross-link Interacts with MMS4 to form a DNA structure-specific endonuclease with substrate preference for branched DNA structures with a 5'-end at the branch nick. | Meiosis | CO RESOLUTION | Meiosis/mitosis | 5 | Hofstatter & Lahr 2019 |
| **MER3** | DExH-box ATP-dependent RNA helicase DExH17 | SEC63 | PF02889 | Meiosis-specific DEAD-box helicase. Promotes Holliday junction resolution together with proteins including MSH4 and MSH5 | Meiosis | CO RESOLUTION | Meiosis/mitosis | 5 | Hofstatter et al 2018; Hofstatter & Lahr 2019; Malik et al 2008 |
| **PMS1** | DNA mismatch repair protein PMS1 | MUTL_C, DNA_mismatch_repair | PF08676, PF01119 | Forms heterodimer with Mlh1 for repair of heteroduplex DNA; interacts with Msh2/Msh3 | Meiosis | Mismatch repair | Meiosis/mitosis | 5 | Malik et al 2008; Ramesh et al 2005 |
| **PMS2** | Mismatch repair endonuclease PMS2 | MUTL_C | PF08676 | Component of the post-replicative DNA mismatch repair system (MMR). Heterodimerizes with MLH1 to form MutL alpha. DNA repair is initiated by MutS alpha (MSH2-MSH6) or MutS beta (MSH2-MSH3) binding to a dsDNA mismatch, | Meiosis | Mismatch repair | Meiosis/mitosis | 5 (mouse sequence) | Hoftstatter et al 2018 |
| **MLH2** | DNA mismatch repair protein MLH2 | DNA_mis_repair | PF01119 | Forms a heterodimer with Mlh1. Interacts with Msh2/3 or Msh2/6 for removal of cisplatin adducts | Meiosis | Mismatch repair | Meiosis/mitosis | 4 | Malik et al 2008; Ramesh et al 2005 |
| **SPO22** | Sporulation-specific protein 22 | SPO22/ZIP4 | PF08631 | SPO22/ZIP4 in yeast is a meiosis specific protein involved in sporulation. It has been shown to regulate crossover distribution by promoting synaptonemal complex formation | Meiosis | Homologous alignment and synaptonemal complex | Meiosis/mitosis | 3 | Hoftstatter et al 2018; Hofstatter & Lahr 2019 |
| **EXO1** | Exonuclease 1 | XPG_I, XPG_N | PF00867, PF00752 | 5'->3' double-stranded DNA exonuclease that could act in a pathway that corrects mismatched base pairs | Meiosis | CO RESOLUTION | Meiosis/mitosis | 5 | Hoftstatter et al 2018 |
| **SMC2** | Structural maintenance of chromosomes protein 2 | SMC_N, SMC_hinge | PF02463, PF06470 | Forms a heterodimer with Smc4 to form core condensin subunits, ring shape, essential for chromosome assembly and segregation. | Meiosis | Cohesion complex | Meiosis/mitosis | 5 | Malik et al 2008 |
| **SMC4** | Structural maintenance of chromosomes protein 4 | SMC_N, SMC_hinge | PF02463, PF06470 | Forms a heterodimer with Smc2 to form core condensin subunits, ring shape, essential for chromosome assembly and segregation | Meiosis | Cohesion complex | Meiosis/mitosis | 5 | Malik et al 2008 |
| **SMC5** | Structural maintenance of chromosomes protein 5 | SMC_N | PF02463 | Forms a heterodimer with Smc6 (Rad18) and is involved in DNA repair and checkpoint responses | Meiosis | Homologous alignment and synaptonemal complex | Meiosis/mitosis | 5 | Malik et al 2008 |
| **SCC1** | Sister chromatid cohesion protein 1 | RAD21_REC8_N, RAD21_REC8 | PF04825, PF04824 | Holds Smc1 and Smc3, holds sister chromatids together during mitosis and meiosis | Meiosis | Cohesion complex | Meiosis/mitosis | 5 | Malik et al 2008 |
| **LZ3wCH** | Leucine zipper with capping helix domain | LZ3wCH | PF18517 | Interaction with HOP2 and MND1 | Meiosis | Recombination | Meiosis/mitosis | 4 | Schwelm, A., Fogelqvist, J., Knaust, A., Jülke, S., Lilja, T., Bonilla-Rosso, G., Karlsson, M., Shevchenko, A., Dhandapani, V., Choi, S.R., Kim, H.G., Park, J.Y., Lim, Y.P., Ludwig-Müller, J., Dixelius, C., 2015. The Plasmodiophora brassicae genome reveals insights in its life cycle and ancestry of chitin synthases. Sci Rep 5, 11153. https://doi.org/10.1038/srep11153 |
| **RAD54** | DNA repair and recombination protein RAD54 | RAD54_N, SNF2-rel_dom | PF08658, PF00176 | Loading of RAD51/DMC1, stand invasion, CO, mismatch repair | Meiosis | CO RESOLUTION | Meiosis/mitosis | 5 | Worden, A.Z., Lee, J.-H., Mock, T., Rouzé, P., Simmons, M.P., Aerts, A.L., Allen, A.E., Cuvelier, M.L., Derelle, E., Everett, M.V., Foulon, E., Grimwood, J., Gundlach, H., Henrissat, B., Napoli, C., McDonald, S.M., Parker, M.S., Rombauts, S., Salamov, A., Von Dassow, P., Badger, J.H., Coutinho, P.M., Demir, E., Dubchak, I., Gentemann, C., Eikrem, W., Gready, J.E., John, U., Lanier, W., Lindquist, E.A., Lucas, S., Mayer, K.F.X., Moreau, H., Not, F., Otillar, R., Panaud, O., Pangilinan, J., Paulsen, I., Piegu, B., Poliakov, A., Robbens, S., Schmutz, J., Toulza, E., Wyss, T., Zelensky, A., Zhou, K., Armbrust, E.V., Bhattacharya, D., Goodenough, U.W., Van de Peer, Y., Grigoriev, I.V., 2009. Green Evolution and Dynamic Adaptations Revealed by Genomes of the Marine Picoeukaryotes *Micromonas*. Science 324, 268–272. https://doi.org/10.1126/science.1167222 |
| **NBS1** | Nibrin homolog | FHA | PF00498 | Participates to formation of Mre11 complex, telomere maintenance | Meiosis | DSB | Meiosis/mitosis | - | Ueno, M., Nakazaki, T., Akamatsu, Y., Watanabe, K., Tomita, K., Lindsay, H.D., Shinagawa, H., Iwasaki, H., 2003. Molecular Characterization of the *Schizosaccharomyces pombe nbs1 +* Gene Involved in DNA Repair and Telomere Maintenance. Molecular and Cellular Biology 23, 6553–6563. https://doi.org/10.1128/MCB.23.18.6553-6563.2003 |
| **MNS1 (YEAST)** | Meiosis-specific nuclear structural protein 1 | GLYCO_HYDRO_47 | PF01532 | Involved in organisation of the nuclear or perinuclear architecture and may affect the nuclear morphology during meiotic prophase in spermatocytes | Meiosis | Structural | Meiosis/mitosis | 5 | Grishaeva, T.M., Bogdanov, Y.F., 2018. Conservation of meiosis-specific nuclear proteins in eukaryotes: a comparative approach. Nucleus 61, 175–182. https://doi.org/10.1007/s13237-018-0253-8 |
| **MNS1 (Rhizaria)** | Meiosis-specific nuclear structural protein 1 | TPH | PF13868 | Involved in organisation of the nuclear or perinuclear architecture and may affect the nuclear morphology during meiotic prophase in spermatocytes | Meiosis | Structural | Meiosis/mitosis | 2 | Grishaeva, T.M., Bogdanov, Y.F., 2018. Conservation of meiosis-specific nuclear proteins in eukaryotes: a comparative approach. Nucleus 61, 175–182. https://doi.org/10.1007/s13237-018-0253-8 |
| **RED1** | Protein RED1 | REC10_RED1 | PF07964 | Rec10 / Red1 is involved in meiotic recombination and chromosome segregation during homologous chromosome formation | Meiosis | Recombination | Meiosis/mitosis | 5 | Shah, S., Chen, Y., Bhattacharya, D., Chan, C.X., 2020. Sex in Symbiodiniaceae dinoflagellates: genomic evidence for independent loss of the canonical synaptonemal complex. Sci Rep 10, 9792. https://doi.org/10.1038/s41598-020-66429-4 |
| **SYNGAMY GENETIC TOOLKIT (gene id)** | **Protein name** | **Domain ID** | **PFAM** | **Function** | **Cellular process** | **Molecular process** | **Specificity** | **Uniprot annotation score (protist sequences)** | **Reference** |
| **FUS1** | Nuclear fusion protein FUS1 | SH3_1 (not function specific) | PF00018 | Required for cell fusion, induced by incubation of haploid chlamydomonas + cells with mating pheromone | Syngamy | Cell-cell adhesion | Gamete specific | 5 | Hernandez et al 2017; Ning et al 2013 |
| **HAP2/GCS1** | Hapless 2 / Generative Cell Specific-1 | HAP2-GSC1 | PF10699 | Considered ancestral gamete fusogan, required for membrane fusion in protists, flowering plants and some invertebrates | Syngamy | Plasmogamy | Gamete specific | 5 | Hofstatter & Lahr 2019 |
| **FIG1** | Factor-induced gene 1 protein | FIG1 | PF12351 | During the mating process of yeast cells, two Ca2+ influx pathways become activated. The resulting elevation of cytosolic free Ca2+ activates downstream signaling factors that promote long term survival of unmated cells. Fig1 is a regulator of the low affinity Ca2+ influx system (LACS), and is also required for efficient membrane fusion during yeast mating. | Syngamy | Plasmogamy | Gamete specific | 4 | Tekle et al 2020 |
| **KAR5** | Nuclear fusion protein KAR5 | Tht1 | PF04163 | Nuclear fusion protein KAR5 is an integral membrane protein that is thought to be required for the fusion of nuclear envelopes during karyogamy in yeasts | Syngamy | Karyogamy | Gamete specific | 5 | Ning et al 2013 |
| **FUS2** | Nuclear fusion protein FUS2 | RhoGEF (not function specific domain) | PF00621 | Involved in pore expansion during plasmogamy in yeasts | Syngamy | Plasmogamy | Gamete specific | 4 | Merlini, L., Dudin, O., Martin, S.G., 2013. Mate and fuse: how yeast cells do it. Open Biol. 3, 130008. https://doi.org/10.1098/rsob.130008 |
| **KAR4** | Karyogamy protein KAR4 | MT-A70 (not function specific) | PF05063 | May assist STE12 in the pheromone-dependent expression of KAR3 and CIK1. Also required for meiosis in yeasts | Syngamy | Karyogamy | Gamete specific | 4 | Tekle et al 2020 |
| **GSM1** | Gamete-specific minus 1 | Homeobox_KN (not function specific) | PF05920 | Chlamydomonas, - cells | Syngamy | Zygote development | Gamete specific | 1 | Ning et al 2013 |
| **GSP1** | Gamete-specific homeodomain protein | Homeobox_KN (not function specific) | PF05920 | Chlamydomonas, + cells | Syngamy | Zygote development | Gamete specific | 1 | Ning et al 2013 |
| **CFA20** | Cilia- and flagella-associated protein 20 | CFA20 | PF05018 | Cilia and flagella associated protein | Syngamy | Flagellar protein | - | 5 | Merchant, S.S., Prochnik, S.E., Vallon, O., Harris, E.H., Karpowicz, S.J., Witman, G.B., Terry, A., Salamov, A., Fritz-Laylin, L.K., Maréchal-Drouard, L., Marshall, W.F., Qu, L.-H., Nelson, D.R., Sanderfoot, A.A., Spalding, M.H., Kapitonov, V.V., Ren, Q., Ferris, P., Lindquist, E., Shapiro, H., Lucas, S.M., Grimwood, J., Schmutz, J., Cardol, P., Cerutti, H., Chanfreau, G., Chen, C.-L., Cognat, V., Croft, M.T., Dent, R., Dutcher, S., Fernández, E., Fukuzawa, H., González-Ballester, D., González-Halphen, D., Hallmann, A., Hanikenne, M., Hippler, M., Inwood, W., Jabbari, K., Kalanon, M., Kuras, R., Lefebvre, P.A., Lemaire, S.D., Lobanov, A.V., Lohr, M., Manuell, A., Meier, I., Mets, L., Mittag, M., Mittelmeier, T., Moroney, J.V., Moseley, J., Napoli, C., Nedelcu, A.M., Niyogi, K., Novoselov, S.V., Paulsen, I.T., Pazour, G., Purton, S., Ral, J.-P., Riaño-Pachón, D.M., Riekhof, W., Rymarquis, L., Schroda, M., Stern, D., Umen, J., Willows, R., Wilson, N., Zimmer, S.L., Allmer, J., Balk, J., Bisova, K., Chen, C.-J., Elias, M., Gendler, K., Hauser, C., Lamb, M.R., Ledford, H., Long, J.C., Minagawa, J., Page, M.D., Pan, J., Pootakham, W., Roje, S., Rose, A., Stahlberg, E., Terauchi, A.M., Yang, P., Ball, S., Bowler, C., Dieckmann, C.L., Gladyshev, V.N., Green, P., Jorgensen, R., Mayfield, S., Mueller-Roeber, B., Rajamani, S., Sayre, R.T., Brokstein, P., Dubchak, I., Goodstein, D., Hornick, L., Huang, Y.W., Jhaveri, J., Luo, Y., Martínez, D., Ngau, W.C.A., Otillar, B., Poliakov, A., Porter, A., Szajkowski, L., Werner, G., Zhou, K., Grigoriev, I.V., Rokhsar, D.S., Grossman, A.R., 2007. The *Chlamydomonas* Genome Reveals the Evolution of Key Animal and Plant Functions. Science 318, 245–250. https://doi.org/10.1126/science.1143609 |
| **GEX2** | Protein GAMETE EXPRESSED 2 | Filamin (not function specific) | PF00630 | Involved in gamete adhesion in flowering plants | Syngamy | Cell-cell adhesion | Gamete specific | 2 | Brukman, N.G., Uygur, B., Podbilewicz, B., Chernomordik, L.V., 2019. How cells fuse. Journal of Cell Biology, 72 218, 1436–1451. https://doi.org/10.1083/jcb.201901017 |

**Table S3. List of most expressed life stage-specific orthologous gene groups differentially expressed in single-cell transcriptomes and their predicted annotation.**

| OG | Life stage specificity | Size (sequence nb) | BlastP (eval) | InterPro | MyCLADE (eval<1E-06) | Phyre2 (cov %, conf %) | Potential role in life cycle | Expression trend (\|log2FC\| > 1-fold) |
|---|---|---|---|---|---|---|---|---|
| OG0000953 | All | 13 | XP_003074738.1, Histone core [Ostreococcus tauri] (2E-65) | Histone H2A (IPR002119) | PF00125 - Core histone H2A/H2B/H3/H4 ; PF16211 - C-terminus of histone H2A | structural protein/dna, histone h2a.z (77, 100) | Histone | Vegetative cells (A7GrVE, A8GrVE, A9GrVE): down-regulated, small swarmers (A2ViSW): up-regulated, cysts (A2ViCY): up-regulated |
| OG0000185 | All | 33 | ETO35540.1, mitogen-activated protein kinase 2 [Reticulomyxa filosa] (5E-56) | protein phosphorylation (GO:0006468) ; protein kinase activity (GO:0004672) ; ATP binding (GO:0005524) | PF00069 - Protein kinase domain ; PF15663 - Zinc-finger containing family | transferase, mitogen-activated protein kinase 4 (91, 99.7) | Signalling | Vegetative cells (A7GrVE, A8GrVE): down-regulated, small swarmers: up-regulated, big swarmers: \|log2FC\| < 1-fold |
| OG0000008 | All | 162 | - | Actin family (IPR004000) | PF00022 - Actin ; PF13833 - EF-hand domain pair, Ca binding domain ; PF00036 - EF hand ; PF08070 - DTHCT (NUC029) region | contractile protein, skeletal muscle, actin (99, 100) | Cytoskeleton movements / Signalling | Vegetative cells: up-regulated, small swarmers: \|log2FC\| < 1-fold, big swarmers: down-regulated |
| OG0000065 | All | 57 | ETO34930.1, hypothetical protein RFI_02145 [Reticulomyxa filosa] (8E-105) | Small GTPase superfamily, ARF/SAR type (IPR006689) ; Small GTPase superfamily, ARF type (IPR024156) ; P-loop containing nucleoside triphosphate hydrolase (IPR027417) | PF00025 - ADP-ribosylation factor family ; PF03029 - Conserved hypothetical ATP binding protein ; PF09439 - Signal recognition particle receptor beta subunit | P-loop containing nucleoside triphosphate hydrolases (98, 99.9) | Signalling | Vegetative cells: up-reguled, other life stages: \|log2FC\| < 1-fold |
| OG0002516 | Swarmer | 8 | - | - | **PF04350 - Pilus assembly protein, PilO** | hydrolase, proteasome activator complex (16, 39) | **Cell fusion related protein** | up-regulated |
| OG0005737 | Swarmer | 11 | - | Coil | **PF10277 - Frag1/DRAM/Sfk1 family, conserved membrane related proteins ; PF07695 - trasnmembrane region of intracellular receptors** | hydrolase (32, 43) | **Membrane related protein** | up-regulated |
| OG0008001 | Swarmer | 4 | - | Coil | **PF05384 - sensor protein** DegS control of transition from exponential to stationary growth ; **PF03148 - Tektin cytoskeletal proteins** | membrane protein, merozoite (asexual form) protein 1 (msp-1) (75, 45) | **Cell cycle control** | up-regulated |
| OG0011445 | Swarmer | 3 | XP_059452423.1, RING-H2 finger protein ATL54-like [Corylus avellana] (2E-11) | Zinc finger, RING-type (IPR001841) ; Zinc finger, RING/FYVE/PHD-type (IPR013083) | **PF09729 - Gti1/Pac2 family, controls sexual development** of *Schizosaccharomyces pombe* ; **PF12861 - Anaphase-promoting complex subunit** 11 RING-H2 finger (Apc11) ; PF11793 - FANCL C-terminal domain, ubiquitin ligase subunit | Cytosolic protein, ring finger and chy finger domain-containing protein (46, 99.1) | **Sexual cycle control** | up-regulated |
| OG0011522 | Swarmer | 3 | ETO30159.1, hypothetical protein RFI_06961 [Reticulomyxa filosa] (2E-26) | Coil | **PF14197 - Centrosome localisation domain** of PPC89 | hydrolase (30, 77) | **Cell division** | up-regulated |
| OG0011521 | Swarmer | 3 | - | EF-Hand 1, calcium-binding site (IPR018247) ; EF-hand domain pair (IPR011992) | PF13499 - EF-hand domain pair ; PF08976 - EF-hand domain, this domain is found predominantly in DJ binding proteins | metal binding protein, crystal structure of calaxin with calcium (99, 78) | Signalling | up-regulated |
| OG0007975 | Swarmer | 4 | - | - | - | membrane protein, crystal structure of the pleckstrin homology domain of human sin1, a2 torc2 subunit (17, 46) | Unknown | up-regulated |
| OG0001449 | Swarmer | 11 | - | Coil | - | cell adhesion (35, 19.4) | Unknown | up-regulated |
| OG0001229 | Swarmer | 12 | KAH8099125.1, hypothetical protein JL720_2105 [Aureococcus anophagefferens] (2E-59) | IPR029032 AhpD-like (IPR029032) ; Alkylhydroperoxidase AhpD core (IPR004675) | PF02627 - Carboxymuconolactone decarboxylase family, Carboxymuconolactone decarboxylase (CMD) | cell adhesion (23, 18) | Unknown | up-regulated |
| OG0011538 | Swarmer | 3 | - | EF-hand domain pair (IPR011992) | PF04116 - Fatty acid hydroxylase ; PF00036 - EF hand ; PF13833 - EF-hand domain pair ; PF13405 - EF-hand domain | dna binding protein, ure3-bp sequence specific dna binding protein (71, 93) | Signalling | down-regulated |
| OG0011511 | Swarmer | 3 | WP_000084700.1, scaffolding protein D [Bacteria] (4E-107) | Scaffold protein D (IPR004196) | PF02925 - Bacteriophage scaffolding protein D | Scaffolding protein gpD of bacteriophage procapsid (91, 100) | Signalling | down-regulated |
| OG0011510 | Swarmer | 3 | WP_000065572.1, major capsid protein [Bacteria] (0) | Microviridae F protein (IPR003514), Capsid/spike protein, ssDNA virus (IPR016184) | PF02305 - Capsid protein (F protein) | Nucleoplasmin-like/VP (viral coat and capsid proteins), Microviridae-like VP (100, 100) | Unknown | down-regulated |
| OG0011509 | Swarmer | 3 | CAB1112048.1, unnamed protein product [Ectocarpus sp. CCAP 1310/34] (0) | **DNA replication (GO:0006260)** | **PF05840 - Bacteriophage replication gene A protein (GPA)** | Recombination, helitron transposase bound to Its (30, 70) | DNA replication | down-regulated |
| OG0011404 | Swarmer | 3 | KLS36010.1, Minor spike protein [Neisseria gonorrhoeae MIA_2011_03-09] (0) | Minor spike protein (IPR006777) | PF04687 - Microvirus H protein (pilot protein) | Viral protein, bacteriophage phix174 h protein residues 143-282 (100, 100) | Unknown | down-regulated |
| OG0007969 | Swarmer | 4 | - | - | PF02957 - TT viral ORF2 | Virus, core protein p3 (26, 55.4) | Unknown | down-regulated |
| OG0005738 | Swarmer | 5 | - | Armadillo-type fold (IPR016024) ; Armadillo-like helical (IPR011989) | PF04826 - Armadillo-like ; PF05804 - Kinesin-associated protein (KAP) | Hydrolase, nad(+) hydrolase tir-1 (98, 91.2) | Signalling | down-regulated |
| OG0001070 | Swarmer | 13 | - | Coil | PF08614 - Autophagy protein 16 (ATG16) ; PF01576 - Myosin tail ; **PF05557 - Mitotic checkpoint protein** ; PF10174 - RIM-binding protein of the cytomatrix active zone ; PF05701 - Weak chloroplast movement under blue light | Structural protein, 'flagella associated protein' (98, 98.9) | Cell cycle control | down-regulated |
| OG0008021 | Swarmer | 4 | XP_003293760.1, uncharacterized protein DICPUDRAFT_99749 [Dictyostelium purpureum] (1E-05) | nucleic acid binding (GO:0003676) ; RNA binding (GO:0003723) | PF13893 - RNA recognition motif. (a.k.a. RRM, RBD, or RNP domain) | splicing, rna-binding protein (50, 93.8) | Post-transcriptional modifications | Small swarmers: down-regulated, big swarmers: up-regulated |
| OG0007993 | Swarmer | 4 | XP_008065105.1, polyadenylate-binding protein 1-like [Carlito syrichta] (1E-19) | nucleic acid binding (GO:0003676) ; RNA binding (GO:0003723) | PF10567 - RNA-recognition motif ; PF06075 - Plant protein of unknown function (DUF936) | rna binding protein, polyadenylate-binding protein, cytoplasmic and nuclear (98, 100) | Post-transcriptional modifications | Small swarmers: down-regulated, big swarmers: up-regulated |
| OG0007950 | Swarmer | 4 | CAJ1328202.1, unnamed protein product [Effrenium voratum] (1E-14) | Acyltransferase 3 domain (IPR002656) | PF01757 - Acyltransferase family | transferase (88, 98.7) | Lipid metabolism | Small swarmers: down-regulated, big swarmers: up-regulated |
| OG0001693 | Swarmer | 10 | - | - | - | oxidoreductase, ribonucleoside-diphosphate reductase, beta subunit 1 (45, 22.3) | Unknown | Small swarmers: down-regulated, big swarmers: up-regulated |
| OG0000206 | Swarmer | 32 | CAK0863910.1, unnamed protein product [Polarella glacialis] (5E-08) | - | - | transcription, neurogenic locus notch homolog protein 3 (83, 99) | Extracellular signalling | Small swarmers: \|log2FC\| < 1-fold, big swarmers: down-regulated |
| OG0000146 | Swarmer | 38 | CAB9506314.1, expressed unknown protein [Seminavis robusta] (7E-31) | - | - | lyase, phosphomethylpyrimidine synthase (45, 45.3) | Unknown | Small swarmers: down-regulated, big swarmers: up-regulated |
| OG0002041 | Swarmer | 9 | - | B30.2/SPRY domain superfamily (IPR043136) ; SPRY domain (IPR003877) ; Concanavalin A-like lectin/glucanase domain superfamily (IPR013320) | PF07177 - Neuralized ; PF00622 - SPRY domain | ligase (59, 42.7) | Protein-protein interaction | Small swarmers: up-regulated, big swarmers: down-regulated |
| OG0004254 | Swarmer | 6 | XP_044558944.1, uncharacterized protein FDP41_006841 [Naegleria fowleri] (8E-80) | Tubulin, C-terminal (IPR023123) ; Tubulin/FtsZ, GTPase domain superfamily (IPR036525) ; Tubulin (IPR000217) ; Tubulin/FtsZ, C-terminal (IPR008280) ; Tubulin/FtsZ, 2-layer sandwich domain (IPR018316) ; Tubulin/FtsZ-like, C-terminal domain (IPR037103) | PF14881 - Tubulin domain ; PF03953 - Tubulin C-terminal domain | structural protein, tubulin alpha-1a chain (97, 100) | Structural cell modifications | Small swarmers: up-regulated, big swarmers: down-regulated |
| OG0007988 | Swarmer | 4 | - | - | - | oxidoreductase, lyase (22, 64.1) | Unknown | Small swarmers: up-regulated, big swarmers: down-regulated |
| OG0008007 | Swarmer | 4 | - | - | - | ValRS/IleRS/LeuRS editing domain (54, 36.9) | Unknown | Small swarmers: up-regulated, big swarmers: down-regulated |
| OG0011473 | Swarmer | 3 | - | - | - | transcription, integrator complex subunit 5 (35, 6.2) | Unknown | Small swarmers: up-regulated, big swarmers: down-regulated |
| OG0018738 | Swarmer big | 2 | - | - | **PF05806 - Noggin glycoprotein, development** in vertebrate body tissues | Ribosome (61, 8.5) | **Cellular development/ growth** | up-regulated |
| OG0018740 | Swarmer big | 2 | - | - | **PF14228 - Cell morphogenesis central region** | Transcription (17, 8.7) | **Cellular development/ growth** | up-regulated |
| OG0018922 | Swarmer big | 2 | CAE8618588.1, unnamed protein product [Polarella glacialis] (8E-46) | Endoplasmic reticulum vesicle transporter (IPR045888) ; Thioredoxin-like superfamily (IPR036249) ; Endoplasmic reticulum vesicle transporter, C-terminal (IPR012936) ; Thioredoxin domain (IPR013766) ; Endoplasmic reticulum vesicle transporter, N-terminal (IPR039542) | PF07970 - Endoplasmic reticulum vesicle transporter ; PF07912 - ERp29, N-terminal domain ; PF13850 - Endoplasmic Reticulum-Golgi Intermediate Compartment (ERGIC) | protein transport (48, 100) | Cell secretion | up-regulated |
| OG0018791 | Swarmer big | 2 | - | - | - | de novo protein (29, 3.7) | Unknown | up-regulated |
| OG0011613 | Swarmer big | 3 | - | - | - | immune system, interleukin enhancer-binding factor 3 (45, 8.5) | Unknown | up-regulated |
| OG0011589 | Swarmer big | 3 | - | Coil | PF08583 - Cytochrome c oxidase biogenesis protein Cmc1 like | transport protein/dna binding protein (40, 6.4) | Unknown | up-regulated |
| OG0004292 | Swarmer big | 6 | - | - | - | dna-binding protein/dna (60, 3.5) | Unknown | up-regulated |
| OG0000384 | Swarmer big | 23 | KHO45317.1, hypothetical protein QS98_C0011G0012 [archaeon GW2011_AR3] (4E-10) | - | - | rna binding protein/rna/dna (36, 7.1) | Unknown | up-regulated |
| OG0019047 | Swarmer big | 2 | - | - | - | transferase (56, 13.8) | Unknown | down-regulated |
| OG0018936 | Swarmer big | 2 | SPQ95657.1, unnamed protein product (mitochondrion) [Plasmodiophora brassicae] (2E-124) | Nucleotide-binding alpha-beta plait domain superfamily (IPR012677) ; RNA recognition motif domain (IPR000504) ; PABP, RNA recognition motif 2 (IPR045305) ; RNA-binding domain superfamily (IPR035979) ; RNA recognition motif domain, eukaryote (IPR003954) | - | rna binding protein (100, 84) | Post-transcriptional modifications | down-regulated |
| OG0018829 | Swarmer big | 2 | HEX3134612.1, TPA: nucleoside monophosphate kinase [Planctomycetota bacterium] (5E-25) | P-loop containing nucleoside triphosphate hydrolase (IPR027417) | PF13238 - AAA domain ; PF01331 - mRNA capping enzyme, catalytic domain, PF01583 - Adenylylsulphate kinase | transferase, adenylate kinase (99.6, 25) | Signalling | down-regulated |
| OG0018783 | Swarmer big | 2 | CAK0906332.1, unnamed protein product [Polarella glacialis] (8E-16) | - | PF01826 - Trypsin Inhibitor like cysteine rich domain | translation (27, 12.7) | Signalling | down-regulated |
| OG0018757 | Swarmer big | 2 | - | - | - | structural genomics, unknown function (54, 5.1) | Unknown | down-regulated |
| OG0018746 | Swarmer big | 2 | - | - | - | cell adhesion (15, 12.3) | Unknown | down-regulated |
| OG0018742 | Swarmer big | 2 | - | - | **PF08317 - Spc7 kinetochore protein** | toxin (42, 31) | **Cell division** | down-regulated |
| OG0011577 | Swarmer big | 3 | - | - | PF11259 - Protein of unknown function (DUF3060) | photosynthesis (6, 12) | Unknown | down-regulated |
| OG0003283 | Cyst | 6 | - | Coil | PF12825 - Domain of unknown function in PX-proteins (DUF3818) | **Contractile protein**, 10s myosin ii (49, 77) | **Cell shape modification** | Cyst (A2ViCY): up-regulated |
| OG0004368 | Cyst | 5 | - | Coil | PF01157 - Ribosomal protein L21e | Structural protein, 'wd40 domain protein' (95, 46) | Protein-protein or proten-DNA interaction | Cyst (A2ViCY): up-regulated |
| OG0005819 | Cyst | 4 | - | - | - | Structural protein, 'tubulin beta-4b chain', 96nm repeat of human respiratory doublet microtubule and associated axonemal complexes (15, 43.5) | Unknown | Cyst (A2ViCY): up-regulated |
| OG00011896 | Cyst | 2 | - | - | - | Structural genomics, unknown function (12.8, 57) | Unknown | Cyst (A2ViCY): down-regulated |
| OG00011831 | Cyst | 2 | - | - | PF15686 - Lysine-rich CEACAM1 co-isolated protein family | Viral protein, uncharacterized protein orf47 (23, 17.2) | Unknown | Cyst (A2ViCY): down-regulated |
| OG00011812 | Cyst | 2 | - | - | - | Chaperone, complex of hsp90 and p50 (61, 13.8) | Unknown | Cyst (A2ViCY): down-regulated |
| OG00011783 | Cyst | 2 | - | - | PF08885 - GSCFA family | Oxidoreductase, lysine-specific demethylase 3b (44, 11.5) | Unknown | Cyst (A2ViCY): down-regulated |
| OG00011782 | Cyst | 2 | - | - | - | Hydrolase, saved domain-containing protein, structure of the calpl/t10 complex (46, 20.5) | Unknown | Cyst (A2ViCY): down-regulated |
| OG0003246 | Swarmer - Cyst | 6 | - | - | PF07541 - Eukaryotic translation initiation factor 2 alpha subunit (EIF_2_alpha) ; PF06827 - zing finger found in FPG and IleRS ; PF18551 - TackOD1 (Thaumarchaeal output domain 1) is a predicted metal-binding domain ; PF04810 - Sec23/Sec24 zinc finger ; **PF08457 - Sfi1 spindle body protein** | Ribosome (60, 31) | **Cell division related** | Cyst (A2ViCY): up-regulated, small swarmers: up-regulated, big swarmers: up-regulated |
| OG0004364 | Swarmer - Cyst | 5 | - | - | PF05064 - nucleoprotein Nsp1-like C-terminal region ; **PF05483 - Synaptonemal complex protein 1 (SCP-1) ;** PF12761 - Actin cytoskeleton regulatory complex protein END3 ; **PF17078 - SWI5 (homologous recombination repair protein)-dependent HO expression protein 3** ; PF13747 - Domain of unknown function (DUF4164) | Cell adhesion, surface-exposed lipoprotein (44, 22.4) | **Meiosis related** | Cyst (A2ViCY): \|log2FC\| < 1-fold, small swarmers (A2ViSW): up-regulated, big swarmers: down-regulated |
| OG0001269 | Swarmer - Cyst | 11 | XP_039262999.1, radial spoke head protein 4 homolog A-like [Styela clava] (2E-130) | Radial spokehead-like protein (IPR006802) | PF04712 - Radial spokehead-like protein | Structural protein, radial spoke head protein 4 homolog a (76, 100) | **Flagella associated** | Big swarmers: down-regulated, other life stages: \|log2FC\| < 1-fold |
| OG0000766 | Swarmer - Cyst | 15 | - | - | - | De novo protein (13.9, 32) | Unknown | Cyst (A2ViCY): up-regulated, small swarmers (A2ViSW): down-regulated, big swarmers: up-regulated |
| OG0000641 | Swarmer - Cyst | 17 | XP_042335297.1, heat shock factor protein 3-like isoform X1 [Sceloporus undulatus] (1E-08) | Heat shock transcription factor family (IPR027725) | PF00447 - HSF-type DNA-binding | - | Heat shock protein | Cyst (A2ViCY): up-regulated, small swarmers: \|log2FC\| < 1-fold, big swarmers: up-regulated |
| OG0001260 | Swarmer - Cyst | 11 | XP_039599047.1, calmodulin-like [Polypterus senegalus] (3E-21) | EF-Hand 1, calcium-binding site (IPR018247) ; EF-hand domain (IPR002048) ; EF-hand domain pair (IPR011992) | PF13499 - EF-hand domain pair ; PF14513 - Diacylglycerol kinase N-terminus | Hydrolase (100, 99.9) | Signalling | Cyst (A2ViCY): down-regulated, small swarmers: \|log2FC\| < 1-fold, big swarmers: up-regulated |
| OG0001854 | Swarmer big - Vegetative | 9 | HXJ95225.1, TPA: SUMF1/EgtB/PvdO family nonheme iron enzyme [Terriglobia bacterium] (4E-148) | Sulfatase-modifying factor enzyme (IPR005532) ; C-type lectin fold (IPR016187) ; Sulfatase-modifying factor enzyme superfamily (IPR042095) | PF14295 - PAN domain ; PF03781 - Sulfatase-modifying factor enzyme 1 ; PF10708 - Protein of unknown function (DUF2510) | C-type lectin-like, Sulfatase-modifying factor-like (89, 100) | Protein modification | down-regulated |
| OG0001900 | Swarmer big - Vegetative | 9 | XP_023322804.1\| pleckstrin homology-like domain family B member 1 [Eurytemora carolleeae] (1E-79) | Pleckstrin homology domain IPR001849, PH-like domain superfamily IPR011993 | PF00169 - PH domain, pleckstrin homology, wide range of proteins involved in intracellular signaling or as constituents of the cytoskeleton ; PF15413 - Pleckstrin homology domain ; PF15409 - Pleckstrin homology domain | Lipid binding protein, cytohesin-3, adp-ribosylation factor 6 (97, 99.9) | Signalling | down-regulated |
| OG0003624 | Swarmer big - Vegetative | 6 | CAH0372151.1, unnamed protein product [Pelagomonas calceolata] (3E-27) | PH-like domain superfamily (IPR011993) ; Pleckstrin homology domain (IPR001849) | PF10503 - Esterase PHB depolymerase ; PF00756 - Putative esterase | Hydrolase, est1e from butyrivibrio proteoclasticus (65, 99) | Catabolism | down-regulated |
| OG0004095 | Swarmer big - Vegetative | 6 | KAJ0398490.1, hypothetical protein P43SY_006648 [Pythium insidiosum] (9E-19) | WD40-repeat-containing domain superfamily (IPR036322) ; WD40/YVTN repeat-like-containing domain superfamily (IPR015943) ; WD40 repeat (IPR001680) ; Quinoprotein alcohol dehydrogenase-like superfamily (IPR011047) | PF02239 - Cytochrome D1 heme domain ; PF11715 - Nucleoporin Nup120/160 ; PF15492 - Neuroblastoma-amplified sequence, N terminal ; PF08662 - Eukaryotic translation initiation factor eIF2A ; PF16529 - WD40 region of Ge1, enhancer of mRNA-decapping protein | Motor protein, wd repeat-containing protein 34 (96, 99.4) | Protein-protein interaction | down-regulated |
| OG0000314 | Swarmer - Vegetative | 25 | MBL6833635.1, RING-HC finger protein [Candidatus Actinomarina sp.] (1E-09) | RING finger protein 141-like (IPR047126) | PF13920 - Zinc finger, C3HC4 type (RING finger) ; PF13857, PF12796 - Ankyrin repeats ; PF09976 - Tetratricopeptide repeat-like domain ; PF05565 - Siphovirus Gp157 ; PF16559 - GIT coiled-coil Rho guanine nucleotide exchange factor ; **PF05141** - Pyoverdine/dityrosine biosynthesis protein, DIT1 is involved in synthesising dityrosine, a **sporulation-specific component** of the yeast ascospore wall that is essential for the resistance of the spores to adverse environmental conditions ; PF13445 - RING-type zinc-finger | DNA binding protein, DNA repair protein RAD5 (78, 96.1) | **Reproduction related** | Vegetative cells: down-regulated, small swarmers (A2ViSW): up-regulated, big swarmers: up-regulated |
| OG0000544 | Swarmer - Vegetative | 19 | XP_007515318.1, predicted protein [Bathycoccus prasinos] (3E-95) | Small ribosomal subunit protein eS10 (IPR037447) | PF03501 - Plectin/S10 domain | ribosome, 40s ribosomal protein s10 (87, 100) | Ribosome | Vegetative cells: up-regulated, small swarmers (A1ViSW): down-regulated, big swarmers: down-regulated |
| OG0000894 | Swarmer - Vegetative | 14 | KAK4048103.1, iso-1-cytochrome c [Microbotryomycetes sp. JL201] iso-1-cytochrome c [Microbotryomycetes sp. JL201] (1E-65) | Cytochrome c, class IA/ IB (IPR002327) | PF00034 - Cytochrome c | Cytochrome c (98, 99.9) | Energy production | Vegetative cells: up-regulated, small swarmers (A2ViSW): down-regulated, big swarmers: down-regulated |
| OG0002425 | Swarmer - Vegetative | 8 | ETO35508.1, triosephosphate isomerase [Reticulomyxa filosa] (8E-82) | Triosephosphate isomerase (IPR000652) | PF00121 - Triosephosphate isomerase | Triosephosphate isomerase (TIM) (98, 100) | Energy production | Vegetative cells: up-regulated, small swarmers (A1ViSW): down-regulated, big swarmers: down-regulated |
| OG0000254 | Swarmer - Vegetative | 28 | HLG25242.1, TPA: cold shock domain-containing protein [Candidatus Gracilibacteria bacterium] (1E-12) | Nucleic acid-binding, OB-fold (IPR012340) ; Cold-shock protein, DNA-binding ; (IPR002059) ; Cold shock domain (IPR011129) | PF00313 - Cold-shock' DNA-binding domain | Rna binding protein, upstream of n-ras (75, 99.9) | Nucleic acid binding | Vegetative cells (A6GrVE, A9GrVE): down-regulated, small swarmers: up-regulated, big swarmers: up-regulated |
| OG0000349 | Swarmer - Vegetative | 24 | CAE8715553.1, unnamed protein product [Polarella glacialis] (6E-101) | 3-beta hydroxysteroid dehydrogenase/isomerase (IPR002225) ; NAD(P)-binding domain superfamily (IPR036291) | PF05368 - NmrA-like family ; PF01073 - 3-beta hydroxysteroid dehydrogenase/isomerase family ; PF01370 - NAD dependent epimerase/dehydratase family | Oxidoreductase (89, 99.8) | Catabolism | Vegetative cells: up-regulated, small swarmers: \|log2FC\| < 1-fold, big swarmers: down-regulated |
| OG0000903 | Swarmer - Vegetative | 14 | KAJ6635732.1, Chromobox protein like 3 [Pseudolycoriella hygida] (7E-44) | Chromo/chromo shadow domain (IPR000953) | PF00847 - AP2 domain ; PF00385 - Chromo (CHRromatin Organisation MOdifier) domain ; PF00397 - WW domain ; PF00628 - PHD-finger ; PF01393 - Chromo shadow domain ; PF12796 - Ankyrin repeats (3 copies) | Ligase, e3 ubiquitin-protein ligase itchy (47, 99.9) | Chromatin structure | Vegetative cells: down-regulated, small swarmers: \|log2FC\| < 1-fold, big swarmers: \|log2FC\| < 1-fold |
| OG0001395 | Swarmer - Vegetative | 11 | ALD47951.1, adaptor protein complex 1 subunit sigma [Isochrysis galbana] (5E-99) | Adaptor protein complex, sigma subunit (IPR016635) | PF01217 - Clathrin adaptor complex small chain | Profilin-like, SNARE-like, Clathrin coat assembly domain (99, 100) | Signalling | Vegetative cells: down-regulated, small swarmers: \|log2FC\| < 1-fold, big swarmers: \|log2FC\| < 1-fold |
| OG0001390 | Swarmer - Vegetative | 11 | NWG23444.1, sulfotransferase domain-containing protein [Pseudorhodoplanes sp.] (2E-15) | Heparan sulfate sulfotransferase (IPR037359) ; Sulfotransferase domain (IPR000863) ; P-loop containing nucleoside triphosphate hydrolase (IPR027417) | PF00685 - Sulfotransferase domain ; PF13469 - Sulfotransferase family | Transferase (83, 99.8) | Catabolism | Vegetative cells: down-regulated, small swarmers: \|log2FC\| < 1-fold, big swarmers: down-regulated |
| OG0002281 | Swarmer - Vegetative | 8 | MAA70164.1, 50S ribosomal protein L13 [Bermanella sp.] (7E-99) | Large ribosomal subunit protein uL13 (IPR005822) | PF00572 - Ribosomal protein L13 ; PF00380 - Ribosomal protein S9/S16 | Ribosomal protein L13 (99, 100) | Ribosome | Vegetative cells: down-regulated, small swarmers: down-regulated, big swarmers: down-regulated |
| OG0000969 | Cyst - Vegetative | 13 | CEO99023.1, hypothetical protein PBRA_007137 [Plasmodiophora brassicae] (4E-62) | cAMP-dependent protein kinase regulatory subunit (IPR012198) | PF00027 - Cyclic nucleotide-binding domain ; PF00536 - SAM domain (Sterile alpha motif) ; PF04831 - Popeye protein conserved region ; PF13095 - Kinetochore Sim4 complex subunit FTA2 | Transferase, cgmp-dependent protein kinase, putative (77, 100) | Signalling | down-regulated |
| OG0007516 | Vegetative | 4 | ETO30159.1, hypothetical protein RFI_06961 [Reticulomyxa filosa] (2E-25) | - | - | lyase, 2-c-methyl-d-erythritol 2,4-cyclodiphosphate synthase (33, 15) | Unknown | up-regulated |

**Table S4. List of lineage-specific HMM profiles that were manually created for this study.** The last two columns (i.e., kmer length and cut-ga) are parameters relative to a homemade R script (cf. Data availability). In this script, the kmer length was manually selected according to the number of available sequences for the created of each HMM profile and, then, the internal threshold of the program hmmsearch (i.e., cut-ga) was manually calculated.

| Gene ID | Domain ID | Domain/gene name | PFAM/InterPro ID | Function | Cellular process | Molecular process | Specificity | Lineage | Taxonomy accession (Uniprot) | Link | Nb seq | kmer length for cut-ga | Manual cut ga (max domain score) |
|---|---|---|---|---|---|---|---|---|---|---|---|---|---|
| **GEX1** | - | Protein GAMETE EXPRESSED 1 | **IPR040346** | Required for karyogamy in flowering plants | Syngamy | Karyogamy | Gamete specific | SAR | 2698737 | https://www.ebi.ac.uk/interpro/entry/InterPro/IPR040346/protein/UniProt/taxonomy/uniprot/2698737/#table | 135 | 4 | 23.3 |
| **GEX1** | - | Protein GAMETE EXPRESSED 1 | **IPR040346** | Required for karyogamy in flowering plants | Syngamy | Karyogamy | Gamete specific | Streptophyta | 35493 | https://www.ebi.ac.uk/interpro/entry/InterPro/IPR040346/protein/UniProt/taxonomy/uniprot/35493/#table | 617 | 3 | 31.4 |
| **HAP2/GCS1** | **HAP2/GCS1** | Hapless 2 / Generative Cell Specific-1 | **PF10699** | Considered ancestral gamete fusogan, required for membrane fusion in protists, flowering plants and some invertebrates | Syngamy | Plasmogamy | Gamete specific | Alveolata | 33630 | https://www.ebi.ac.uk/interpro/entry/pfam/PF10699/protein/UniProt/taxonomy/uniprot/33630/#table | 177 | 4 | 20 |
| **HAP2/GCS1** | **HAP2/GCS1** | Hapless 2 / Generative Cell Specific-1 | **PF10699** | Considered ancestral gamete fusogan, required for membrane fusion in protists, flowering plants and some invertebrates | Syngamy | Plasmogamy | Gamete specific | Rhizaria (Retaria) | 46433 | https://www.ebi.ac.uk/interpro/entry/pfam/PF10699/protein/UniProt/taxonomy/uniprot/46433/#table | 2 | 4 | 35 |
| **HAP2/GCS1** | **HAP2/GCS1** | Hapless 2 / Generative Cell Specific-1 | **PF10699** | Considered ancestral gamete fusogan, required for membrane fusion in protists, flowering plants and some invertebrates | Syngamy | Plasmogamy | Gamete specific | Stramenopiles | 33634 | https://www.ebi.ac.uk/interpro/entry/pfam/PF10699/protein/UniProt/taxonomy/uniprot/33634/#table | 6 | 4 | 23.8 |
| **HAP2/GCS1** | **HAP2/GCS1** | Hapless 2 / Generative Cell Specific-1 | **PF10699** | Considered ancestral gamete fusogan, required for membrane fusion in protists, flowering plants and some invertebrates | Syngamy | Plasmogamy | Gamete specific | Chloroplastida (Chlorophyceae) | 3166 | https://www.ebi.ac.uk/interpro/entry/pfam/PF10699/protein/UniProt/taxonomy/uniprot/3166/#table | 25 | 4 | 20.2 |
| **HAP2/GCS1** | **HAP2/GCS1** | Hapless 2 / Generative Cell Specific-1 | **PF10699** | Considered ancestral gamete fusogan, required for membrane fusion in protists, flowering plants and some invertebrates | Syngamy | Plasmogamy | Gamete specific | Metazoa | 33208 | https://www.ebi.ac.uk/interpro/entry/pfam/PF10699/protein/UniProt/taxonomy/uniprot/33208/#table | 198 | 4 | 28 |
| **HAP2/GCS1** | **HAP2/GCS1** | Hapless 2 / Generative Cell Specific-1 | **PF10699** | Considered ancestral gamete fusogan, required for membrane fusion in protists, flowering plants and some invertebrates | Syngamy | Plasmogamy | Gamete specific | Amoebozoa | 554915 | https://www.ebi.ac.uk/interpro/entry/pfam/PF10699/protein/UniProt/taxonomy/uniprot/554915/#table | 16 | 5 | 11.9 |
| **CFA20** | **CFA20** | Cilia- and flagella-associated protein 20 | **PF05018** | Cilia and flagella associated protein | Syngamy | Flagellar protein | Cilia/flagella | Alveolata | 33630 | https://www.ebi.ac.uk/interpro/entry/pfam/PF05018/protein/UniProt/taxonomy/uniprot/33630/#table | 299 | 3 | 37.1 |
| **CFA20** | **CFA20** | Cilia- and flagella-associated protein 20 | **PF05018** | Cilia and flagella associated protein | Syngamy | Flagellar protein | Cilia/flagella | Rhizaria | 543769 | https://www.ebi.ac.uk/interpro/entry/pfam/PF05018/protein/UniProt/taxonomy/uniprot/543769/#table | 15 | 3 | 12.7 |
| **CFA20** | **CFA20** | Cilia- and flagella-associated protein 20 | **PF05018** | Cilia and flagella associated protein | Syngamy | Flagellar protein | Cilia/flagella | Stramenopiles | 33634 | https://www.ebi.ac.uk/interpro/entry/pfam/PF05018/protein/UniProt/taxonomy/uniprot/33634/#table | 295 | 3 | 16.3 |
| **CFA20** | **CFA20** | Cilia- and flagella-associated protein 20 | **PF05018** | Cilia and flagella associated protein | Syngamy | Flagellar protein | Cilia/flagella | Chloroplastida (Clorophyta) | 3041 | https://www.ebi.ac.uk/interpro/entry/pfam/PF05018/protein/UniProt/taxonomy/uniprot/3041/#table | 112 | 3 | 14 |
| **CFA20** | **CFA20** | Cilia- and flagella-associated protein 20 | **PF05018** | Cilia and flagella associated protein | Syngamy | Flagellar protein | Cilia/flagella | Metazoa (Choanoflagellata) | 28009 | https://www.ebi.ac.uk/interpro/entry/pfam/PF05018/protein/UniProt/taxonomy/uniprot/28009/#table | 3 | 4 | 20.8 |
| **CFA20** | **CFA20** | Cilia- and flagella-associated protein 20 | **PF05018** | Cilia and flagella associated protein | Syngamy | Flagellar protein | Cilia/flagella | Amoebozoa | 554915 | https://www.ebi.ac.uk/interpro/entry/pfam/PF05018/protein/UniProt/taxonomy/uniprot/554915/#table | 7 | 4 | 20 |
| **REC8** | RAD21_REC8 | Recombination protein 8 | **PF04824** | Sister chromatid cohesion | Meiosis | Cohesion complex | Meiosis specific | Alveolata | 33630 | https://www.ebi.ac.uk/interpro/entry/pfam/PF04824/protein/UniProt/taxonomy/uniprot/33630/#table | 58 | 5 | 19.7 |
| **REC8** | RAD21_REC8 | Recombination protein 8 | **PF04824** | Sister chromatid cohesion | Meiosis | Cohesion complex | Meiosis specific | Rhizaria | 543769 | https://www.ebi.ac.uk/interpro/entry/pfam/PF04824/protein/UniProt/taxonomy/uniprot/543769/#table | 6 | 5 | 28.2 |
| **REC8** | RAD21_REC8 | Recombination protein 8 | **PF04824** | Sister chromatid cohesion | Meiosis | Cohesion complex | Meiosis specific | Stramenopiles | 33634 | https://www.ebi.ac.uk/interpro/entry/pfam/PF04824/protein/UniProt/taxonomy/uniprot/33634/#table | 101 | 5 | 38.7 |
| **REC8** | RAD21_REC8 | Recombination protein 8 | **PF04824** | Sister chromatid cohesion | Meiosis | Cohesion complex | Meiosis specific | Chloroplastida (Clorophyta) | 3041 | https://www.ebi.ac.uk/interpro/entry/pfam/PF04824/protein/UniProt/taxonomy/uniprot/3041/#table | 86 | 4 | 26.9 |
| **REC8** | RAD21_REC8 | Recombination protein 8 | **PF04824** | Sister chromatid cohesion | Meiosis | Cohesion complex | Meiosis specific | Metazoa (Choanoflagellata) | 28009 | https://www.ebi.ac.uk/interpro/entry/pfam/PF04824/protein/UniProt/taxonomy/uniprot/28009/#table | 2 | 5 | 27.6 |
| **REC8** | RAD21_REC8 | Recombination protein 8 | **PF04824** | Sister chromatid cohesion | Meiosis | Cohesion complex | Meiosis specific | Amoebozoa | 554915 | https://www.ebi.ac.uk/interpro/entry/pfam/PF04824/protein/UniProt/taxonomy/uniprot/554915/#table | 17 | 5 | 23.3 |
| **REC8_N** | RAD21_REC8_N | Recombination protein 8 | **PF04825** | Sister chromatid cohesion | Meiosis | Cohesion complex | Meiosis specific | Alveolata | 33630 | https://www.ebi.ac.uk/interpro/entry/pfam/PF04825/protein/UniProt/taxonomy/uniprot/33630/#table | 146 | 5 | 22.7 |
| **REC8_N** | RAD21_REC8_N | Recombination protein 8 | **PF04825** | Sister chromatid cohesion | Meiosis | Cohesion complex | Meiosis specific | Rhizaria | 543769 | https://www.ebi.ac.uk/interpro/entry/pfam/PF04825/protein/UniProt/taxonomy/uniprot/543769/#table | 5 | 5 | 26.6 |
| **REC8_N** | RAD21_REC8_N | Recombination protein 8 | **PF04825** | Sister chromatid cohesion | Meiosis | Cohesion complex | Meiosis specific | Stramenopiles | 33634 | https://www.ebi.ac.uk/interpro/entry/pfam/PF04825/protein/UniProt/taxonomy/uniprot/33634/#table | 104 | 4 | 44.2 |
| **REC8_N** | RAD21_REC8_N | Recombination protein 8 | **PF04825** | Sister chromatid cohesion | Meiosis | Cohesion complex | Meiosis specific | Chloroplastida (Clorophyta) | 3041 | https://www.ebi.ac.uk/interpro/entry/pfam/PF04825/protein/UniProt/taxonomy/uniprot/3041/#table | 114 | 4 | 16.6 |
| **REC8_N** | RAD21_REC8_N | Recombination protein 8 | **PF04825** | Sister chromatid cohesion | Meiosis | Cohesion complex | Meiosis specific | Metazoa (Choanoflagellata) | 28009 | https://www.ebi.ac.uk/interpro/entry/pfam/PF04825/protein/UniProt/taxonomy/uniprot/28009/#table | 3 | 5 | 17.4 |
| **REC8_N** | RAD21_REC8_N | Recombination protein 8 | **PF04825** | Sister chromatid cohesion | Meiosis | Cohesion complex | Meiosis specific | Amoebozoa | 554915 | https://www.ebi.ac.uk/interpro/entry/pfam/PF04825/protein/UniProt/taxonomy/uniprot/554915/#table | 18 | 5 | 33.4 |
| **HOP1** | HORMA | Meiosis-specific protein HOP1 | PF02301 | Synaptonemal complex (SC) protein involved in chromosome pairing during meiosis | Meiosis | Homologous alignment and synaptonemal complex | Meiosis specific | Alveolata | 33630 | https://www.ebi.ac.uk/interpro/entry/pfam/PF02301/protein/UniProt/taxonomy/uniprot/33630/#table | 130 | 4 | 20.8 |
| **HOP1** | HORMA | Meiosis-specific protein HOP1 | PF02301 | Synaptonemal complex (SC) protein involved in chromosome pairing during meiosis | Meiosis | Homologous alignment and synaptonemal complex | Meiosis specific | Rhizaria | 543769 | https://www.ebi.ac.uk/interpro/entry/pfam/PF02301/protein/UniProt/taxonomy/uniprot/543769/#table | 11 | 5 | 20 |
| **HOP1** | HORMA | Meiosis-specific protein HOP1 | PF02301 | Synaptonemal complex (SC) protein involved in chromosome pairing during meiosis | Meiosis | Homologous alignment and synaptonemal complex | Meiosis specific | Stramenopiles | 33634 | https://www.ebi.ac.uk/interpro/entry/pfam/PF02301/protein/UniProt/taxonomy/uniprot/33634/#table | 166 | 4 | 32.3 |
| **HOP1** | HORMA | Meiosis-specific protein HOP1 | PF02301 | Synaptonemal complex (SC) protein involved in chromosome pairing during meiosis | Meiosis | Homologous alignment and synaptonemal complex | Meiosis specific | Chloroplastida (Clorophyta) | 3041 | https://www.ebi.ac.uk/interpro/entry/pfam/PF02301/protein/UniProt/taxonomy/uniprot/3041/#table | 104 | 4 | 29.2 |
| **HOP1** | HORMA | Meiosis-specific protein HOP1 | PF02301 | Synaptonemal complex (SC) protein involved in chromosome pairing during meiosis | Meiosis | Homologous alignment and synaptonemal complex | Meiosis specific | Metazoa (Choanoflagellata) | 28009 | https://www.ebi.ac.uk/interpro/entry/pfam/PF02301/protein/UniProt/taxonomy/uniprot/28009/#table | 5 | 5 | 19.6 |
| **HOP1** | HORMA | Meiosis-specific protein HOP1 | PF02301 | Synaptonemal complex (SC) protein involved in chromosome pairing during meiosis | Meiosis | Homologous alignment and synaptonemal complex | Meiosis specific | Amoebozoa | 554915 | https://www.ebi.ac.uk/interpro/entry/pfam/PF02301/protein/UniProt/taxonomy/uniprot/554915/#table | 18 | 5 | 24.8 |
| **SPO22** | **SPO22/ZIP4** | Sporulation-specific protein 22 | PF08631 | SPO22/ZIP4 in yeast is a meiosis specific protein involved in sporulation. It has been shown to regulate crossover distribution by promoting synaptonemal complex formation | Meiosis | Homologous alignment and synaptonemal complex | Meiosis/mitosis | Rhizaria/ Stramenopiles | 543769, 33634 | https://www.ebi.ac.uk/interpro/entry/pfam/PF02301/taxonomy/uniprot/#tree | 3 | 6 | 20.3 |
| **SPO22** | **SPO22/ZIP4** | Sporulation-specific protein 22 | PF08631 | SPO22/ZIP4 in yeast is a meiosis specific protein involved in sporulation. It has been shown to regulate crossover distribution by promoting synaptonemal complex formation | Meiosis | Homologous alignment and synaptonemal complex | Meiosis/mitosis | Chloroplastida (Clorophyta) | 3041 | https://www.ebi.ac.uk/interpro/entry/pfam/PF08631/protein/UniProt/taxonomy/uniprot/3041/#table | 5 | 5 | 26 |
| **SPO22** | **SPO22/ZIP4** | Sporulation-specific protein 22 | PF08631 | SPO22/ZIP4 in yeast is a meiosis specific protein involved in sporulation. It has been shown to regulate crossover distribution by promoting synaptonemal complex formation | Meiosis | Homologous alignment and synaptonemal complex | Meiosis/mitosis | Amoebozoa | 554915 | https://www.ebi.ac.uk/interpro/entry/pfam/PF08631/protein/UniProt/taxonomy/uniprot/554915/#table | 2 | 5 | 24.5 |
| **SPO11** | TOP6A-Spo11_Toprim | Meiosis-specific protein SPO11 | PF21180 | Creation of double strand breaks to initiation meiotic recombination | Meiosis | DSB | Meiosis specific | Alveolata | 33630 | https://www.ebi.ac.uk/interpro/entry/pfam/PF21180/protein/UniProt/taxonomy/uniprot/33630/#table | 226 | 4 | 27.4 |
| **SPO11** | TOP6A-Spo11_Toprim | Meiosis-specific protein SPO11 | PF21180 | Creation of double strand breaks to initiation meiotic recombination | Meiosis | DSB | Meiosis specific | Rhizaria | 543769 | https://www.ebi.ac.uk/interpro/entry/pfam/PF21180/protein/UniProt/taxonomy/uniprot/543769/#table | 4 | 4 | 20.3 |
| **SPO11** | TOP6A-Spo11_Toprim | Meiosis-specific protein SPO11 | PF21180 | Creation of double strand breaks to initiation meiotic recombination | Meiosis | DSB | Meiosis specific | Stramenopiles | 33634 | https://www.ebi.ac.uk/interpro/entry/pfam/PF21180/protein/UniProt/taxonomy/uniprot/33634/#table | 131 | 4 | 29.3 |
| **SPO11** | TOP6A-Spo11_Toprim | Meiosis-specific protein SPO11 | PF21180 | Creation of double strand breaks to initiation meiotic recombination | Meiosis | DSB | Meiosis specific | Chloroplastida (Clorophyta) | 3041 | https://www.ebi.ac.uk/interpro/entry/pfam/PF21180/protein/UniProt/taxonomy/uniprot/3041/#table | 96 | 4 | 26.6 |
| **SPO11** | TOP6A-Spo11_Toprim | Meiosis-specific protein SPO11 | PF21180 | Creation of double strand breaks to initiation meiotic recombination | Meiosis | DSB | Meiosis specific | Metazoa (Choanoflagellata) | 28009 | https://www.ebi.ac.uk/interpro/entry/pfam/PF21180/protein/UniProt/taxonomy/uniprot/28009/#table | 2 | 4 | 20.3 |
| **SPO11** | TOP6A-Spo11_Toprim | Meiosis-specific protein SPO11 | PF21180 | Creation of double strand breaks to initiation meiotic recombination | Meiosis | DSB | Meiosis specific | Amoebozoa | 554915 | https://www.ebi.ac.uk/interpro/entry/pfam/PF21180/protein/UniProt/taxonomy/uniprot/554915/#table | 26 | 4 | 24.4 |
| **HOP2** | TBPIP | Homologous-pairing protein 2 | PF07106 | Forms heterodimer with MND1 promoting interhomolog meiotic recombination | Meiosis | Recombination | Meiosis specific | Alveolata | 33630 | https://www.ebi.ac.uk/interpro/entry/pfam/PF07106/protein/UniProt/taxonomy/uniprot/33630/#table | 141 | 4 | 32.5 |
| **HOP2** | TBPIP | Homologous-pairing protein 2 | PF07106 | Forms heterodimer with MND1 promoting interhomolog meiotic recombination | Meiosis | Recombination | Meiosis specific | Stramenopiles | 33634 | https://www.ebi.ac.uk/interpro/entry/pfam/PF07106/protein/UniProt/taxonomy/uniprot/33634/#table | 77 | 4 | 31.2 |
| **HOP2** | TBPIP | Homologous-pairing protein 2 | PF07106 | Forms heterodimer with MND1 promoting interhomolog meiotic recombination | Meiosis | Recombination | Meiosis specific | Amoebozoa | 554915 | https://www.ebi.ac.uk/interpro/entry/pfam/PF07106/protein/UniProt/taxonomy/uniprot/554915/#table | 18 | 4 | 27.9 |
| **MND1** | HTH | Meiotic nuclear division protein 1 | PF03962 | Forms heterodimer with HOP2 promoting interhomolog meiotic recombination | Meiosis | Recombination | Meiosis specific | Alveolata | 33630 | https://www.ebi.ac.uk/interpro/entry/pfam/PF03962/taxonomy/uniprot/#tree | 123 | 3 | 22 |
| **MND1** | HTH | Meiotic nuclear division protein 1 | PF03962 | Forms heterodimer with HOP2 promoting interhomolog meiotic recombination | Meiosis | Recombination | Meiosis specific | Rhizaria | 543769 | https://www.ebi.ac.uk/interpro/entry/pfam/PF03962/protein/UniProt/taxonomy/uniprot/543769/#table | 5 | 3 | 25.7 |
| **MND1** | HTH | Meiotic nuclear division protein 1 | PF03962 | Forms heterodimer with HOP2 promoting interhomolog meiotic recombination | Meiosis | Recombination | Meiosis specific | Stramenopiles | 33634 | https://www.ebi.ac.uk/interpro/entry/pfam/PF03962/protein/UniProt/taxonomy/uniprot/33634/#table | 147 | 3 | 29.7 |
| **MND1** | HTH | Meiotic nuclear division protein 1 | PF03962 | Forms heterodimer with HOP2 promoting interhomolog meiotic recombination | Meiosis | Recombination | Meiosis specific | Chloroplastida (Clorophyta) | 3041 | https://www.ebi.ac.uk/interpro/entry/pfam/PF03962/taxonomy/uniprot/#tree | 52 | 3 | 33.5 |
| **MND1** | HTH | Meiotic nuclear division protein 1 | PF03962 | Forms heterodimer with HOP2 promoting interhomolog meiotic recombination | Meiosis | Recombination | Meiosis specific | Amoebozoa | 554915 | https://www.ebi.ac.uk/interpro/entry/pfam/PF03962/protein/UniProt/taxonomy/uniprot/554915/#table | 19 | 3 | 23.2 |
| **MND1** | Leucine zipper with capping helix domain | Meiotic nuclear division protein 1 | PF18517 | Forms heterodimer with HOP2 promoting interhomolog meiotic recombination | Meiosis | Recombination | Meiosis specific | Alveolata | 33630 | https://www.ebi.ac.uk/interpro/entry/pfam/PF18517/protein/UniProt/taxonomy/uniprot/33630/#table | 54 | 3 | 18 |
| **MND1** | Leucine zipper with capping helix domain | Meiotic nuclear division protein 1 | PF18517 | Forms heterodimer with HOP2 promoting interhomolog meiotic recombination | Meiosis | Recombination | Meiosis specific | Rhizaria | 543769 | https://www.ebi.ac.uk/interpro/entry/pfam/PF18517/protein/UniProt/taxonomy/uniprot/543769/#table | 3 | 4 | 18.3 |
| **MND1** | Leucine zipper with capping helix domain | Meiotic nuclear division protein 1 | PF18517 | Forms heterodimer with HOP2 promoting interhomolog meiotic recombination | Meiosis | Recombination | Meiosis specific | Stramenopiles | 33634 | https://www.ebi.ac.uk/interpro/entry/pfam/PF18517/protein/UniProt/taxonomy/uniprot/33634/#table | 119 | 3 | 27.2 |
| **MND1** | Leucine zipper with capping helix domain | Meiotic nuclear division protein 1 | PF18517 | Forms heterodimer with HOP2 promoting interhomolog meiotic recombination | Meiosis | Recombination | Meiosis specific | Chloroplastida (Clorophyta) | 3041 | https://www.ebi.ac.uk/interpro/entry/pfam/PF18517/protein/UniProt/taxonomy/uniprot/3041/ | 79 | 3 | 23.2 |
| **MND1** | Leucine zipper with capping helix domain | Meiotic nuclear division protein 1 | PF18517 | Forms heterodimer with HOP2 promoting interhomolog meiotic recombination | Meiosis | Recombination | Meiosis specific | Amoebozoa | 554915 | https://www.ebi.ac.uk/interpro/entry/pfam/PF18517/protein/UniProt/taxonomy/uniprot/554915/#table | 33 | 3 | 19.7 |
| **DMC1** | - | Meiotic recombination protein DMC1 | **IPR011940** | Meiosis-specific homolog of Rad51, has similar function but promotes interhomolog recombination | Meiosis | Recombination | Meiosis specific | Alveolata | 33630 | https://www.ebi.ac.uk/interpro/entry/InterPro/IPR011940/protein/UniProt/taxonomy/uniprot/33630/#table | 108 | 3 | 20.9 |
| **DMC1** | - | Meiotic recombination protein DMC1 | **IPR011940** | Meiosis-specific homolog of Rad51, has similar function but promotes interhomolog recombination | Meiosis | Recombination | Meiosis specific | Rhizaria | 543769 | https://www.ebi.ac.uk/interpro/entry/InterPro/IPR011940/protein/UniProt/taxonomy/uniprot/543769/#table | 2 | 3 | 18.2 |
| **DMC1** | - | Meiotic recombination protein DMC1 | **IPR011940** | Meiosis-specific homolog of Rad51, has similar function but promotes interhomolog recombination | Meiosis | Recombination | Meiosis specific | Stramenopiles | 33634 | https://www.ebi.ac.uk/interpro/entry/InterPro/IPR011940/protein/UniProt/taxonomy/uniprot/33634/#table | 22 | 3 | 22.4 |
| **DMC1** | - | Meiotic recombination protein DMC1 | **IPR011940** | Meiosis-specific homolog of Rad51, has similar function but promotes interhomolog recombination | Meiosis | Recombination | Meiosis specific | Chloroplastida (Clorophyta) | 3041 | https://www.ebi.ac.uk/interpro/entry/InterPro/IPR011940/protein/UniProt/taxonomy/uniprot/3041/#table | 18 | 4 | 26.7 |
| **DMC1** | - | Meiotic recombination protein DMC1 | **IPR011940** | Meiosis-specific homolog of Rad51, has similar function but promotes interhomolog recombination | Meiosis | Recombination | Meiosis specific | Amoebozoa | 554915 | https://www.ebi.ac.uk/interpro/entry/InterPro/IPR011940/protein/UniProt/taxonomy/uniprot/554915/#table | 12 | 3 | 19.7 |
